# Reward timescale controls the rate of behavioral and dopaminergic learning

**DOI:** 10.1101/2023.03.31.535173

**Authors:** Dennis A Burke, Annie Taylor, Huijeong Jeong, SeulAh Lee, Brenda Wu, Joseph R Floeder, Vijay Mohan K Namboodiri

**Affiliations:** Department of Neurology, University of California, San Francisco, CA, USA; Neuroscience Graduate Program, University of California, San Francisco, CA, USA; University of California, Berkeley, CA, USA; Weill Institute for Neurosciences, Kavli Institute for Fundamental Neuroscience, Center for Integrative Neuroscience, University of California, San Francisco, CA, USA

## Abstract

Learning the causes of rewards is necessary for survival. Thus, it is critical to understand the mechanisms of such a vital biological process. Cue-reward learning is controlled by mesolimbic dopamine and improves with spacing of cue-reward pairings. However, whether a mathematical rule governs such improvements in learning rate, and if so, whether a unifying mechanism captures this rule and dopamine dynamics during learning remain unknown. Here, we investigate the behavioral, algorithmic, and dopaminergic mechanisms governing cue-reward learning rate. Across a range of conditions in mice, we show a strong, mathematically proportional relationship between both behavioral and dopaminergic learning rates and the duration between rewards. Due to this relationship, removing up to 19 out of 20 cue-reward pairings over a fixed duration has no influence on overall learning. These findings are explained by a dopamine-based model of retrospective learning, thereby providing a unified account of the biological mechanisms of learning.

## Introduction

A perplexing feature of learning is that learning becomes more effective per experience when experiences are more temporally spaced (“spacing effect”) (1, 2). This concept is so widely known that students are regularly advised that study sessions spread over time are more effective than “cramming” before an exam (3–5). Such spacing effects have been demonstrated across many domains of learning in species ranging from invertebrates to humans (1, 6–13), including mammalian cue-reward learning (14–20) (i.e., learning that a cue predicts/precedes reward). Whether spacing effects on cue-reward learning rate are a function of the spacing between cues, the spacing between rewards, or some combination thereof could reveal fundamental algorithms of learning. Yet, most demonstrations of faster learning with more spacing are qualitative and do not propose a fundamental mathematical rule governing learning rate. In contrast, a meta-analysis has suggested that under some settings, learning rate is mathematically proportional to the ratio of the spacing between cue-reward experiences (commonly referred to as “intertrial interval” or ITI) to the cue-reward interval (21, 22). However, whether this rule is a general property of learning is unknown (23–25).

Why is discovering such a mathematical rule governing learning rate important? For one, it can instruct principles to optimize learning from each experience. Further, identification of quantitative empirical rules has historically been the basis for discoveries of many fundamental mechanisms across fields. For instance, the mechanisms governing gravity (26), genetic inheritance (27), and action potential generation (28) are all examples of discoveries greatly aided by the observation of quantitative rules in experimental data (path of planetary orbits, proportion of heritable traits, and the conduction of action potentials, respectively). Yet, quantitative rules of learning rate dependence on trial spacing have largely been ignored in neurobiological conceptions of cue-reward learning, which have demonstrated that mesolimbic dopamine is a critical teaching signal for cue-reward learning (29–32). Therefore, neurobiological algorithms have primarily focused on explaining mesolimbic dopamine dynamics during such learning (33–43). However, a fundamental mechanism for cue-reward learning should not only capture known neurobiological findings related to dopamine dynamics, but also quantitative rules of learning rate control.

To address this major gap, here we investigate the behavioral, algorithmic, and dopaminergic mechanisms underlying learning rate control. Since quantitative rules of learning rate control have not been tested in trace conditioning—the dominant behavioral paradigm supporting models of dopamine function (33, 35)—we tested the impact of ITI on learning rate during cue-reward trace conditioning. We also measured the evolution of mesolimbic dopaminergic cue responses across conditioning (“dopaminergic learning”). Surprisingly, we find that there is a strong, mathematically proportional relationship between both behavioral and dopaminergic learning rate and the duration between rewards (“inter-reward interval”, IRI)—a relationship different from that suggested by the earlier metaanalysis (21, 22). As a consequence, we empirically demonstrate that 1) removal of up to 19 out of 20 cue-reward experiences has zero impact on overall behavioral and dopaminergic learning if total conditioning time is maintained (thereby implying that repetition over a fixed total time has no impact on learning), 2) for a behavior that typically takes hundreds of trials to learn with commonly used short ITIs (35, 42, 44), dopaminergic learning with a one hour IRI occurs in 2 trials and behavioral learning occurs in 3-4 trials, and 3) behavioral and dopaminergic learning only require half as many rewards when reducing reward probability by 50% (consistent with a doubling of the IRI and learning rate). Unlike other models, a model of retrospective learning triggered by rewards (i.e., learning whether a cue precedes reward) that accounts for known mesolimbic dopaminergic dynamics (41) naturally explains the scaling of behavioral and dopaminergic learning rates by IRI.

## Results

### Behavioral learning in one-tenth the experiences with ten times the trial spacing

We wanted to test whether there are strong, mathematical rules underlying behavioral learning rate in a behavioral task commonly used to study dopamine function. Since prior studies suggesting proportionality between learning rate and the ratio of ITI to trial duration primarily tested ratios within an order of magnitude, we sought to first test whether such a rule holds in trace conditioning, and if so, whether it holds over much larger ratios. To this end, we classically conditioned two groups of thirsty head-fixed mice with an ITI either >1 or >2 orders of magnitude longer than the trial period. Mice were conditioned to associate a brief auditory tone (0.25 s, 12 kHz) with the delivery of sucrose solution reward (15% w/v, 2-3 µL) through a spout positioned in front of their mouth (45) (**Fig. 1A**). Two groups of mice were presented with this same trial structure, with one group experiencing 60 s ITIs (“60 s ITI” mice) and another group experiencing 600 s ITIs (“600 s ITI” mice). Both groups were trained for 1hr per day. So, 60 s ITI mice were presented 50 cue-reward pairings a day, while 600 s ITI mice were presented 6 cue-reward pairings a day (this accounts for a fixed reward consumption period; see *Methods*). The head-fixed preparation is critical to test long ITIs with brief trial periods. By placing mice at a fixed distance from the spout, brief cues can be used, which allows for conditioning with very short trial periods relative to ITI. This approach also enables conditioning to begin without the need for pre-training mice to collect rewards, which can lead to the formation of other learned associations.

**Fig. 1.**
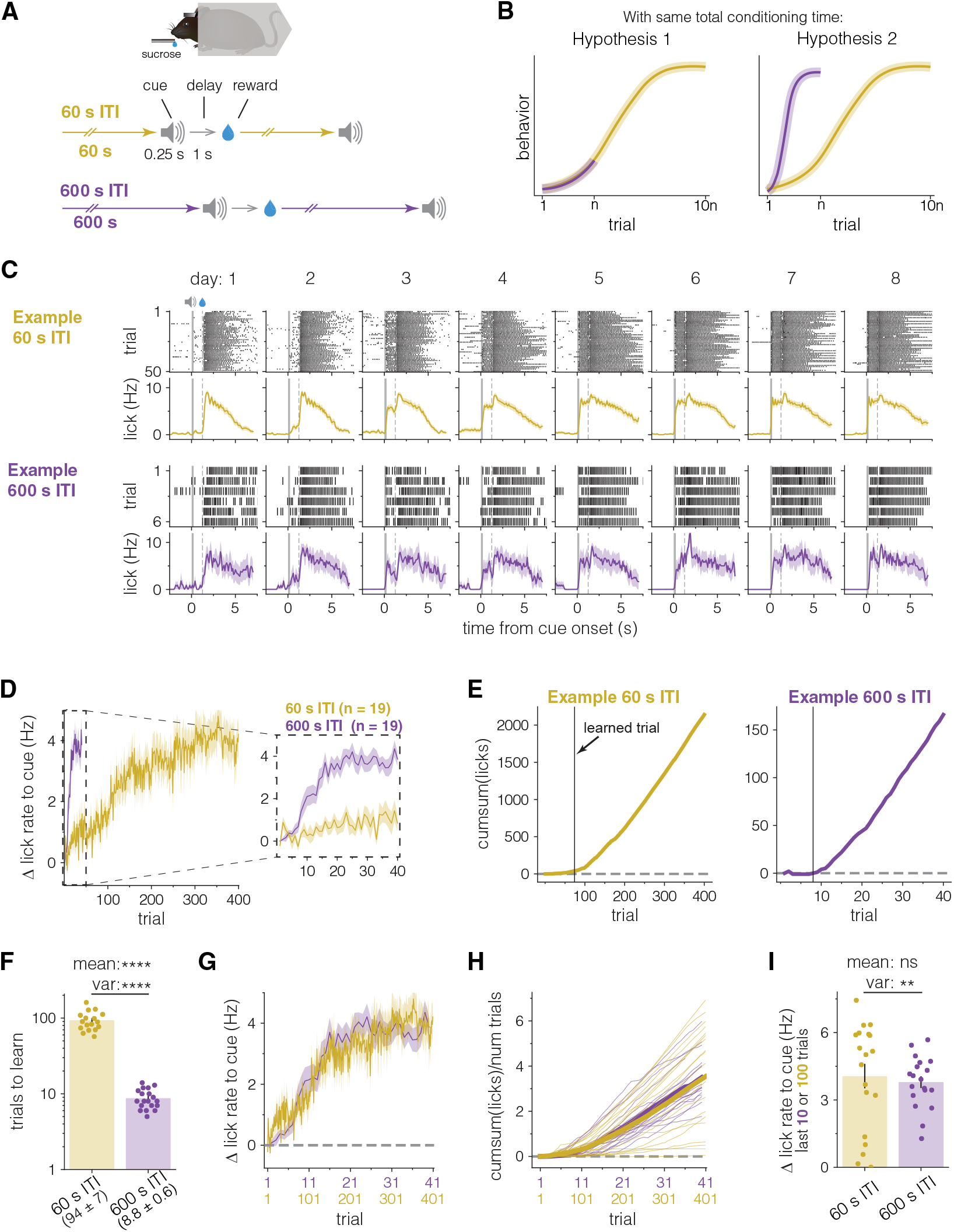
Behavioral learning in one-tenth the experiences with ten times the trial spacing. **A**, Schematic of experimental setup. Head-fixed mice were divided into two groups that were each presented with identical cue-reward pairing trials, but each group differed in the duration from a reward to the next cue (i.e., the inter-trial interval or ITI). Mice were conditioned for one hour per day resulting in 50 (60 s ITI) or 6 (600 s ITI) trials per session (see *Methods*). **B**, Illustration of two hypothetical experimental outcomes. Learning curves display the possible relationship between group-averaged learning rates for 60 s and 600 s ITI as a function of trial number. **C**, Example lick raster plots (*upper row*) and lick peri-stimulus time histograms (PSTH) (*lower row*) for one example mouse from either 60 s ITI group (*top*, gold) or 600 s ITI group (*bottom*, purple) showing every cue and reward presentation across eight days of conditioning. Each column represents a single day of conditioning. Graphs are aligned to cue onset (cue duration denoted by gray shading). Reward delivery is denoted by the vertical gray dashed line. This convention is followed for all similar figures later in the manuscript. Both example animals begin to show evidence of learning (an increase in licking following cue onset before reward is delivered) on day 2. **D**, 600 s ITI mice learn and reach asymptotic behavior in fewer trials than 60 s ITI mice. *Left*, Timecourse showing the average change in cue-evoked lick rate (the baseline subtracted lick rate between cue onset and reward delivery, see *Methods*) over 40 (600 s ITI, purple, n = 19 mice) or 400 (60 s ITI, gold, n = 19 mice) cue-reward presentations. *Inset right*, Zoom in of first 40 trials for both groups. Lines represent mean across animals and shaded area represents the SEM. **E**, Cumulative sum (cumsum) of cue-evoked licks across trials from the same example mice as in **C**. Using the cumsum curve from each animal to determine the trial after which mice first show evidence of learning (**fig. S2**; see *Methods*), we found that the example 60 s ITI mouse (left, gold) learns after trial 74, while 600 s ITI group (right, purple) learns after trial 8 (i.e., “few shot” reward learning). Learned trial is denoted by the solid black vertical line. **F**, 600 s ITI mice learn in about ten times fewer trials than 60 s ITI mice. Bar height represents mean trial after which mice show evidence of learning for 60 s ITI group (left, gold, n = 17) and 600 s ITI group (right, purple, n = 19), plotted on a log scale. Error bar represents SEM. Circles represent individual mice. Values under labels represent mean ± SEM. Two mice that did not show evidence of learning are excluded from comparison (**fig. S2**; see *Methods*). **** p < 0.0001, Welch’s t-test, F-test. **G - H**, On average, learning between groups progresses similarly as a function of total conditioning time, and thus scales with the ratio of ITIs. **G**. Average cue-evoked lick rates for 600 s ITI and 60 s ITI groups across scaled trial numbers (same data as in **D**), showing that the 600 s ITI group learns ten times more per experience compared to the 60 s ITI group. **H**. Cumsum of cue-evoked licks plotted on the same scaled x-axis. Thick lines represent group means and thin lines represent individual animals. Note the higher individual variability in the 60 s ITI group compared to the 600 s ITI group (quantified in **I**). **I**, Asymptotic cue-evoked lick rates have similar group means, but different variances. Bars represent mean cue-evoked lick rates during trials 301–400 (60 s ITI) or trials 31-40 (600 s ITI). Error bars represent SEMs and circles represent individual mice. ns: not significant, Welch’s t-test; **p<0.01, F-test. *See* ***table S1*** *for full statistical test details from all figures including test statistics, n’s, degrees of freedom, and both corrected and uncorrected p-values. All error bars and shading throughout the manuscript represent SEM unless otherwise noted. Values displayed under bar graph labels represent mean ± SEM*.

Using these groups of mice, we tested between two hypotheses (**Fig. 1B**). Hypothesis 1 is based on a trial-based learning framework, which predicts that once the ITI is sufficiently longer than the trial duration, trial-by-trial learning rate should be equivalent between groups presented with the same cue to reward delay, regardless of ITI. Because 60 s ITI mice will experience 10 times more cue-reward pairings, they will show greater evidence of overall learning than 600 s ITI mice. Hypothesis 2 instead is based on a mathematically proportional relationship between the learning rate per trial and trial spacing. Stated differently, hypothesis 2 is that 10 times increase in ITI (and thus, 10 times increase in IRI and “inter-cue interval”, ICI) will produce 10 times fewer experiences but 10 times learning per experience to result in equivalent overall learning. If true, this means that removing 9 out 10 experiences from 60 s ITI mice has no influence on overall learning.

We measured behavioral learning using cue-evoked anticipatory licks before reward delivery (35, 44, 46). Mice from both groups began to show cue-evoked licks in the first few days of conditioning (**Fig. 1C** and **fig. S1A**). When looking at cue-evoked licking as a function of cue-reward experiences, however, 600 s ITI mice learned and reached asymptotic behavior in many fewer trials than 60 s ITI mice (**Fig. 1D**). By trial 40, 600 s ITI mice showed significantly more cue-evoked licking (60 s ITI: 1.1 ± 0.4 Hz, 600 s ITI: 3.7 ± 0.3 Hz, p<0.0001; **Fig. 1D** and **fig. S1B**) and were significantly more likely to respond to the cue (60 s ITI: 0.29 ± 0.06, 600 s ITI: 0.92 ± 0.04, <0.0001; **fig. S1, C and D**) than 60 s ITI mice.

To fully compare learning rates between groups, we determined the trial after which each mouse showed evidence of learning using the cumulative sum of cue-evoked licks (17, 41, 47–50) (see *Methods*; **Fig. 1E** and **fig. S2, A to D**). Remarkably, 600 s ITI mice learned in 9 trials on average (8.8 ± 0.6), significantly less than the 94 (94 ± 7) trials needed for 60 s ITI mice to learn (p<0.0001; **Fig. 1F**). By lengthening the ITI by a factor of 10, cue-reward learning required 10 times fewer trials, showing a quantitative, proportional scaling relationship between trial spacing and per-trial learning. This scalar relationship was not just limited to the learned trial number, as a single trial for 600 s ITI mice was worth 10 trials for 60 s ITI mice throughout the learning process (**Fig. 1, G and H**, and **fig. S2, E and F**). Because 600 s ITI mice have the same experience as 60 s ITI mice but with the removal of 9 out of 10 trials (i.e., 10 times the ITI), the overlap of the learning curves demonstrates that those missing trials have no effect on overall learning across cumulative conditioning time.

Further suggesting that learning between groups was simply scaled, average asymptotic cue-evoked lick rates (60 s ITI: 4.06 ± 0.54 Hz, 600 s ITI: 3.80 ± 0.26 Hz, p = 0.66; **Fig. 1I**), the likelihood of responses to the cue (60 s ITI: 0.77 ± 0.08, 600 s ITI: 0.92 ± 0.03, p = 0.098; **fig. S1G**), and the abruptness of change, a measure of the steepness of individual animal learning curves (60 s ITI: 0.18 ± 0.02, 600 s ITI: 0.18 ± 0.02, p = 0.97; **fig. S1H**), were all similar between groups at the end of conditioning. Interestingly, despite similar average rates of asymptotic cue-evoked licking, 60 s ITI mice showed significantly more variance in individual behavior compared to 600 s ITI at the end of conditioning (p<0.01; **Fig. 1, H and I**; two 60 s ITI mice did not learn the cue-reward association, **fig. S2C**). This variance was also seen in the number of trials to learn when comparing mice that showed evidence of learning (p<0.0001; **Fig. 1F**). This suggests that individual variability in learning is driven in part by environmental factors and is not just a reflection of innate abilities.

### Dopaminergic learning in one-tenth the experiences with ten times the trial spacing

The dominance of trialbased accounts of associative learning is supported in large part by the concordance between mesolimbic dopamine signaling and the reward prediction error (RPE) term in temporal difference reinforcement learning (TDRL) models (34–36, 46, 51–53). In temporal difference cue-reward learning, the goal is to use RPEs to estimate the value of a cue, which is used to drive behavior. If the value guiding behavior is learned using a dopaminergic error signal, dopamine should be tightly coupled to behavior (33, 46, 54). Thus, to understand how our results of learning rate scaling fit with common conceptions of dopamine in associative learning, it is important to understand how dopamine release to the cue evolves over the course of learning (i.e., “dopaminergic learning”) in both 60 s ITI and 600 s ITI mice (**Fig. 2**). Given the vastly different number of trials to acquisition in each group, we hypothesized two possible relationships between dopaminergic and behavioral learning (**Fig. 2B**). Hypothesis 1 is that the development of cue-evoked dopamine precedes the emergence of learned behavior by a fixed number of trials in both groups. Because 600 s ITI mice learn in ten times fewer experiences than 60 s ITI mice (**Fig. 1F**), Hypothesis 2 is that the development of cue-evoked dopamine also occurs proportionally earlier in ten times fewer experiences and hence precedes behavioral learning by ten times fewer experiences.

**Fig. 2.**
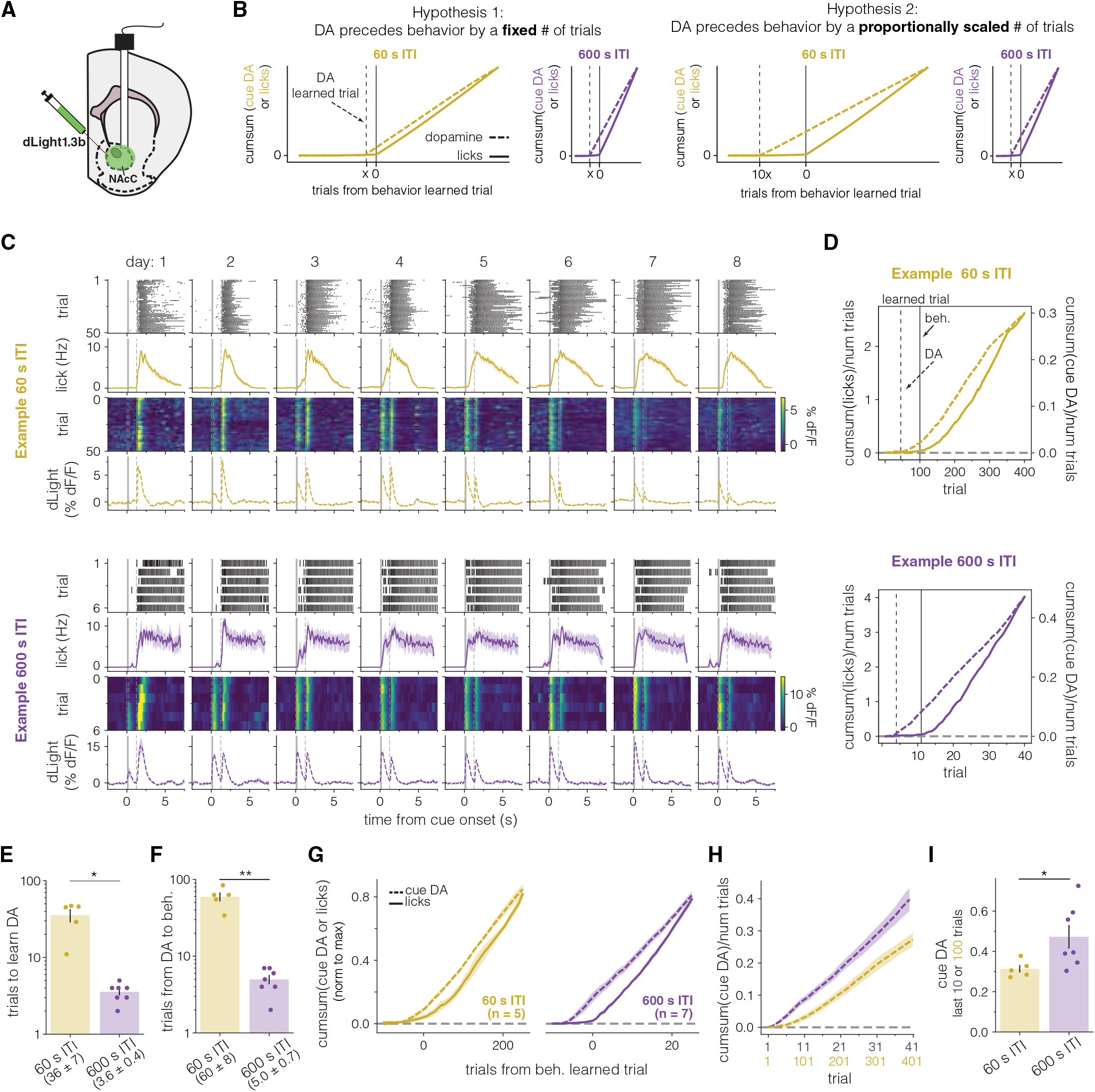
Dopaminergic learning in one-tenth the experiences with ten times the trial spacing. **A**, Schematic of mesolimbic dopamine measurements from nucleus accumbens core (NAcC; see *Methods*). **B**, Diagrams of two hypothetical relationships between dopaminergic and behavioral learning in 60 s and 600 s ITI mice. **C**, Example lick raster plots (*upper row*), lick PSTH (*2*^*nd*^ *row*), heatmap of dopamine responses on each trial (*3*^*rd*^ *row*) and average dopamine PSTH for the day (*lower row*) for one example mouse from either 60 s ITI group (*top*, gold) or 600 s ITI group (*bottom*, purple) showing every cue and reward presentation across eight days of conditioning. Lick data presented as in **Fig. 1C**. Dopamine signals plotted as % dF/F. Graphs aligned to cue onset (cue duration denoted by gray shading). Reward delivery is denoted by vertical gray dashed line. **D**, Cumsum of cue-evoked licks (solid, left axis) or of cue-evoked dopamine (dashed, right axis) for the same example mice as in **C**. Both lick and cue-evoked dopamine values were divided by total trial number to display average responses across conditioning. Before taking the cumsum, cue-evoked dopamine responses were normalized by max reward responses for each animal (see *Methods*). Cumsum curves were used to determine the trial after which cue-evoked dopamine or cue-evoked licking emerge (“learned trial”, see *Methods*). Solid vertical lines represent learned behavior trial and dashed vertical lines represent dopamine learned trial. This convention is followed for all similar figures later in the manuscript. **E**, Dopamine responses to cue develop in ten times fewer trials in 600 s ITI mice compared to 60 s ITI mice. Values under labels represent mean ±SEM across mice. * p<0.05, Welch’s t-test. **F**, On average, DA cue responses develop 60 trials before the emergence of cue-evoked licking in 60 s ITI mice and 5 trials before in 600 s ITI mice. **p<0.01. Welch’s t-test. **G**, Average cumsum of cue-evoked licks (solid) and dopamine (dashed) across groups. Data were normalized by each animal’s final trial of conditioning and aligned to their learned trial before averaging. Lines represent means, and shading represents SEM. **H**, Average cumsum of normalized cue-evoked dopamine responses in 60 s ITI (gold) and 600 s ITI (purple) mice. Cumsum curves divided by number of trials to account for differences between groups. **I**, Mean asymptotic normalized cue-evoked dopamine after learning. Bars represent means for trials 301-400 (60 s ITI) or trials 31-40 (600 s ITI). Error bars represent SEM and circles represent individual mice. *p<0.05, Welch’s t-test.

To test these hypotheses, we measured dopamine release in the nucleus accumbens core in a subset of 60 s ITI and 600 s ITI mice with fiber photometry recordings of the optical dopamine sensor dLight1.3b (**Fig. 2A** and **fig. S3**). As seen in example mice, dopamine was evoked by reward receipt beginning on the first trial, but cue-evoked dopamine release developed over trials and preceded the emergence of behavioral learning, in line with prior work (41, 51, 55–57) (**Fig. 2C** and **fig. S3A**). To determine the trial at which cue-evoked dopamine emerged, we applied the same algorithm used to determine the behavior learned trial on the cumulative sum of the cue-evoked dopamine (**Fig. 2D** and **fig. S3B**; see *Methods*). Again, we found the same proportional scaling relationship between IRI and dopaminergic learning. Dopaminergic learning in 600 s ITI mice began on average after trials three or four (3.6 ± 0.4), significantly earlier than 60 s ITI mice, which began to show cue-evoked dopamine responses after trial 36 (36 ± 7) (p<0.05; **Fig. 2E**). We then calculated the lag between dopaminergic and behavioral learning. In 600 s ITI mice, dopaminergic learning precedes behavior by 5 trials on average (5.0 ± 0.7), significantly fewer than the 60 (60 ± 7) trials between dopaminergic and behavioral learning in 60 s ITI mice (p<0.01; **Fig. 2F**). Thus, per-trial development of cue-evoked dopamine responses also scales proportionally with trial spacing across learning: by increasing the ITI (and thus IRI and ICI) by a factor of ten, cue-evoked dopamine appears in ten times fewer trials and precedes behavioral learning in ten times fewer trials (**Fig. 2G** and **fig. S4C**). This scaling is consistent with Hypothesis 2 (**Fig. 2B**).

Interestingly, despite the scaling in the onset of dopaminergic learning, cue-evoked dopamine in 600 s ITI mice rose to asymptotic levels more rapidly and increased by more than a factor of ten per trial as compared to 60 s ITI mice (**Fig. 2H** and **fig. S4, D to F**). Asymptotic cue-evoked dopamine (relative to maximum reward response) was also significantly higher at the end of conditioning in 600 s ITI mice compared to 60 s ITI mice (0.47 ± 0.06 vs. 0.31 ± 0.02, p<0.05; **Fig. 2I**).

Furthermore, although there were differences in dopamine dynamics and learning rates between the groups, we observed a similar pattern in the dopamine reward response in both groups. Specifically, the reward response did not start at its maximum value during the first trial but rather increased during early conditioning as observed previously (41, 42), reaching its peak prior to the onset of behavior (**fig. S4, G and H**).

### Learning rate scales proportionally with inter-reward interval

To further probe the relationship between trial spacing and learning rate, we conditioned two additional groups of mice with the same cue-reward delay as above separated by either a 30 s (“30 s ITI” mice, 100 trials/day) or 300 s (“300 s ITI” mice, 11 trials/day) ITI (**Fig. 3, A to C**). 300 s ITI mice learned and reached asymptotic behavior in many fewer trials than 30 s ITI mice (**Fig. 3D** and **fig. S5A**), showing significantly more cue-evoked licking by trial 80 (30 s: 1.0 ± 0.5 Hz, 300 s: 4.8 ± 0.6 Hz, p <0.001; **fig. S5B**). When examining individual learned trials, we found that similar to 600 vs 60 s ITI mice, 300 s ITI mice learned in about ten times fewer trials than 30 s ITI mice (300 s: 16.7 ± 4.1; 30 s: 176 ± 36, p<0.05; **Fig. 3E** and **fig. S5C**), while showing comparable asymptotic lick rates (30s: 3.9 ± 1 Hz, 300s: 4.7 ± 0.7 Hz, p = 0.49; **Fig. 3F** and **fig. S5D**). Since total duration between trials in the experiments so far is equal to the IRI (or equivalently, the ICI since reward probability is 100%), here, we calculate learning curves as a function of IRI (though additional experiments are needed to test whether the learning rate is specifically proportional to the IRI). If trial numbers are scaled by IRI to maintain equivalent conditioning time across groups the learning of all four groups of mice, 30 s – 600 s ITI, progressed similarly over conditioning (**Fig. 3G**). When further quantifying the relationship between IRI and learning rate on a log-log plot, we found a strong linear relationship between log(trials to learn) and log(IRI) with a slope statistically indistinguishable from -1 (-1.06, p = 0.107, **Fig. 3H**), indicating inverse proportionality between trials to learn and IRI, or a proportional relationship between learning rate and IRI: for every increase in IRI by a factor of n, animals learn in n times fewer trials. In addition, the slope of the line between the mean dopamine learned trials for 60 s and 600 s ITI mice as a function of IRI on a log-log plot was -1.03, suggesting the same quantitative proportional scaling relationship holds for dopaminergic learning (**fig. S5E**)

**Fig. 3.**
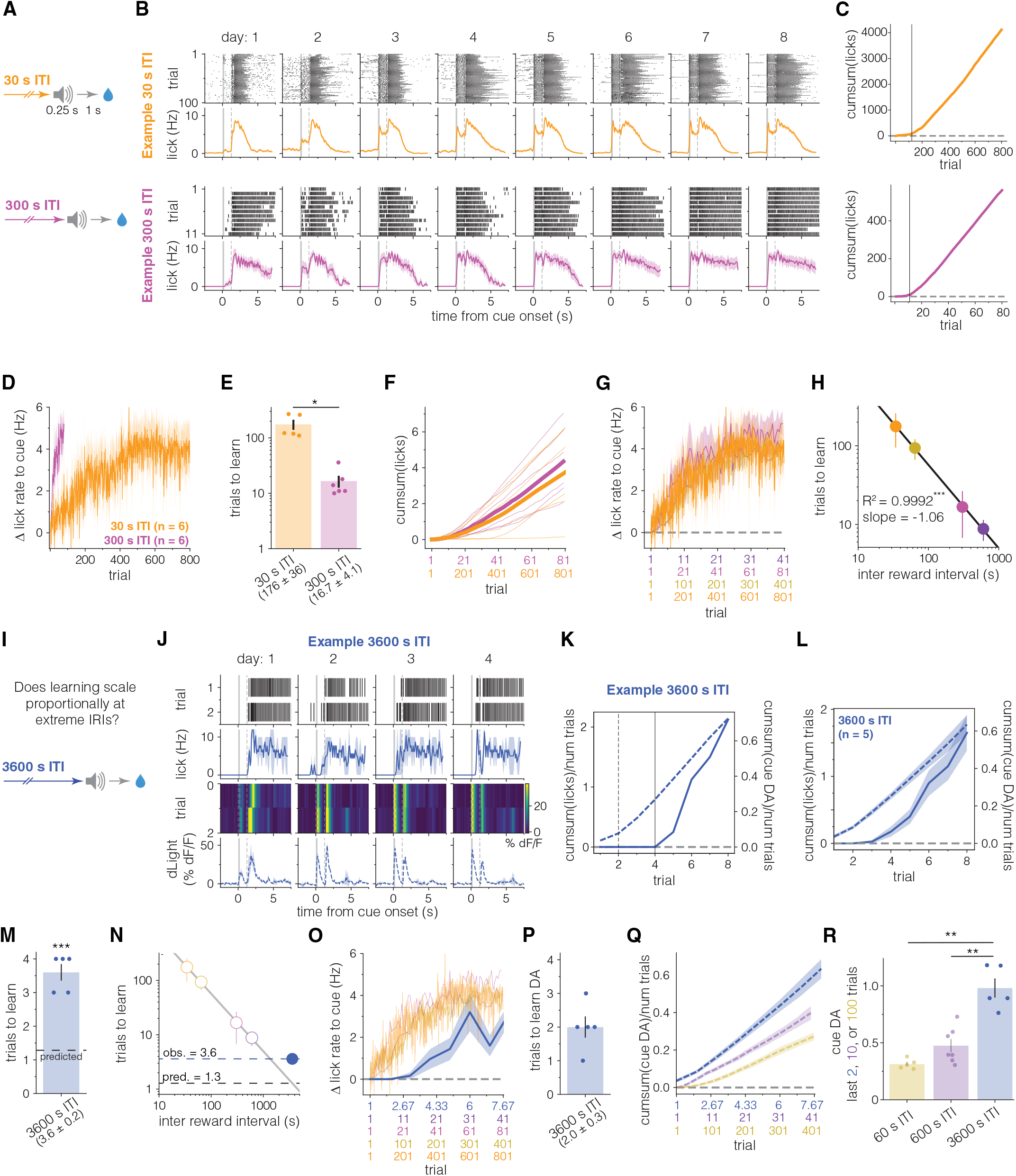
Learning rate scales proportionally with reward frequency across a range of trial spacing intervals. **A**, 30 s ITI and 300 s ITI conditioning (total number of trials a day 30 s: 100, 300 s: 11). **B**, Example lick raster plots (*upper row*) and lick PSTH (*lower row*) for one example mouse from either 30 s ITI (*top*, orange) or 300 s ITI groups (*bottom*, pink) showing every cue and reward presentation across eight days of conditioning. Data presented as in **Fig. 1C. C**, Cumsum of cue-evoked licks across trials from the same example mice as in **B**. Learned trial is denoted by the solid black vertical line. **D**, 300 s ITI mice learn and reach asymptotic behavior in fewer trials than 30 s ITI mice. Timecourse showing the average change in cue-evoked lick rate over 80 (300 s ITI, pink, n = 6) or 800 (30 s ITI, orange, n = 6) cue-reward presentations. **E**, 300 s ITI mice learn in about ten times fewer trials than 30 s ITI mice. One mouse that did not show evidence of learning is excluded from comparison (**fig. S5C**; see *Methods*). * p < 0.05, Welch’s t-test. **F**, Cumsum of cue-evoked licks (i.e., of data in **D**) plotted across scaled trials. **G**, Average cue-evoked lick rates for 30 s (orange, same data as **D**), 60 s (gold, same data as **Fig. 1, D and G**), 300 s (pink, same data as **D**), and 600 s (purple, same data as **Fig. 1, D and G**) across scaled trial numbers. On average, learning between groups scales with the ratio of ITIs and progresses similarly as a function of total conditioning time. Lines represent mean across animals and shaded area represents the SEM. **H**, Mean trials to learn for different ITI groups as a function of inter-reward interval (IRI) plotted on a log-log axis. Circles represent mean trials to learn per group and error bars represent standard deviation. Solid black line is best fit regression line (R^2^ = 0.9992, *** p<0.001). Slope is not significantly different from -1 (one-sample t-test) indicating a proportional quantitative scaling relationship between IRI and learning rate (1/trials to learn). **I**, A group of animals with a 3600 s ITI were conditioned similarly to previous groups to test if the observed power law relationship between IRI and trials to learn holds at extreme IRIs. Animals were presented with two cue-reward pairings per session for a total session duration of 2 hours. **J**, Example lick raster plots (*upper row*), lick PSTH (*2*^*nd*^ *row*), heatmap of dopamine responses on each trial (*3*^*rd*^ *row*) and dopamine PSTH (*lower row*) for one example 3600 s ITI mouse showing every cue and reward presentation across eight days of conditioning. Data presented as in **Fig. 2C. K**, Cumsum of cue-evoked licks (solid, left axis) or of normalized cue-evoked dopamine (dashed, right axis) for the same example mouse as in **J**. Data presented as in **Fig. 2D. L**, Average cumsums of cue-evoked licks and normalized cue-evoked dopamine in 3600 s ITI mice (n = 5). **M**, Number of trials for 3600 s ITI (n = 5) mice to learn cue-reward association. Bar height represents mean trial after which mice show evidence of learning. Dashed line represents predicted trials to learn based on relationship observed for 30 s – 600 s ITI groups (best fit line in **H**). Values under labels represent mean ± SEM. *** p< 0.001, one-sample t-test. **N**, 3600 s ITI mice (blue circle, filled) take more trials to learn than predicted from the proportional scaling relationship in **H**. White filled circles and line are same data as **H**. Dashed lines represent predicted and mean observed trials to learn for 3600 s ITI. **O**, Learning in 3600 s ITI mice does not scale proportionally with other groups. Average cue-evoked lick rates for 30 s, 60 s, 300 s, 600 s, (30-600 s ITI, same data as **G**) and 3600 s ITI (blue, n = 5) mice across scaled trial numbers. Previously displayed data are shown without error to aid visualization. **P**, Mean dopamine learned trial for 3600 s ITI mice (n = 5). Values under labels represent mean ± SEM. **Q**, Average cumsum of normalized cue-evoked dopamine responses in 60 s ITI (gold, same data as **Fig. 2H**), 600 s ITI (purple, same data as **Fig. 2H**), and 3600 s ITI (blue, n = 5) mice. Cumsum curves divided by number of trials to account for differences between groups. Transparency added to previously displayed data to aid visualization. Shaded region represents SEM. **R**, Asymptotic normalized cue-evoked dopamine after learning is highest in 3600 s ITI mice. Bars represent means for trials 301-400 (60 s ITI, same data as **Fig. 2I**), trials 31-40 (600 s ITI, same data as **Fig. 2I**), or trials 15-16 (3600 s ITI mice). Transparency added to previously displayed data to aid visualization. *p<0.05, Welch’s t-test.

This strong proportional relationship between learning rate and IRI implies that at extreme IRIs, animals could learn cue-reward associations in one or two trials. To test this, we conditioned another group of mice while recording dopamine with the same trial structure as previous groups, but with a 3600 s ITI (“3600 s ITI mice”, 2 trials per day, 2-hour session time; **Fig. 3I**), which is predicted to lead to behavioral learning within 1.3 trials based on the relationship observed above (**Fig. 3H**). While 3600 s ITI mice rapidly learned the cue-reward association within 3.6 trials on average, this was significantly more than predicted (p<0.001, **Fig. 3, J to N**, and **fig. S6A**). However, 3600 s ITI mice did learn significantly faster than 600 s ITI mice (p<0.0001, **fig. S6B**). Thus, extending the ITI or IRI increases the learning rate, but the proportional scaling relationship breaks down at extreme intervals (**Fig. 3, N to O**). Similar results were observed when examining the emergence of cue-evoked dopamine. Dopaminergic learning occurred after 2 trials on average (**Fig. 3P**), significantly fewer than in 600 s ITI mice (p<0.01, **fig. S6E**), but not in line with the proportional scaling relationship seen with shorter ITIs and IRIs (**fig. S6D**). Interestingly, cue-evoked dopamine in 3600 s ITI mice had a significantly higher asymptote (0.98 ± 0.08) than in 60 s ITI (0.31± 0.02, p<0.01) or 600 s ITI mice (0.47± 0.06, p<0.01) (**Fig. 3, Q and R** and **fig. S6, F to K**). Thus, the proportional scaling of learning rate with IRI breaks down at extreme IRIs, but extreme IRIs nevertheless increase learning rate and asymptotic dopamine.

### ANCCR, a model of retrospective causal learning, captures the proportional scaling of learning rate by inter-reward interval

Given the experimentally observed proportional scaling of learning rate with the time between trials, we assessed whether any model of learning can naturally explain these results. We recently proposed a new framework of behavioral and dopaminergic learning named Adjusted Net Contingency of Causal Relations (ANCCR) that operates retrospectively at the time of reward to learn whether a cue consistently precedes it (41). The learned retrospective association (i.e., how often does the cue precede the reward?) is then converted to a prospective association (i.e., how often does the reward follow the cue?) using a form of Bayes’ rule. This states that if a cue precedes a reward consistently, then the reward follows the cue at a rate proportional to the ratio of the overall rate of the reward to the overall rate of the cue. Due to the retrospective nature of learning with associative updates only happening at rewards, the learning rate of a cue-reward association should be adjusted by reward frequency (i.e., frequency of retrospective updates) in ANCCR for the Bayes rule to be valid, such that for infrequent rewards, one should learn proportionally more compared to frequent rewards (i.e., 10 times learning per reward for a 10 times less frequent reward) (see *Methods*). Mathematically, the learning rate is predicted to be proportional to the IRI. This rule is consistent with the experimental results showing proportionality between learning rate and trial spacing (**Fig 3H**). As a result, it is consistent with the experimental observation that the total time needed to acquire conditioning is constant for a 20-fold variation in the rate of trials (**Fig 4, A and B**).

**Fig. 4.**
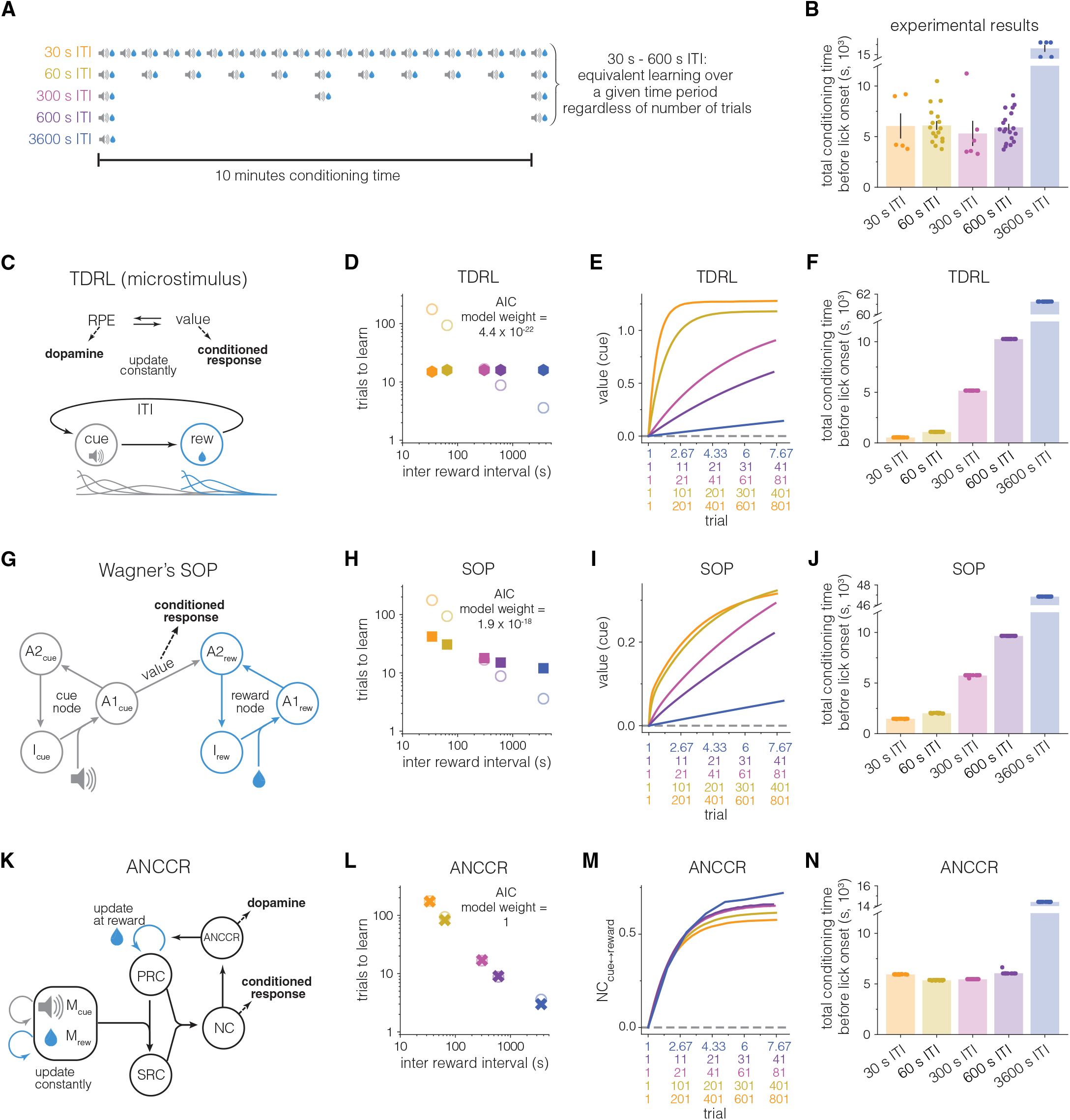
Proportional scaling of learning rate by inter-reward interval is only captured by a retrospective learning model. **A**, Diagram of number of trials experienced by each ITI group over a ten-minute interval. Based on the experimentally observed proportional scaling between trial spacing and learning rate (**Fig. 3H**), learning is equivalent during this interval regardless of the number of trials experienced for 30 s – 600 s ITI, but not 3600 s ITI mice. Note that multiple 3600 s trials are not shown to maintain the scale of the diagram. **B**, Total conditioning time prior to the emergence of anticipatory licking from experiments in **Figs. 1 and 3**. Symbols represent individual mice, and bar height represents mean across ITI group. **C**, Schematic diagram of microstimulus implementation of temporal difference reinforcement learning (TDRL). At each moment, a value estimate of future rewards is updated by a reward prediction error (RPE), representing a deviation from current value estimate that is thought to be encoded in the brain by dopamine signaling. Value is used to drive behavioral responding. The microstimulus implementation of TDRL was specifically chosen because external stimuli (i.e. cues and rewards) evoke microstimuli which spread representations of the external stimuli in time. Because the model contains ITI states that themselves can acquire value, the model contains a potential mechanism to explain how the spacing between trials could affect learning rate. Parameter combinations were swept to determine if any set could capture the quantitative scaling observed in the experimental results where learning rate varied proportionally with the IRI. **D**, Trials to learn (when value > threshold) as a function of IRI for TDRL simulation with parameter combination that best fits experimental results (see *Methods*). Hexagons represent mean trials to learn across iterations (n = 20 each) for all conditions. Open circles represent experimental data (same as **Fig. 3, H and N**). Akaike information criterion (AIC) model weight comes from comparison with best fit SOP (**F**), and ANCCR (**I**) models. Please note that this AIC model does not penalize free parameters, i.e., is a conservative estimate biased against ANCCR (see *Methods* for rationale and **table S1** for AIC model weight when penalizing for parameters). **E**, Timecourse of value on each trial (maximum between cue and reward) for best fit TDRL model (**B**) averaged across iterations (n = 20 each) and plotted across scaled trials for all conditions. SEM occluded by thickness of mean lines. **F**, Total conditioning time prior to the emergence of behavior (threshold crossing). Symbols represent time for a single iteration, and bar height represents mean (n = 20 iterations per ITI condition). **G**, Schematic diagram of Wagner’s SOP (Standard Operating Procedure or Sometimes Opponent Process) model of associative learning. Presentations of cues or rewards evoke presumed mental representations or processing nodes consisting of many informational elements. These stimulus representations are dynamic: presentation of a stimulus moves a portion of elements from (only) the inactive (I) state into the primary active state (A1). Elements then decay into the secondary active state (A2) and then decay again back to the inactive state while the stimulus is absent. Elements transition between states according to individually specified probabilities. Cue-reward associations (value) are strengthened and learned when cue elements in A1_cue_ and reward elements in A1_reward_ overlap in time. Following learning, cues evoke conditioned responding by directly activating reward elements to their A2 state. One way in which SOP has been hypothesized to explain ITI impact on learning is that more time between trials allows more elements to decay to the inactive state, allowing for greater number of elements to transition to the A1 active state upon next cue and reward presentation. Parameter combinations were swept through to determine if any set of parameters could capture the quantitative scaling observed in the experimental results. **H**, Trials to learn (when value > threshold) as a function of IRI for SOP simulation with best-fit parameter combination of experimental results (see *Methods*). Squares represent mean trials to learn across iterations (n = 20 each) for all conditions. Open circles represent experimental data (same as **Fig. 3, H and N**). Akaike information criterion (AIC) model weight comes from comparison with best fit TDRL (**B)** and ANCCR (**I**) models. Please note that this AIC model does not penalize free parameters, i.e., is a conservative estimate biased against ANCCR (see *Methods* for rationale and t**able S1** for AIC model weight when penalizing for parameters). **I**, Timecourse of value on each trial (maximum between cue and reward) for best fit SOP model (**F**) averaged across iterations (n = 20 each) and plotted across scaled trials. SEM occluded by thickness of mean lines. **J**, Total conditioning time prior to the emergence of behavior (threshold crossing). Symbols represent time for a single iteration, and bar height represents mean (n = 20 iterations per ITI condition). **K**, Schematic diagram of ANCCR. See text for explanation of learning rate scaling in ANCCR. **L**, Trials to learn (when NC_cue↔reward_ (net contingency) > threshold) as a function of IRI for ANCCR simulation with parameter combination that best fits experimental results (see *Methods*). Crosses represent mean trials to learn across iterations (n = 20 each). Open circles represent experimental data (same as **Fig. 3, H and N**). Akaike information criterion (AIC) model weight comes from comparison with best fit TDRL (**B**) and SOP (**F**) models. **M**, Timecourse of NC_cue↔reward_ on each trial for best fit ANCCR model (**I)** averaged across iterations (n = 20 each) and plotted across scaled trial units. SEM occluded by thickness of mean lines. **N**, Total conditioning time prior to the emergence of behavior (threshold crossing). Symbols represent time for a single iteration, and bar height represents mean (n = 20 iterations per ITI condition).

While the experimentally observed proportional scaling relationship is consistent with the predictions of ANCCR, this does not exclude the possibility that other models of learning may better fit the empirical results. To quantitatively test this, we considered three models of conditioning with the potential to explain the experimental findings (**Fig. 4**): the microstimulus implementation of TDRL (**Fig. 4C**), Wagner’s SOP (Standard Operating Procedure or Sometimes Opponent Process, **Fig. 4G)**, and ANCCR (**Fig. 4K**). The microstimulus implementation of TDRL (58) was chosen because it contains ITI states that can acquire value, thereby containing a mechanism to produce trial spacing effect (longer ITI results in lower ITI value and larger RPE at cue; similar to “context extinction”(16). Though it is not a dopaminergic learning model, Wagner’s SOP model (59, 60) was chosen because it has previously been suggested to explain the trial spacing effect. Finally, in ANCCR, associations are formed by retrospectively inferring the cause of rewards and predicts a strong proportional relationship between learning rate and IRI.

To determine whether TDRL, SOP, or ANCCR best fits the behavioral results, we performed a quantitative model comparison using an Akaike information criterion (AIC)-derived relative weight (see *Methods*). Over the range of the parameters tested, the microstimulus model with the best fit parameter combination did not capture the proportional scaling observed experimentally in either the value (presumed to drive behavior) or the RPE signal (presumed to be encoded by dopamine) and requires more task duration to learn with increasing trial spacing (AIC 445.5, model weight compared to best fit SOP and ANCCR 4.4 × 10^−22^ **Fig. 4, C to F**, and **fig. S7**). A TDRL model with learning rate scaled by IRI using the relationship derived from ANCCR provides an approximate fit to behavioral learning (AIC 352.7, model weight compared to best fit SOP and ANCCR 0.0643), but nevertheless requires more task duration to learn with increasing trial spacing (**fig. S7, A and D**). It further did not capture the asymptotic dopamine cue response. The best-fit SOP model exhibited a qualitative trial spacing effect (faster learning at longer ITIs) as discussed previously (16, 20, 25) but did not match the proportional scaling observed experimentally, and requires more task duration to learn with increasing trial spacing (AIC 373.5, model weight compared to best fit TDRL and ANCCR 1.9×10^−6^ **Fig. 4, F and G**, and **fig. S8**). Therefore, proportional scaling with trial spacing is categorically absent in SOP and common TDRL frameworks. In contrast to these models, ANCCR captured both the proportional scaling of learning rate and the asymptotic dopamine cue responses and requires constant task duration to learn with increasing trial spacing from 30 s to 600 s (AIC 347.2, model weight compared to best fit TDRL and SOP 1 **Fig. 4, I to K**, and **fig. S7, A and E**).

### Learning rate scaling is not explained by number of experiences per day, context extinction, overall rate of auditory stimuli, or overall rate of rewards

Though the proportional scaling of learning rate by IRI appears inconsistent with trial-based learning theories, many uncontrolled confounding variables preclude a strong conclusion. First, mice conditioned with longer ITIs and fewer rewards per day may experience identical cues and rewards as more salient either due to higher novelty or lower satiety. As more salient stimuli lead to greater conditioning in trial-based accounts of learning (61, 62), this is a possible way for trial-based learning theories to explain our results. Furthermore, overnight consolidation could be an important driver of learning, as mice from all groups learned the cue-reward association during days 2-3. To control for these factors, we conditioned an additional group of mice (“60 s ITI-few”) with the same ITI as 60 s ITI mice (mean: 60 s) and the same number of trials per day as 600 s ITI mice (six), while recording dopamine from a subset (**Fig. 5A**). Across trials, dopaminergic and behavioral learning in these mice progressed similarly to 60 s ITI mice (**Fig. 5B** and **fig. S9A**). Late in conditioning, both cue-evoked licking and dopamine in 60 s ITI-few mice was significantly lower than 600 s ITI mice, comparable to 60 s ITI mice during the same trial numbers (lick: 60 s ITI-few: 0.9 ± 0.3 Hz vs. 600 s ITI: 3.7 ± 0.3 Hz, p<0.0001; vs. 60 s ITI: 1.1 ± 0.4 Hz, p>0.99; dopamine: 60 s ITI-few: 0.16 ± 0.07 vs. 600 s ITI: 0.49 ± 0.08, p<0.05; vs. 60 s ITI: 0.01 ± 0.06, p = 0.33; **Fig. 5C** and **fig. S9B**). Further, over the first 6 trials of conditioning when satiety or day effects do not differ between groups, cue-evoked dopamine grows in magnitude in 600 s ITI mice, while remaining flat in both 60 s ITI and 60 s ITI-few mice (**fig. S9, C to E**). These results suggest that satiety is not the main factor driving differences in behavioral and dopaminergic learning between groups, as is further indicated by consistent consummatory licking on the first and last six trials of conditioning with short ITIs (**fig. S9F**).

**Fig. 5.**
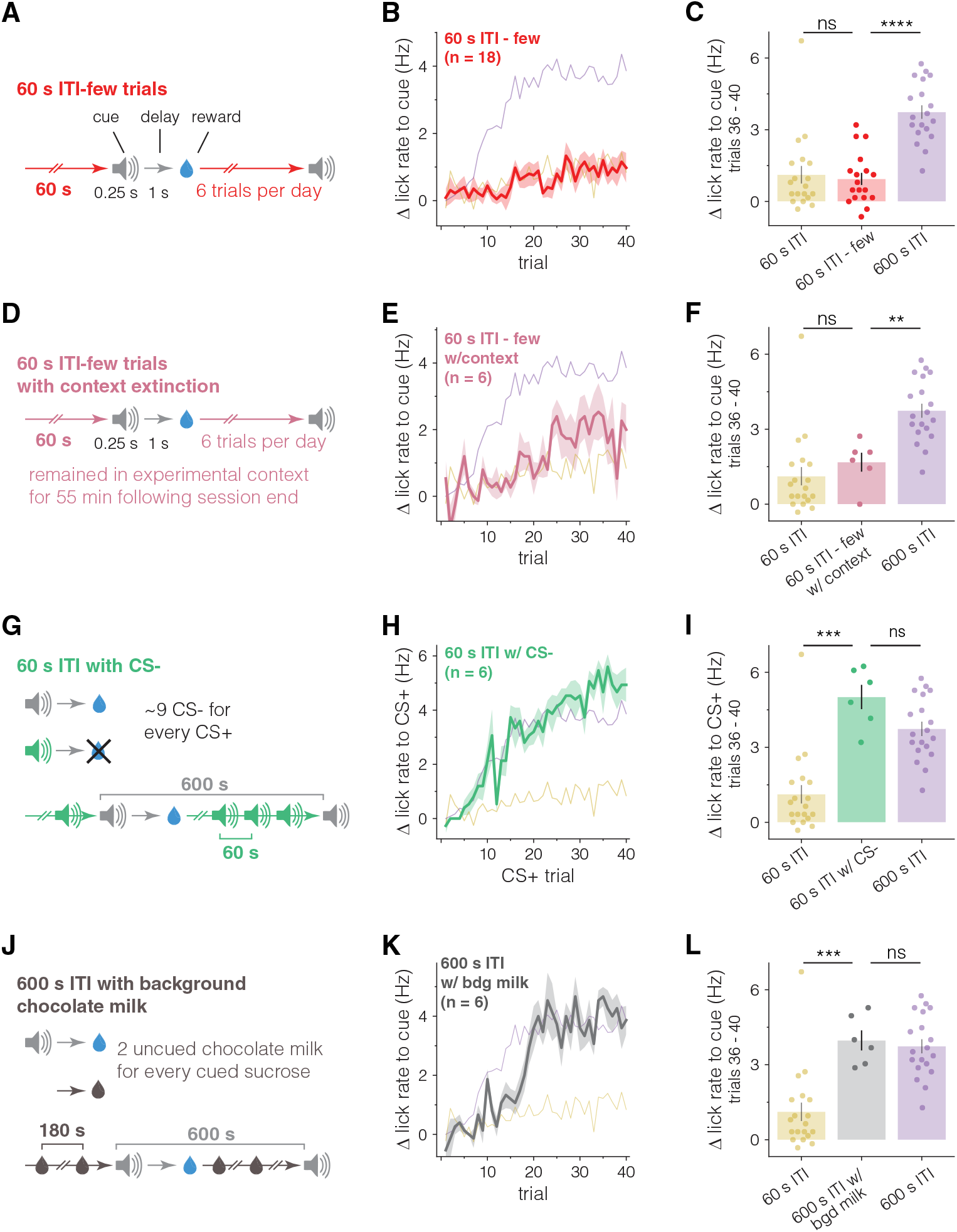
Learning rate scaling is not explained by number of experiences per day, context extinction, overall rate of auditory cues, or overall rate of rewards. A - C, Number of trials per day does not explain difference in learning rates between 60 s and 600 s ITI mice. **A** Schematic of conditioning for ‘60 s ITI-few’ group, which were conditioned with the same ITI as 60 s ITI mice (mean: 60 s) and the same number of trials/rewards per day as 600 s ITI mice (six). **B** Timecourse of average cue-evoked lick rate for 60 s ITI-few (red, n = 18), 60 s ITI (same data as **Fig. 1D**), and 600 s ITI (same data as **Fig. 1D**) mice over 40 trials of conditioning. 60 s and 600 s ITI timecourses are shown transparent and without error for visualization purposes. **C** Mean cue-evoked lick rate between trials 36 and 40 across all three groups. 60 s ITI-few mice show significantly less cue-evoked licking than 600 s ITI mice and behave like 60 s ITI mice. ****p<0.0001, ns: not significant; Welch’s t-tests. **D - F**, Context extinction does not explain difference in learning rates between 60 s and 600 s ITI mice. **D** Schematic of conditioning for “60 s ITI-few with context extinction” group. Mice were conditioned similarly to 60 s ITI-few mice but remained in the experimental context for ∼55 minutes following the end of conditioning trials, matching 600 s ITI group’s time in context and number of cue-reward experiences, while the rate of rewards during trials matched the 60 s ITI group. **E** Timecourse of average cue-evoked lick rate for 60 s ITI-few with context extinction (light pink, n = 6), 60 s ITI (same data as **Fig. 1D**), and 600 s ITI (same data as **Fig. 1D**) mice over 40 trials of conditioning. **F** Mean cue-evoked lick rate between trials 36 and 40 across all three groups. 60 s ITI-few with context extinction mice show significantly less cue-evoked licking than 600 s ITI mice and are not significantly different from 60 s ITI mice. **p<0.01, ns: not significant; Welch’s t-tests. **G - I**, Overall rate of auditory cues does not explain difference in learning rates between 60 s and 600 s ITI mice. **G** Schematic of conditioning for “60 s ITI with CS-” group. Mice were conditioned similarly to 600 s ITI mice but during the (∼600 s) interval between CS+→reward trials, distractor CS-cues (3kHz pure tone) were presented on average every 60 s to match the rate of auditory cues experienced by 60 s ITI mice. All mice could hear and respond to CS-as evidenced by some amount of generalized licking observed during conditioning (**fig. S10, C and D**). **H** Timecourse of average cue-evoked lick rate for 60 s ITI with CS-(green, n = 6), 60 s ITI (same data as **Fig. 1D**), and 600 s ITI (same data as **Fig. 1D**) mice over 40 trials of conditioning. **I** Average cue-evoked lick rate between trials 36 and 40 across all three groups. 60 s ITI with CS-mice show significantly more cue-evoked licking compared to 60 s ITI and are not significantly different from 600 s ITI mice. ***p<0.01, ns: not significant; Welch’s t-tests. **J - L**. Learning rate is not scaled by overall rate of rewards. **J** Schematic of conditioning for “600 s ITI with background chocolate milk” group. Mice were conditioned similarly to 600 s ITI mice but during the (∼600 s) interval between cue→sucrose trials, mice received 2 un-cued deliveries of chocolate milk ∼180 s apart to test whether cue-sucrose learning rate is affected by the general or identity specific rate of rewards. Mice readily consumed chocolate milk rewards upon delivery (**fig. S10H**). **K** Timecourse of average cue-evoked lick rate for 600 s ITI with background chocolate milk (gray, n = 6), 60 s ITI (same data as **Fig. 1D**), and 600 s ITI (same data as **Fig. 1D**) mice over 40 trials of conditioning. **L** Average cue-evoked lick rate between trials 36 and 40 across all three groups. 600 s ITI with background chocolate milk mice show significantly more cue-evoked licking compared to 60 s ITI and are not significantly different from 600 s ITI mice. ***p<0.01, ns: not significant; Welch’s t-tests.

A second confounding variable is the duration of unrewarded time in the conditioning context, which could extinguish context-reward associations with longer ITI and drive faster learning (16). To test this, we conditioned a group of mice (“60 s ITI-few with context extinction”) similarly to 60 s ITI-few mice but kept them in the experimental context for 55 minutes following the end of conditioning trials. This matches the time in context and number of cue-reward experience for 600 s ITI group and matches the rate of rewards during trials for the 60 s ITI group (**Fig. 5D**). As with 60 s ITI-few mice, learning in 60 s ITI-few with context extinction mice progressed similarly to 60 s ITI mice (**Fig. 5E**). Late in conditioning, 60 s ITI-few with context extinction mice licked significantly less to the cue (1.7 ± 0.4 Hz), than 600 s ITI mice (p<0.01) and similarly to 60 s ITI mice during the same trial numbers (p>0.99; **Fig. 5F**). Moreover, lick rates during the ITI positively correlate with learning rate within ITI groups, suggesting that context extinction is not the sole driver of cue-reward learning rate (**fig. S9, G to K**).

A third confounding variable is the overall rate of auditory stimuli during conditioning. The differing rates of auditory cues experienced by the different ITI groups could drive differences in learning rate by either affecting cue salience (sparser cues are more salient leading to increased learning of the cue) or by driving differences in replays/reactivations of cue-reward experiences (more time without external stimuli due to longer ITIs allows time for “virtual trials” to substitute for real cue-reward experiences) (63). To control for the overall rate of auditory cues, we conditioned an additional group of mice (“60 s ITI with CS-”) similarly to 600 s ITI mice but presented distractor CS-cues (3kHz tone) on average every 60 s during the (600 s) interval between CS+-reward trials to match the overall rate of auditory cues experienced by 60 s ITI mice (**Fig. 5G**). Across trials, learning in these mice progressed similarly to 600 s ITI mice (**Fig. 5H**). Late in conditioning, 60 s ITI with CS-mice licked to the CS+ (5.0 ± 0.5 Hz) comparable to 600 s ITI mice (p = 0.22), significantly more than 60 s ITI mice during the same trial numbers (p<0.001; **Fig. 5I**). When comparing the number of trials to learn, a similar relationship was observed: 60 s ITI with CS-mice learned the CS+-reward relationship in 9 trials, no different than 600 s ITI mice (p>0.99), yet significantly fewer trials than 60 s ITI mice (p<0.0001; **fig. S10, A and B**). Note that all mice could hear and respond to CS-cues as evidenced by some generalized licking observed during conditioning (**fig. S10, C and D**).

Finally, a fourth confounding variable is the overall rate of rewards. This is a confound because ANCCR predicts that the IRI of a specific reward identity (“identity specific IRI”), and not the overall reward rate, scales the learning rate. To test this, we conditioned an additional group of mice (“600 s ITI with background chocolate milk”) similarly to 600 s ITI mice but delivered 2 un-cued deliveries of chocolate milk during the (600 s) interval between cue-sucrose trials (**Fig. 5J**). Mice readily consumed chocolate milk rewards (**fig. S10H**). Across trials, learning in these mice progressed similarly to 600 s ITI mice (**Fig. 5K**). Late in conditioning, cue-evoked licking in 600 s ITI with background chocolate milk mice (4.0 ± 0.4 Hz) was comparable to 600 s ITI mice (p>0.99), significantly more than 60 s ITI mice during the same trial numbers (p<0.001; **Fig. 5L**). Based on the relationship between trials to learn and IRI observed above (**Fig. 3H**), if learning is scaled by a general IRI, 600 s ITI with background chocolate milk mice would be predicted to learn in 26.8 trials, while an identity specific IRI predicts learning in 8.5 trials. Mice learned in 12 ± 1 trials, significantly less than the general IRI prediction (p<0.0001), but also significantly more than both the identity specific IRI prediction (assuming perfect discriminability between reward types) (p<0.01; **fig. S10, E and F**) and 600 s ITI mice (p<0.001; **fig. S10G**). Because the learned trial is much closer to the prediction based on identity specific than overall rate of rewards, these results are consistent with learning rate scaling by an identity specific IRI with a small degree of generalization between rewards (likely due to their shared sensory properties as both are sweet liquids). Taken together, these results suggest that the learning rate across ITI conditions is scaled by an identity specific IRI and is not driven by satiety, overnight consolidation, the total number of cues and rewards experienced per day, context extinction, or the rate of auditory stimuli.

### Partial reinforcement scales learning rate by increasing the inter-reward interval

Though ANCCR predicts that the learning rate is scaled by IRI, the previous experiments did not disambiguate the IRI from the ICI for CS+ cues because they are equivalent when rewards follow cues with 100% probability. A counterintuitive prediction of the hypothesis that learning rate is scaled by IRI, and not ICI, is that reducing the probability of reward delivery, and thus increasing the IRI without affecting the ICI, would decrease the number of rewarded trials needed to learn. This contrasts with the prediction that learning rate is controlled by the ratio between ITI and trial duration, which implies a constant number of rewards to learn regardless of partial reinforcement (21). To test the prediction of IRI scaling, we conditioned an additional group of mice (“60 s ITI-50%”) identically to 60 s ITI mice (**Fig. 1**) but with 50% reward probability, while recoding dopamine in a subset. Reducing the reward probability by 50% leads to a doubling of the IRI to 120 s on average across a session while maintaining the ITI and ICI. (**Fig. 6A**). If the learning rate is scaled by the ICI, it would be predicted that mice would require 91.2 rewarded trials to learn the cue-reward association, based on the proportional scaling relationship discovered above (**Fig. 3H**), similar to 60 s ITI mice. However, if the learning rate is scaled by the IRI, it would be predicted that mice would need 43.8 rewarded trials to learn the association. Remarkably, 60 s ITI-50% mice took 42 ± 5 rewarded trials to learn, significantly less than 60 s ITI mice (p<0.0001; **Fig. 6, B and C**) or the ICI predicted 91 (p<0.0001), but not different from the IRI predicted 44 (p>0.99; **fig. S11A**). This IRI scaled relationship also held for dopaminergic learning as the emergence of cue-evoked dopamine took 20 ± 4 rewarded trials, significantly less than the 35.6 predicted by ICI scaling (p<0.05) and not different from the 17.5 predicted by IRI scaling (p>0.99; **fig. S11, B and C**). As rewarded trials make up half the experienced trials, 60 s ITI-50% mice learn the behavior in an equivalent number of trials/cue-presentations (87 ± 6) as 60 s ITI mice (p = 0.44; **fig. S11, D and E**) despite only half being rewarded. This same relationship holds for dopaminergic learning as well (46 ± 6 trials, p = 0.286 vs. 60 s ITI, **fig. S11F**).

**Fig. 6.**
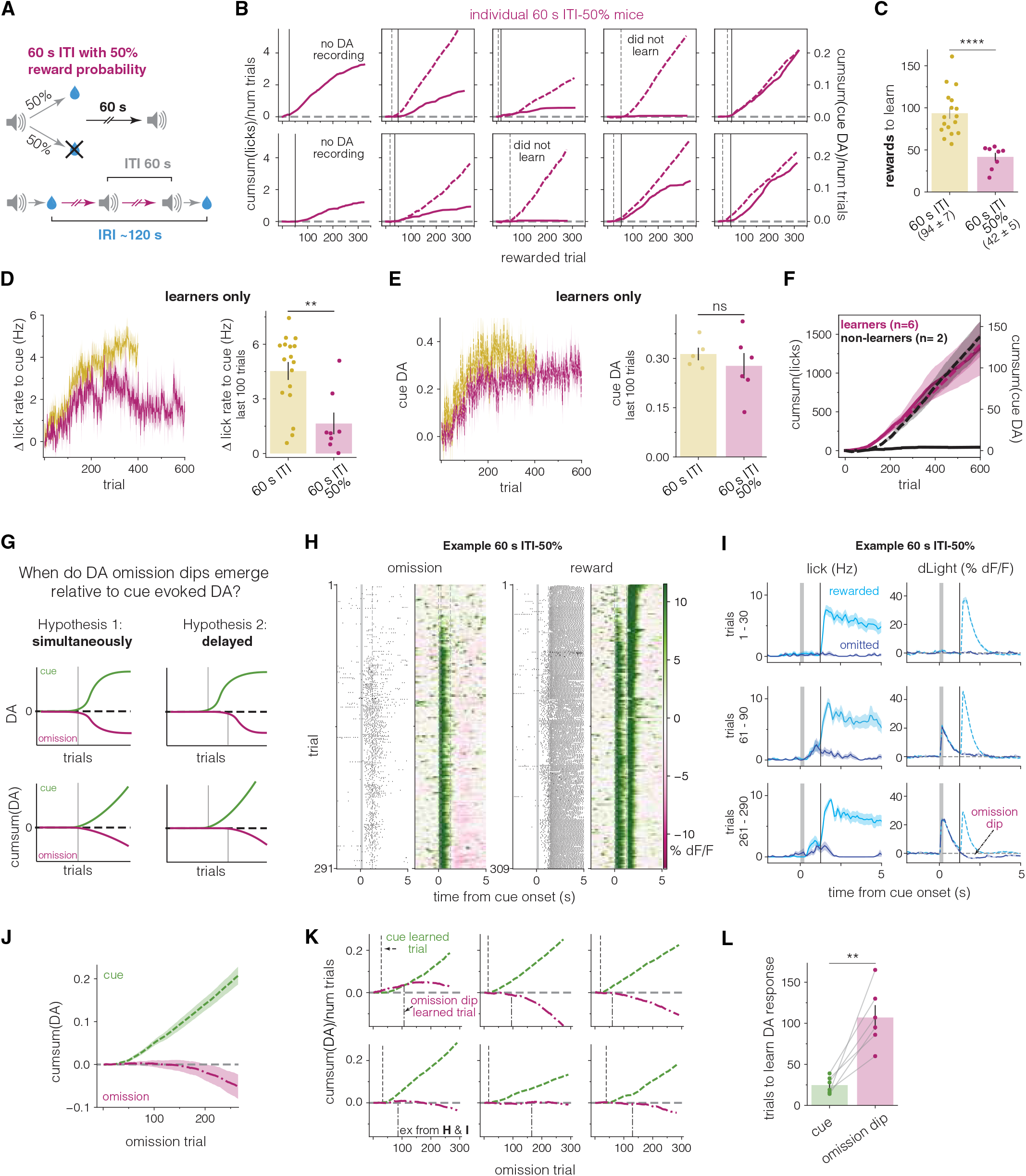
Partial reinforcement scales learning rate by increasing the inter-reward interval. **A**, Schematic of 60 s ITI-50% partial reinforcement conditioning. Mice were conditioned identically to 60 s ITI mice (**Fig. 1**) except rewards were delivered with 50% reward probability for 50 trials with ∼25 rewards a day. Reducing the reward probability by 50% leads to a doubling of the IRI to ∼120 s on average across a session while maintaining the ITI and ICI. Mice were conditioned for 12 days. **B**, Cumsum of cue-evoked licks (solid, left axis) or normalized cue-evoked dopamine (dashed, right axis) as a function of rewarded trials for all 60 s ITI-50% mice. Data presented as in **Fig. 2D**. Dopamine was not recorded from two initial mice and two other mice did not show evidence of behavioral learning (see *Methods*). **C**, 60 s ITI-50% mice (magenta, n = 8) learn the cue-reward association in about half the number of rewards as 60 s ITI mice (same data as **Fig. 1F**). Bar height represents mean number of rewards after which mice show evidence of learning.. Values under labels represent mean ± SEM. Mice that did not show evidence of learning (**B**) were excluded from analysis. **** p<0.0001, Welch’s t-test. **D-E**, Among mice that learned the cue-reward association, 60 s ITI-50% mice show lower asymptotic lick rates than 60 s ITI mice, consistent with prior literature on partial reinforcement; however, asymptotic dopamine responses do not show statistically significant difference between groups suggesting a dissociation between behavior and dopamine. *Left*, Timecourse of cue-evoked lick rate (**D)** or normalized cue-evoked dopamine (**E)** across all conditioning trials for 60 s ITI-50% (magenta, n = 8 behavior, n = 6 dopamine) and 60 s ITI (gold, n = 17 behavior, 5 dopamine) mice. *Right*, Mean cue-evoked lick rate (**D)** and normalized cue-evoked dopamine (**E)** during the last 100 trials of conditioning for both groups. **p<0.01, ns: not significant, Welch’s t-test **F**, Further suggesting a dissociation between behavior and dopamine, non-learner mice (red, n = 2) show intact dopaminergic learning similar to behavioral learner mice (black, n = 6). Cumsum of cue-evoked licks (left axis, solid lines) or normalized cue-evoked dopamine (right axis, dashed lines) over all 600 trials. **G**, Diagrams of two possible relationships between the onset of cue-evoked dopamine and the emergence of reward omission driven dopamine dips over the course of learning. **H**, Lick raster and heatmap of dopamine aligned to cue onset for one example 60 s ITI-50% mouse during omission (left) and rewarded (right) trials across conditioning. **I**, Lick (*left*) and dopamine (*right*) PSTHs aligned to cue onset for the same example mouse as in **H**. Light blue represents data from trials where rewards were delivered, while dark blue represents trials where rewards were omitted. Data are binned into early conditioning (trials 1-30, *top*), middle conditioning (trials 61-90, *middle*), and late conditioning (trials 261-290, *bottom*). In the middle row, note the prominent cue-evoked dopamine and licking and the absence of a dopamine dip following reward omission. **J**, On average, reward omission driven dopamine dips emerge later than cue-evoked dopamine in 60 s ITI-50% mice (n = 6). Average cumsum of normalized dopamine responses to cue presentation (green, dashed) or reward omission (magenta, dashdot) across reward omission trials. **K**, Omission dips emerge later than cue-evoked dopamine in individual mice. Cumsum of normalized dopamine responses to cue presentation (green, dashed) or reward omission (magenta, dashdot) across omission trials for all 60 s ITI-50% mice with dopamine recordings that learned the cue-reward association. Dashed lines on upper half of plots represent the omission trial after which cue-evoked dopamine emerges (dopamine learned trial), and dashdot lines on bottom of plot represent the omission trial after which dips in dopamine following reward omission emerge. **L**, On average, cue-evoked dopamine emerges 82 omission trials before dips in dopamine following reward omission. Bar height represents mean number of omissions before cue-evoked dopamine (green) or reward omission driven dips in dopamine (magenta) begin. Error bar represents SEM. Circles represent individual mice, and lines connect data from a single mouse. ** p <0.01, paired t-test.

In addition, while two out of ten total 60 s ITI-50% mice did not learn the cue-reward association, of the mice that did learn, cue-evoked lick rate at the end of conditioning was less than half that of 60 s ITI mice (60 s: 4.5 ± 0.5 Hz, 60s-50%: 1.65 ± 0.6 Hz, p<0.01; **Fig. 6D**), consistent with prior literature on partial reinforcement (64). However, cue-evoked dopamine was not significantly different between 60 s ITI-50% and 60 s ITI mice (60 s: 0.31 ± 0.02, 60s-50%: 0.28 ± 0.04 Hz, p = 0.43; **Fig. 6E**), demonstrating a dissociation between dopamine and behavior. Further suggesting a dissociation between dopamine and behavior, non-learner 60 s ITI-50% mice develop cue-evoked dopamine responses, consistent with the emergence of behavior (but not dopamine) requiring a threshold crossing (**Fig. 6F**).

We next used our dataset to further disambiguate between ANCCR and canonical TDRPE based on divergent predictions of the timing of omission dips in dopamine signaling relative to the onset of cue-evoked dopamine. Here we test between two hypotheses: Hypothesis 1 is based on the canonical TDRL framework of dopamine signaling in which a positive RPE at cue implies a positive cue value, and a positive cue value implies a negative RPE upon omission of the reward (this holds even if the omission dip is smaller than the reward burst). Thus, positive cue response and negative omission response co-evolves over trials despite differences in kinetics or magnitude. Hypothesis 2 is based on the ANCCR framework, which assumes that a reward omission becomes a meaningful event only after the cue-reward association exceeds some internal threshold (41). Thus, ANCCR predicts that the suppression of dopamine response to reward omission should occur well after the emergence of the dopamine cue response (**Fig. 6G**). When examining data from individual animals, there is a clear distinction in the onset of cue-evoked dopamine and the delayed onset of dopamine dips in response to reward omissions (**Fig. 6, H and I**) that can be easily visualized in the mean cumulative sum plots of the dopamine responses across animals (**Fig. 6J**). Using the cumulative sum of individual animals to determine the omission trial at which cue-evoked dopamine and omission dips emerge, we find that cue-evoked dopamine emerges after 25±4 omission trials. Dopamine dips, on the other hand, do not appear until after over 100 omission trials (107 ± 15, p<0.01, **Fig. 6, K and L**). In principle, modifications of TDRPE in which temporal expectation of reward is learned after initial cuereward learning (similar to ANCCR), may account for these results. While these results constrain development of future TDRPE models, they are naturally consistent with ANCCR.

## Discussion

We demonstrate that the rate of behavioral and dopaminergic cue-reward learning scales proportionally with the duration between reward experiences. Furthermore, partial reinforcement effectively extends the IRI and counterintuitively increases learning rate per reward experience. We show that the mathematical rule underlying learning rate control is a natural consequence of retrospective learning triggered by rewards. These results require a reevaluation of the frameworks used to describe associative learning and a reassessment of the implicit assumption that the trial is the fundamental unit of experience (18).

While standard TDRL implementations that ignore the ITI cannot, by definition, explain IRI effects on learning rate, we demonstrate that an extension of TDRL that ascribes lower ITI value during longer ITI is also unable to account for the experimentally observed proportional scaling of dopaminergic and behavioral learning. In principle, our observations may be semantically consistent with a scaling of learning rate by salience. Nevertheless, we do not consider this construct as an “explanation” of these results. The reason is that as a singular construct, salience is typically argued to be mediated by many variables, including perceptual intensity, novelty, and overall rate of stimuli. We demonstrate that controlling for these variables does not abolish the increase in learning rate with longer IRI. Thus, to explain our results, one would need to specifically redefine salience as being determined only by IRI. If so, we believe that IRI is the fundamental variable of relevance, and the construct of salience does not have additional explanatory power. Along similar lines, one could argue that a version of TDRL with learning rate scaled by IRI could explain our data. While such a model approximately fits the behavioral learning rate across IRIs (**fig. S7D**), it is not an “explanation” per se, because there is no known principle by which such a scaling rule can be derived from the core principle in TDRL of recursive value estimation. Further, identity-specific IRIs are not commonly computed in TDRL. On the other hand, this identity-specific IRI scaling rule is a natural consequence of retrospective learning in ANCCR because the Bayes rule conversion of a retrospective to prospective association (the core principle in ANCCR) requires timescale matching of retrospective cue-reward learning with the learning of overall rates for each event type. In sum, we believe that an explanation of our observations requires a model that derives a scaling rule of learning rate with IRI from first principles. In the absence of current alternatives, ANCCR provides a parsimonious explanation of these findings. One potential concern for the generality of the retrospective learning mechanism in ANCCR might be latent inhibition of behavioral learning. Specifically, animals experiencing cues in the absence of rewards learn subsequent cue-reward associations slower (65). Superficially, this may seem inconsistent with retrospective learning driven purely by rewards. However, due to the blocked nature of the latent inhibition experiments (cues are experienced without rewards before cue-reward experiences), ANCCR naturally explains latent inhibition. This is because the baseline cue rate, which is tracked in the absence of rewards, is learned to be high in the first phase of the experiments. This means that more cue-reward experiences are then needed to learn that cues preferentially precede rewards, thereby slowing learning (**fig. S11H**). Collectively, we believe that the retrospective learning mechanism in ANCCR is a general property of cue-reward learning.

One possibility by which “trial-based” frameworks of learning could be used to account for the data presented here is to assume that replays/reactivations of cue-reward experiences during the extended ITI or sleep (66–73) provides “virtual trials” in lieu of the real trials experienced by mice (63). Based on this modification, one can argue that during long ITI periods, the animals constantly replay the real cue-reward experiences, thereby making up for the absence of real experiences with imagined ones. As recognized previously (63), this means that a single real experience should in principle be sufficient to produce a never-ending stream of replays. It has been suggested that this does not happen in practice because other competing experiences, especially those occurring more frequently, would themselves engage replay, thereby drowning out the impact of the single experience. The “60 s ITI with CS-“ experiment (**Fig. 4G**) tests this idea, since the gap between CS+-reward experiences is filled with other competing CS- -omission experiences occurring 9 times more frequently. The fact that these competing experiences do not affect the learning of CS+-reward experiences is challenging to the replay idea. Instead, if the replay idea is modified to preferentially replay only experiences terminating in rewards (as opposed to omissions) (74), the learning from this form of replay is similar to the retrospective learning proposed by ANCCR, because only the causes of meaningful events such as rewards are replayed and learned. Thus, replays/reactivations may be a mechanistic implementation of retrospective learning.

Here, we primarily focused on dopamine-based alternative theories of learning. However, some psychological/ algorithmic theories are applicable to the behavioral learning observed here. These include the SOP theory (59, 60), the analytic theory of learning (TATAL (75)), and the Laplace transform theory of learning (76). Of these, we have shown that while SOP captures a qualitative increase in learning rate by ITI, it does not explain proportional scaling of learning rate by IRI. The behavioral learning data are generally consistent with TATAL and the timescale invariant Laplace transform theory of learning. However, the observation that 50% partial reinforcement produces learning in half as many rewards is inconsistent with TATAL (75) and has yet to be tested for the Laplace transform theory. Additionally, neither theory has yet been proposed as a general model of dopamine signaling. Thus, we believe that ANCCR currently provides the most parsimonious account of behavioral and dopaminergic learning. Further, the core principle of multi-timescale learning in these models, which is consistent with some experimental data (77–81), may reflect a spectrum of baseline learning rates (αo) and event rate time constants in ANCCR. While recent studies have provided accumulating evidence that mesolimbic dopamine does not function strictly as a TDRPE signal (37, 38, 41–43, 57, 82–84), the current results call into question the broader trial-based reinforcement learning framework used to understand dopamine and learning. Collectively, we provide a new framework for understanding dopamine mediated cue-reward learning (41) that explains the proportionality between learning rate and IRI in both dopaminergic and behavioral learning. As retrospective learning updates occur at every reward, our model does not rely on the concept of an experimenter-defined trial, which leads to potentially problematic assumptions (85). Thus, it is uniquely suited to account for experience outside an arbitrarily imposed “trial period”. In this way, the ANCCR theory gets us closer to explaining naturalistic learning outside laboratories where experiences do not have defined trial structures. By providing a framework for understanding the data presented here, grounded in the known dynamics of dopamine-mediated learning (41), our results provide further support for a reevaluation of the neural algorithms underlying associative learning.

## ACKNOWLEDGEMENTS

We thank J. Berke, L. Frank, M. Kheirbek, D. Ron, K. Bender, D. Canzio, G.D. Stuber, M. Andermann, S. Mihalas, I. Trujillo Pisanty, J. Rodriguez-Romaguera, R. Gowrishankar, A. Lempel, M. Duhne, A. Mohebi, M. Bocarsly, and members of the Namboodiri laboratory for helpful discussions. This project was supported by NIH R00MH118422, R01MH129582, R01AA029661, the Scott Alan Myers Endowed Professorship (V.M.K.N.), and NIH F32DA060044 (D.A.B.). The authors have no competing interests.

## AUTHOR CONTRIBUTIONS

D.A.B. and V.M.K.N. conceived the project. D.A.B., H.J., S.L., B.W., and J.R.F. performed experiments. D.A.B. performed analyses. A.T. and H.J. performed simulations. V.M.K.N. oversaw all aspects of the study. D.A.B. and V.M.K.N. wrote the manuscript with help from all authors.

## Methods

### Animals

All experiments and procedures were performed in accordance with guidelines from the National Institutes of Health Guide for the Care and Use of Laboratory Animals and approved by the UCSF Institutional Animal Care and Use Committee. 101 adult (>11 weeks at time of experiments; median: 13 weeks) wild-type male and female C57BL/6J mice (JAX; RRID:IMSR_JAX:000664) were used across ten experimental groups <u>30 s ITI</u> (n = 6; 3F/3M), <u>60 s ITI</u> (n = 19; 13 behavior-only 7F/6M & 6 DA+behavior: 4F/2M), <u>300 s ITI</u> (n = 6; 3F/3M), <u>600 s ITI</u> (n = 19; 12 behavior-only: 5F/7M & 7 DA+behavior: 5F/2M), <u>3600 s ITI</u> (n = 5; all DA+behavior: 3F/3M), <u>60 s ITI – few trials</u> (n = 18; 12 behavior only: 6F/6M & 6 DA+behavior: 3F/3M), <u>60 s ITI – few trials with context extinction</u> (n = 6; 3F/3M), <u>60 s ITI with CS-</u> (n = 6; 3F/3M), <u>600 s ITI with background milk</u> (n = 6; 3F/3M), and <u>60 s ITI-50%</u> (n = 10; 2 behavior-only: 0F/2M & 8 DA+behavior: 4F/4M). One 60 s ITI mouse implanted with an optic fiber was excluded from all dopamine analyses due to a missed fiber placement (**fig. S3**). One additional 60 s ITI mouse that was implanted with an optic fiber failed to learn the cue-reward association (**fig. S2C**) and was excluded from dopamine analyses. Two 60 s ITI-50% mice implanted with dopamine fibers failed to learn the cue-reward association (**Fig. 6B** and **fig. S11D**) and were excluded from comparisons against 60 s ITI mice and omission dip calculations (**Fig. 6B and fig. S11)**. While small sample sizes preclude a rigorous analysis of sex differences across all conditions tested, no significant difference in “trials to learn” was found between females and males in either 60 s or 600 s ITI groups, the two conditions best powered to detect differences. Thus, sexes were pooled for all analyses.

All mice were head-fixed during conditioning and underwent surgery prior to behavior experiments to either implant a custom head-ring for head-fixation (behavior-only) or to inject viral vector and implant an optic fiber and head-ring (DA+behavior) (See *Surgery*). Mice were > 7.5 weeks old at time of surgery (median: 9 weeks). Following surgery, mice were given at least a week to recover before beginning water deprivation. Mice implanted with optic fibers did not begin experiments until > 3.5 weeks following surgery to allow time for virus expression. During water deprivation, mice were given ad libitum access to food but were water deprived to ∼85 – 90% of pre-deprivation bodyweight and maintained in that weight range throughout experiments through daily adjustments to water allotment. Mice were weighed and monitored daily for the duration of deprivation. After surgery, mice with only a head ring implant were group housed in cages containing mice from multiple experimental groups, while fiber implanted mice were single housed. Mice were housed on a reverse 12-h light/dark cycle, and all behavior was run during the dark cycle.

### Surgery

Surgery was performed under aseptic conditions. Mice were anesthetized with isoflurane (5% induction, ∼1-2% throughout surgery) and placed in the stereotaxic device (Kopf Instruments) and kept warm with a heating pad. Prior to incision, mice were administered carprofen (5 mg/kg, SC) for pain relief, saline (0.3 mL, SC) to prevent dehydration, and local lidocaine (1 mg/kg, SC) to the scalp for local anesthesia. All mice were implanted with a custom-designed head ring (5 mm ID, 11 mm OD, 3 mm height) on the skull for head-fixation. The ring was secured to the skull with dental acrylic supported by screws. Following surgery, mice were given buprenorphine (0.1 mg/kg, SC) for pain relief.

To measure dopamine release in a subset of mice, 500 nL of an adeno-associated viral (AAV) vector encoding the dopamine sensor dLight1.3b (AAVDJ-CAG-dLight1.3b, 3.9 × 10^13^ GC/ml diluted in sterile saline to final titer of 3.9 × 10^12^ GC/ml) was injected unilaterally into NAc core (from bregma: AP 1.3, ML +/-1.4, DV -4.55), in either right or left hemisphere, counterbalanced across groups. Viral vectors were injected through a small glass pipette with a Nanoject III (Drummond Scientific) at a rate of 1 nL/s. Injection pipette was kept in place for 5-10 min to allow diffusion, then slowly retracted to prevent backflow up the injection tract. Following injection, an optic fiber (NA 0.66, 400µm, Doric Lenses) was implanted 200-350 µm above the virus injection site. Following fiber implant, the head ring was secured to skull as above. After experiments, fiber implanted mice were transcardially perfused, and brains were fixed in 4% paraformaldehyde. Brains were sectioned at 50 µm and imaged on a Keyence microscope to verify fiber placement. Histology images presented (**fig. S3**) represent composites imaged with a 10x objective. Stitched images of full brain slices were then cropped to focus on fiber placements in nucleus accumbens (**fig. S3B**) or dorsal striatum (**fig. S3C**).

### Conditioning

For experiments in **Figs. 1 to 3**, all animals were conditioned with an identical trial structure (see below), differing only in inter-trial interval (ITI) as well as number of trial presentations to keep total conditioning time roughly equal between groups (∼ one hour). **Figs. 1 and 2**: <u>60 s ITI</u> mice were run for 50 trials a day with a variable ITI with a mean of 60 s (uniformly distributed from 48 s to 72 s). <u>600 s ITI</u> mice were run for 6 trials a day with a variable ITI with a mean of 600 s (uniformly distributed from 480 s to 720 s). **Fig. 3:** <u>30 s ITI</u> mice were run for 100 trials a day with a variable ITI with a mean of 30 s (uniformly distributed from 24 s to 36 s). <u>300 s ITI</u> mice were run for 11 trials a day with a variable ITI with a mean of 300 s (uniformly distributed from 240 s to 360 s). <u>3600 s ITI</u> mice were run for 2 trials a day with fixed ITI of 3600 s and, unlike other groups session time lasted two hours.

For experiments in **Fig. 5**, additional groups of mice were conditioned with parameters matching those of 60 s and/or 600 s ITI groups to control for the influence of factors that varied along with ITI (IRI) manipulations. <u>60 s ITI - few trials</u> mice were run for 6 trials a day (same as 600 s ITI mice) with a mean ITI of 60 s (uniformly distributed from 48 s to 72 s; same as 60 s ITI) to control for the difference in total trial experiences per day between 60 s ITI and 600 s ITI mice. Unlike other groups, sessions lasted ∼6.5 minutes. <u>60 s ITI-few with context extinction</u> were conditioned similar to 60 s ITI-few mice but remained in the experimental context for ∼55 minutes following the end of conditioning trials, matching 600 s ITI group’s time in context and number of cue-reward experiences, while the rate of rewards during trials matched the 60 s ITI group. <u>60 s ITI with CS-</u> mice were conditioned similar to 600 s ITI mice, for 6 (CS+) trials a day with a variable (CS+) ITI with a mean of 600 s (uniformly distributed from 480 s to 720 s); however, during the interval between CS+ trials, distractor CS-cues (0.25 s, 3 kHz constant tone, delivered through a piezo speaker: https://www.adafruit.com/product/1739) were presented. CS-cues were not followed by reward delivery and were delivered on a variable interval (exponentially distributed) with a mean of 60 s to approximate the rate of cue delivery in 60 s ITI mice. All mice could hear and respond to the CS-cue as evidenced by some amount of generalized licking to the CS-over the course of conditioning (**fig. S10, C and D**). <u>600 s ITI with background chocolate milk</u> mice were conditioned similar to 600 s ITI mice but during the interval between cue-sucrose trials (mean of 600 s, uniformly distributed from 480 s to 720 s), mice received 2 un-cued deliveries of chocolate milk (Nesquik Low Fat Chocolate Milk) separated from the previous sucrose or chocolate milk delivery by a variable interval with a mean of 180 s (uniformly distributed from 144 s to 216 s) to test whether cue-sucrose learning rate is affected by the general or identity specific rate of rewards. Volume of chocolate milk was calibrated to match that of sucrose reward delivery (2-3 µL). Mice readily consumed chocolate milk rewards upon delivery (**fig. S10H**).

For experiments in **Fig. 6**, of <u>60 s ITI-50%</u> mice were conditioned identically to 60 s ITI mice (variable ITI with a mean of 60 s, uniformly distributed from 48 s to 72 s), except rewards were delivered with 50% reward probability for 50 trials with ∼25 rewards a day to disambiguate the (CS+) inter-cue interval (ICI) from the IRI. Reducing the reward probability by 50% leads to a doubling of the IRI to ∼120 s on average across a session while maintaining the ITI and ICI. Mice were conditioned for 12 days. Trials (CS+) consisted of a 0.25 s 12 kHz constant tone through a piezo speaker (https://www.adafruit.com/product/1740) followed by a 1 s delay (trace period) after which sucrose sweetened water (2-3 µL; 15% w/v) was delivered through a gravity fed solenoid to a lick spout in front of the mouse, controlled by custom Matlab and Arduino scripts (45). After each outcome, there was a fixed three second period to allow reward consumption. Lick spout was positioned close to the animals such that animals could sense, but were not touched by, delivery of reward. Licks were detected through a complete-the-circuit design and recorded in Matlab. Occasionally, certain mice would have long unbroken contacts with the spout (as measured by lick off – lick on time, due to grabbing the spout with their hands or not breaking contact with their tongues) occluding our ability to measure multiple licks during the period of contact. This was not corrected for as it generally happened following reward delivery or during the ITI and thus did not affect measurements of cue-evoked licks, our main variable of interest.

Mice were not habituated to the head-fixation apparatus or sucrose delivery prior to conditioning to minimize uncued reward exposure, which, we hypothesize, could affect retrospective contingency calculations during initial cue-reward learning. For the majority of mice, the first trial was their first experience of liquid sucrose reward. An initial subset of behavior only 600 s ITI mice (n = 6) ran with a fixed ITI of 600 s and was given a single uncued reward delivery prior to conditioning on day 1. No gross difference in learning compared to subsequent groups was detected, and data were pooled. For all other groups on day 1, mice were placed in the head-fixation apparatus and conditioning commenced. Because a minority of animals from each condition did not initially consume sucrose at time of reward delivery, for all analysis, “trial 1” was defined as the first trial in which a mouse licked to consume sucrose within 5 seconds of reward delivery. This design choice did not affect our main conclusions as analyzing “trials to learn” in 30 s – 3600 s ITI mice without dropping any initial trials from analysis shifted the mean learned trial by <1 trial.

Mice were run for at least 8 days of conditioning, and trial analyses included the first 800 (30 s ITI), 400 trials (60 s ITI), 80 (300 s ITI) 40 (600 s ITI, 60 s ITI-few, 60 s ITI-few with context extinction, 60 s ITI with CS-, 600 s ITI with background chocolate milk), or 7-8 trials (3600 s ITI). For 60 s ITI-50% mice, trial analyses included the first 600 trials. For rewarded or omission trial specific analyses, all reward or omission trials occurring within the first 600 trials were analyzed (∼300).

### Fiber photometry

Fluorescent dLight signals were collected using either a Doric Fiber Photometry Console or pyPhotometry (86) system. For both systems, light from 470 nm and 405 nm LEDs integrated into a fluorescence filter minicube (Doric Lenses) was passed through a low-autofluorescence patchcord (400 µm, 0.57 NA, Doric Lenses) to the mouse. Emission light was collected through the same patchcord, bandpass filtered through the minicube, and measured with a single integrated detector. For the Doric system, excitation LED output was sinusoidally modulated by a Doric Fiber Photometry Console running Neuroscience Studio v5.4 or v6.4 at 530 hz (470) and 209 hz (405). The console demodulated the incoming detector signal producing separate emission signals for 470 nm excitation (dopamine) and 405 nm excitation (dopamine-insensitive isosbestic control). Signals were sampled at 12 kHz and subsequently downsampled to 120 Hz following low pass filtering at 12 Hz. For the pyPhotometry system, excitation 405 nm and 470 nm LEDs were modulated in time, rather than frequency, with separate brief 0.75 ms pulses used to separate isosbestic and signal channels. Data were sampled at 130 Hz and low pass filtered at 12 Hz to match data from Doric system. Due to a software error during Doric system file save, the final trial was not recorded on two occasions (one 60 s ITI, one 600 s ITI) and was excluded from analysis. This error occurred either well before (60 s ITI) or well after (600 s ITI) the emergence of learning, and thus had minimal effect on the resulting analysis. For one 3600 s ITI animal, a pyPhotometry system crash during the ITI between trials 1 and 2 on day 5 resulted in ∼15 minutes of photometry data loss during the ITI but did not affect analysis focused on cue and reward delivery epoch. A TTL pulse signaling behavior session start and stop was recorded by the photometry software to sync and align photometry and behavior data across hardware.

### Analysis

#### Behavior

The behavioral measure of learning here was licking in response to the cue before reward delivery. As mice learn the cue-reward association, cue presentation elicits anticipatory licking behavior toward the reward spout. To measure the cue-evoked change in licking behavior over baseline, the number of licks in the 1.25 s baseline period before cue onset was subtracted from the number of licks in the 1.25 s period from cue onset to reward delivery to calculate the change in licking behavior to the cue (cue-evoked licks). When this number was converted to a rate, it was reported as “Δ lick rate to cue.” To binarize cue-evoked licking behavior, we also measured the proportion of mice in each group that made more than one cue-evoked lick on each trial across conditioning (**Figs. S1 and S2**). To visualize average trial licking behavior for each session in example animal plots (**Figs. 1C, 2C, 3, B and J, and 6I**, and **figs. S1A and S4A**) or reward delivery aligned group averages (**figs. S9F and 10H**), lick peri-stimulus time histograms were generated by binning licks into 0.1 s bins, converting to a rate, and averaging across trials. The resulting average lick rate trace was smoothed with a Gaussian filter to aid visualization.

To calculate the trial at which animals show evidence of learning, we first took the cumulative sum (cumsum) of the cue-evoked licks (17, 41, 47, 48, 50). Then drawing a diagonal from beginning to the end of the cumsum curve, we calculated the first trial that occurred within 75% of the maximum distance from the curve to the diagonal, which corresponded to the trial after which cue-evoked licking behavior emerged (**fig. S2, A to C**). This trial was designated the “learned trial.” Occasionally after learning, cue licks taper off. If at the calculated learned trial the diagonal line was underneath the cumsum curve, which means that the mouse’s lick behavior was decreasing at that point rather than increasing, we iteratively reran the algorithm by drawing the diagonal from the beginning to the point on the cumsum curve corresponding to the previously calculated trial until at the new calculated trial the diagonal was above the cumsum curve (corresponding to the trial in which lick behavior begins to increase). Note that we use the first trial within 75% of maximum distance rather than the overall maximum distance (which would be the largest inflection point in the curve) to account for variability in post-learning behavior that occasionally caused the maximum distance from the diagonal to be at a point after a mouse has consistently licked to the cue for many trials; however, this choice did not affect the main conclusion of the analysis in **Fig. 1** that 600 s ITI mice learn in ten times fewer trials than 60 s ITI mice (**fig. S2D**). Mice that did not show a > 0.5 Hz average increase in lick rate to cue for at least 2 sessions were classified as non-learners (**Fig. 6B** and **figs. S2C, S5C, S6A, and S11D**) and were not considered in comparisons of learned trials (**Figs. 1F, 3E, and 6C**, and **figs. S9, H to K, and S11E**). To measure the steepness of individual animal learning curves, we calculated the abruptness of change at the learned trial as the distance from the cumsum curve to the diagonal described above. This distance was calculated in normalized units where the top of the diagonal was set to equal 1 (**fig. S2H**). Cumsum data are occasionally displayed divided by the number of trials (yielding a y-axis that corresponds to average response across all prior conditioning trials) to better compare across groups that experienced different numbers of trials.

To quantify the relationship between learning rate and inter-reward interval (IRI; **Fig. 3H**), the mean trials to learn for 30 s, 60 s, 300 s, and 600 s ITI groups were plotted against the IRI (mean ITI + 4.25 s [1.25 s trial period + 3 s consummatory period]) on a log-log plot, and a linear least squares regression was used to determine the best fit line yielding the equation: *log*(trials_to_learn) = (−1.0593)*log*(IRI) + 3.8753. The slope and intercept determined here were used use to calculate the predicted trials or rewards to learn for 3600 s ITI (3604.25 s IRI); **Fig. 3, M and N**), 600 s ITI with background milk with a general IRI (604.25 s) or an identity specific IRI (204.25 s; **fig. S10F**), and 60 s ITI-50% as predicted by the ICI (64.25 s) or IRI (128.5 s; **fig. S11A**).

For analysis of ITI lick rates (**fig. S9, G to K**), lick rate was calculated from the period beginning with either the start of the session or the end of the prior consumption bout (consumption bout defined as the period from the first lick following reward delivery through all licks in which the interval between consecutive “lick off” to “lick on” was 500 ms) and ending with the onset of the following cue. ITI lick rate before learned trial was calculated as the mean of the ITI lick rate for every ITI preceding the animal’s learned trial. Animals that did not show evidence of learning were excluded from this analysis. One additional 600 s ITI mouse with many long (>10 s) contacts with the spout during the ITI across days (presumed to be due to holding the spout) was also excluded from this analysis.

#### Dopamine

To analyze the signals, a session-wide dF/F was calculated by applying a least-squares linear fit to the 405 nm signal to scale and align it to the 470 nm signal. The resulting fitted 405 nm signal was then used to normalize the 470 nm signal. Thus, dF/F is defined as 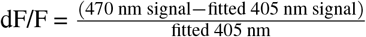, expressed as a percent (87). Cue-evoked dopamine (cue DA) was measured as the area under the curve (AUC) of the dopamine signal for 0.5 s following cue onset minus the AUC of the baseline period 0.5 s directly preceding cue onset. Reward evoked dopamine (reward DA) was measured as the AUC 0.5 s following the first detected lick after reward delivery minus the AUC of the pre-cue baseline period described above. If the onset and offset of a detected lick spanned reward delivery time, the reward AUC was calculated from time of reward delivery. For quantifying dopamine dips in response to omitted rewards (**Fig. 6, J to L**, and **fig. S11G)**, the AUC of a 2 s baseline window was subtracted from the AUC of 2 s window beginning 1.25 s following cue onset (time of reward delivery in rewarded trials). A longer duration window was used to measure dips to account for the slower kinetics and broader shape of dips relative to cue responses. All dopamine responses reported in main figures are AUC measurements, but peak measurements are also plotted as a comparison point (**figs. S4 and S6**). To measure cue and reward peak dopamine responses, the mean dopamine signal during the baseline period was subtracted from the maximum value of the dopamine signal during the cue and reward windows described above for AUC measurements. Similar to AUC measurements, peak responses were also normalized to the mean of the max 3 reward responses in each animal.

To facilitate comparisons across mice with differing levels of virus expression, cue and reward dopamine measurements per mouse were normalized to the average of the three maximum reward responses in that mouse. For omission responses, dopamine measurements were normalized to the individual animal average of the 3 maximum reward responses recorded in a 2 s window following reward delivery to match the window used for dip measurements. All presented dopamine values represent these individual maximum reward normalized measurements, aside from example mice in which dopamine is plotted as % dF/F and cumsum plots in **Fig. 2C** and **fig. S2C** in which data are normalized to the maximum value of each animal’s cumsum curve as described below. Maximum rather than initial reward responses were chosen, as the reward response initially increased across early conditioning trials with different numbers of trials until maximum between conditions (**figs. S4, G and H, and S6, H and K**).

To calculate the trial at which dopamine responses to the cue develop (DA learned trial), we took the cumsum of the normalized cue DA response described. A diagonal was drawn from trial 1 through the point on the cumsum curve at 1.5 times the behavior learned trial to account for decreasing cue responses with extended training (88). The same algorithm described above to determine the behavior learned trial was run on the cue DA curve. The lag between DA and behavioral learned trial (the number of trials between the development of dopamine responses to the cue and the emergence of behavioral learning) was defined as the behavior learned trial minus the DA learned trial (**Fig. 2F**). Omission dip learned trials (**Fig. 6, K and L)** were calculated using the same algorithm on omission dip responses to detect the negative going inflection point across all omission trials.

For one 60 s ITI dLight animal, during an initial conditioning session, a software crash caused the loss of lick data for 50 trials experienced by the animal. An additional 13 trials were presented to the animal that day and recorded following the crash. Photometry data were recorded for all 63 trials. Because the crash occurred prior to the emergence of learning and cue-evoked licking behavior (as confirmed by both online observation by experimenter prior to crash and a -0.14 Hz average cue-evoked change in lick rate for the 13 trials recorded after crash), the 50 trials in which data were lost were coded as 0 cue-evoked licks. All 63 trials the animal experienced were included in analyses.

To visualize the average relationship between DA responses and licking behavior across learning in 60 s and 600 s ITI mice with variability in individual learning rates, signals were aligned to behavior learned trial and plotted through 250 or 25 trials after learning (**Fig. 2C** and **fig. S2C**). For aligned cumsum plots, data were normalized by the value from trial 400 (60 s ITI) or trial 40 (600 s ITI).

To quantify the relationship between dopaminergic learning rate and IRI (**fig. S5E**), the mean dopamine trials to learn 60 s and 600 s ITI groups were plotted against the IRI (mean ITI + 4.25 s [1.25 s trial period + 3 s consummatory period]) on a log-log plot, and line between means was calculated as was done for behavior yielding the equation: *log*(trials_to_learn_DA) = (−1.0260)*log*(IRI) + 3.4062. The slope and intercept determined here were used use to calculate the predicted trials or rewards to learn dopamine for 3600 s ITI (3604.25 s IRI; **fig. S6D**), and 60 s ITI-50% as predicted by the ICI (64.25 s) or IRI (128.5 s; **fig. S11B**).

### Theory and Simulations

We previously proposed a new learning model named ANCCR based on the learning of retrospective associations (41). ANCCR operates by identifying cues consistently preceding meaningful events such as rewards. Thus, it learns whether a cue consistently precedes a reward (i.e., a retrospective cue-reward association). This retrospective association provides a means to estimate whether the reward consistently follows a cue (i.e., the prospective cue-reward association). The core principle of ANCCR is that a cue-reward association is learned and cached as a retrospective predecessor representation (denoted by *M*_←*cr*_), and then converted to a prospective successor representation (denoted by *M*_→*cr*_) using a Bayes’ rule-like normalization: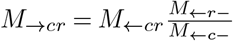. Here, *M*_←*r*−_ is proportional to the baseline rate of a reward in the environment, and *M*_←*c*−_ is proportional to the baseline rate of a cue. Here, we provide a quick intuitive derivation of the scaling of learning rate by the inter-reward interval (IRI). Please note that all the above variables are assumed to be conditioned on the experimental context.

Thus, they should be listed as *M* _←*cr*|*context*_, *M* _→*cr*| *context*_, *M* _←*c* −| *context*_, *M* _←*r*−|*context*_, Since listing these conditional dependences is notationally cumbersome, we omit this in our treatments.

Each term on the right hand side of the Bayes’ normalization (i.e., *M*_←*cr*_, *M*_←*r*−_, and *M*_←*c*−_) is updated using a delta-rule like update in ANCCR. Specifically,

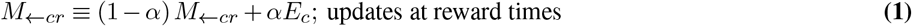

where *E*_*c*_ is the eligibility trace of the cue, *α* is the learning rate for the retrospective update, and ≡ is the symbol for update, and

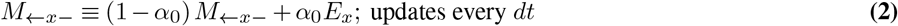

where *x* is any event type (e.g., cue or reward) and *E*_*x*_ is its eligibility trace. As can be seen, *M*_←*cr*_ updates at the time of every reward with a learning rate of *α*, and *M*_←*x*−_ updates every *dt* (i.e., continually) with a learning rate of *α*_0_. Both update rules determine a corresponding timescale of history for each variable (*M*_←*cr*_ or *M*_←*x*−_)—defined as the timescale over which a past event exerts influence on the current value of *M*_←*cr*_ or *M*_←*x*−_. The timescale over which one presentation of the cue influences future values of *M*_←*cr*_, i.e., its timescale of history, depends on both *α* and how frequently the reward occurs. On the other hand, the corresponding timescale for *M*_←*x*−_ depends on *α*_0_ and *dt*.

For the Bayes’ rule-like normalization to work in a (possibly) non-stationary environment, all terms on the right hand side (i.e., *M*_←*cr*_, *M*_←*r*−_, and *M*_←*c*−_) should be calculated over the same timescale of history. This is because normalizing the predecessor representation calculated over 1 hour (say) by baseline rates of reward and cue calculated over 1 min (say) is obviously inappropriate if the environment has the potential to change during the hour. Thus, the quantitative relationship between learning rate and IRI can be obtained by setting the timescale of history for *M*_←*cr*_, *M*_←*r*−_, and *M*_←*c*−_ to be equal to each other. As shown in Equation (2), the baseline rates of reward and cue are updated by a delta-rule with a baseline learning rate *α*_0_ continually (i.e., every timestep *dt*). Assuming that the time constant of decay of the eligibility trace is very short, a single occurrence of *x* will have a net influence on *M*_←*x*−_ of (1 − *α*_0_)^*n*^ after *n* timesteps—an exponentially decaying function of time. Equating this influence with an exponential time decay of 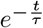 one can calculate the time constant of decay as as 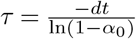. Thus, the net timescale of history for the calculation of the baseline rate of events (cues or rewards) is 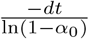, where the numerator is the time interval between consecutive delta-rule updates with learning rate *α*_0_. The timescale of history for the predecessor representation *M*_←*cr*_ has a similar expression with the numerator being the time interval between consecutive updates, which equals the IRI since updates only occur at reward times, and the learning rate in the denominator equals the learning rate of associative update, *α*. Thus, the timescale of history for the predecessor representation is 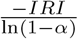. Setting both these timescales of history to be equal, one can show that the learning rate of associative update should be 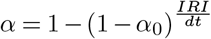. For small learning rates, this expression simplifies to 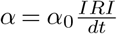.

For a more formal derivation that accounts for the time constant of eligibility trace, see **Supplementary Note 1**.

#### Comparison of learning models

To determine if a model of associative learning could capture the experimentally observed proportional scaling across ITI conditions, we simulated three likely candidates which all account in some way for the time between cue-reward trial experiences: the microstimulus implementation of temporal difference reinforcement learning (TDRL) (58), Wagner’s SOP (Sometimes Opponent Process or Standard Operating Procedure)(59, 60), and ANCCR (Adjusted Net Contingency for Causal Relations)(41). For each model, we simulated the experimental conditions for 30 s through 3600 s ITI and tested combinations of parameters to determine which could best replicate the quantitative, proportional scaling of learning rate by IRI observed experimentally. Our measure against which specific model instances were compared was the “trials to learn” for each of 30 s, 60 s, 300 s, 600 s, 3600 s ITI groups. Simulations of each model were based on published versions (41, 58, 59) with adjustments to ANCCR described above and below. To generate behavior, all models assumed that behavior emerged following a threshold crossing by the association quantity, which corresponded to cue “value” in both TDRL and SOP and net contingency (NC_cue↔reward_) in ANCCR. Thus “learned trial” is defined in TDRL and SOP as the first trial when value > threshold, and in ANCCR it is defined as the first trial where NC > threshold. While we recognize that action selection would likely involve other processes, the above threshold crossing was implemented to ensure that the models generated “learned trials” through comparable operations. For each combination of parameters for all three models, all five ITI conditions (30 s - 3600 s) were simulated for the number of trials that experimental animals experienced over 8 days of conditioning. Each case was iterated 20 times.

To determine the parameter combination from each model that best fit the experimental data, we calculated the residual sum of squares (RSS) of the trials to learn from each simulated parameter combination against the experimental trials to learn for each IRI (**Fig. 3H**). RSS was calculated on log transformed data to account for the wide variation in trials to learn across IRIs. The simulation with the lowest RSS was deemed the best fit experimental results.

After determining the best fit parameter combination for each model, TDRL, SOP, and ANCCR, we measured the Akaike information criterion (AIC) as AIC = 2*k* + *n* ∗*ln*(meanRSS), where n = sample size (number of animals), k = the number of parameters in the model, and meanRSS = the mean residual sum of squares between trials to learn for that parameter combination and experimental data. A lower AIC value represents a better fit to the data accounting for number of parameters needed to fit. Note that all data presented in figures (**Fig. 4** and **fig. S7**) and text and used to calculate model weights represent the AIC calculated in which models are not penalized for parameters by substituting k=0 into the equation above. This yields the formula AIC = *n*∗*ln*(meanRSS). This was implemented as a conservative measure because only a few parameters from most models had the potential to affect the simulated results and the best-fit model (ANCCR) is the one with the fewest parameters. For AIC values penalizing the total number of parameters per model, refer to **table S1**.

The best model between TDRL, SOP, and ANCCR was then determined as the one with the minimum AIC. A relative weight for each model compared to the best model was then calculated as modelweight 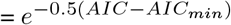. For additional simulation results presented in **figs. S7**, model weights were calculated using the best fit AIC from the other 2 models presented in **Fig. 4**.

#### TDRL simulations

TDRL assumes that animals assign value to each moment following an event (e.g., cue) to predict future reward. Each event elicits multiple states, and the value of each time step can be expressed as a weighted sum of activated states at that moment. If the prediction from the previous moment is different from what is experienced in the current moment, the model updates the value of the previous moment based on this reward prediction error, assumed to be signaled by dopamine. Depending on how the model represents a state, TDRL can be further divided into subtypes. Here, for a representative TDRL algorithm, we used the microstimulus model model (58) because it naturally accounts for the ITI. This model assumes that time states are Gaussian functions of increasing width following each event (cue or reward). The following model parameters were fixed for every iteration: bin size (dt) = 0.25 s (set to cue duration), decay parameter of eligibility trace (*λ*) = 0.99 (set to a high value to allow rapid credit assignment to earlier states), width of Gaussian function (*σ*) = 0.08. For the following parameters, we swept across a range to determine whether any combination could best explain proportional IRI scaling of learning rate: Threshold for behavior generation = [0.1, 0.2, 0.3, 0.4, 0.5, 0.6, 0.7, 0.8], decay parameter of event memory (*d*) = [0.8, 0.9, 0.99, 0.999, 0.9999], temporal discounting factor (*γ*) = [0.8, 0.9, 0.99, 0.999, 0.9999], number of states elicited by each event (*m*) = [3, 10, 100, 1000], learning rate (*α*) = [0.001, 0.01, 0.1]. The best-fit-to-behavior parameter combination (**Fig. 4D**) was: threshold = 0.3, *α* = 0.1, *γ* = 0.99, *m* = 3, and *d* = 0.9.

For comparisons between models (**Fig. 4**), simulations were run for the same number of trials as experimental groups (30 s: 800, 60 s: 400, 300 s: 88, 600 s: 48, 3600 s: 16). We again searched for the best-fit parameter combination (**fig. S7C**) when ITI conditions 60 s – 3600 were run for at least 400 trials. The best fit TDRL parameter combination when each ITI consisted of at least 400 trials per group was: threshold = 0.4, *α* = 0.10, *γ* = 0.80, *m* = 3, *d* = 0.999. AIC and model weight comparisons for this model were run against the best fit SOP and ANCCR models (from **Fig. 4, H and L**)

In principle, a similar rule derived in ANCCR could be applied *ad hoc* to any model of associative learning. Here, we demonstrate that applying such a rule to TDRL improves fit to experimental results. Using the best fit model parameters determined during initial TDRL parameter sweep described above (**Fig. 4, D to F**), we replaced the learning rate, *α*, based on the equation *α* = 1−*e*^(−*k*∗*IRI*)^ and performed another parameter sweep to determine the best fit *k*. We searched over the range *k* = [0.00015, 0.0002, 0.00025, 0.0003, 0.00035, 0.0004] (the range matching experimentally observed learning rate) and found that the best fit to behavior results from *k* = 0.0003 (**fig. S7D**).

#### SOP simulations

In SOP, cues or rewards evoke processing nodes consisting of many elements. These stimulus representations are dynamic: presentation of a stimulus moves a portion of elements from (only) the inactive (I) state into the primary active state (A1). Elements then decay into the secondary active state (A2) (a refractory state) and then decay again back to the inactive state while the stimulus is absent. Elements transition between states according to individually specified probabilities. Cue-reward associations (value) are strengthened and learned when cue elements in A1cue and reward elements in A1reward overlap in time and decreased when cue elements in A1cue and reward elements in A2reward overlap in time. Following learning, cues evoke conditioned responding by directly activating reward elements to their A2 state. One way in which SOP has been hypothesized to explain ITI impact on learning is that more time between trials allows more elements to decay to the inactive state (as opposed to the refractory A2 state), allowing for greater number of elements to transition to the A1 active state upon next cue and reward presentation. Parameter combinations were swept through to determine if any set of parameters could capture the quantitative scaling observed in the experimental results.

The relevant parameters in the model controlling the transition probabilities from I →A1 →A2→ I are *p*1, *pd*1, and *pd*2 respectively. *p*1_*cs*_, *pd*1_*cs*_, *pd*2_*cs*_ refer to the transition probabilities controlling the cue representation, while *p*1_*us*_, *pd*1_*us*_, and *pd*2_*us*_ refer to the transition probabilities controlling transitions between reward representation active states. The following parameters were fixed for every iteration of SOP run: the timestep (*dt*) = 0.25 s (set to cue duration), reward magnitude of CS in A1 (*r*1) = 1, reward magnitude of CS in A2 (*r*2) = 0.5, scale factor for magnitude of activation for coincidence of CS and US in A1(*L*_*plus*_) = 0.2, scale factor for magnitude of inhibition for coincidence of CS in A1 and US in A2(*L*_*minus*_) = 0.1, *p*1_*cs*_ = 0.1, and *p*1_*us*_ = 0.6 based on previously published work (59). The following parameter combinations reflecting the variables hypothesized to drive trial spacing effect were varied: threshold for behavior generation = [0.01, 0.1, 0.2, 0.3, 0.4, 0.5, 0.6, 0.7, 0.8], *pd*1_*us*_= [0.01, 0.2, 0.25, 0.5, 0.75], *pd*2_*us*_ = [0.001, 0.001, 0.01, 0.1], *pd*1_*cs*_ = [0.01, 0.2, 0.25, 0.5, 0.75], *pd*2_*cs*_ = [0.001, 0.001, 0.01, 0.1]. Because SOP implementations make the assumption that *pd*1 > *pd*2 (59) (i.e. the decay from the A1 active state to the A2 state should be faster than the decay from the A2 active state to the inactive state), we constrained our results to parameter combinations that satisfied this inequality. The parameter combination providing the best fit to behavior (**Fig. 4H**) was: threshold = 0.1, *pd*1_*us*_ = 0.25, *pd*2_*us*_ = 0.1 *pd*1_*cs*_ = 0.1, *pd*2_*cs*_ = 0.0001.

#### ANCCR simulations

In ANCCR, we derived a scaling rule for the retrospective learning rate (*α*) and the eligibility trace time constant (*T*) from the core principle of Bayes rule conversion of a retrospective to a prospective association (**Supplementary Note 1**). For the simulations considered here, these rules simplify to: 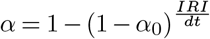 and *T* = *k*∗ *IRI*. For ANCCR, the parameters that were swept to identify the best-fit model were: threshold = [0.1, 0.2, 0.3, 0.4, 0.5, 0.6, 0.7, 0.8], *α*_0_ = [1×10^−4^, 8 ×10^−5^, 6 ×10^−5^, 4 ×10^−5^, 2 ×10^−5^, 1 ×10^−5^, 8 ×10^−6^], *k* = [0.1, 0.3, 0.5, 0.7]. The best-fit parameters were: threshold = 0.4, *α*_0_ = 4 ×10^−5^, *k* = 0.5. The following parameters were fixed: *w* = 0.5, *dt* = 0.2 (same as in (41). The dopamine response to the first reward was relatively high (though this response increases with repeated reward experience, consistent with our previous demonstration (41)). Two possibilities exist to account for this. One is that there is a Bayesian prior for *M*_*r*←*r*_ and *M*_*r*←-_, and the other is that part of the innate meaningfulness of a reward is signaled by dopamine. For simplicity, we assumed the latter and added an innate meaningfulness of 1 to dopamine reward response and 0 to dopamine cue response.

### Statistics

Statistical analyses were performed in Python 3.11 using either the scipy.stats (v1.11.4) or Pingouin (Vallat 2018) (v0.5.4) packages. Welch’s t-test was performed to compare between experimental groups, so as to not assume equal variances between the populations (**Fig. 1, F and I**). To test for equality of variances, F-tests were run using a custom script. Nonparametric tests (Kruskal–Wallis H, Mann–Whitney U) were used to compare simulation results due to the presence of conditions with 0 variance. For the eight experimental comparisons performed in **Fig. 5**, the false discovery rate was controlled using the Benjamini–Yakutieli method to adjust p-values. All other multiple comparisons were corrected for by adjusting p-values with Bonferroni’s correction (**Figs. 3R**, and **figs. S7, S9B, S10, B and F, and S11, A and B**). All statistical tests were two-tailed. N’s reported represent individual animals or, in the case of simulations, the number of iterations. All linear regressions presented fit with least-squares method using “scipy.stats.linregress” function (**Fig. 3H** and **figs. S5E, S7A, S9, I to K**). Full statistical test information is presented in **table S1** including test statistics, n’s, degrees of freedom, and both corrected and uncorrected p-values. Timecourses of the cumulative sum or average of the lick and/or dopamine data are presented as mean between animals/iterations ± SEM. Bar graphs are presented as mean between animals/iterations ± SEM with individual animal (or iteration) data points. Results were considered significant at an alpha of 0.05. * denotes p < 0.05, **p < 0.01, ***p < 0.001, ****p < 0.0001; ns (non-significant) denotes p > 0.05.

## Supplementary Figures

**Fig. S1.**
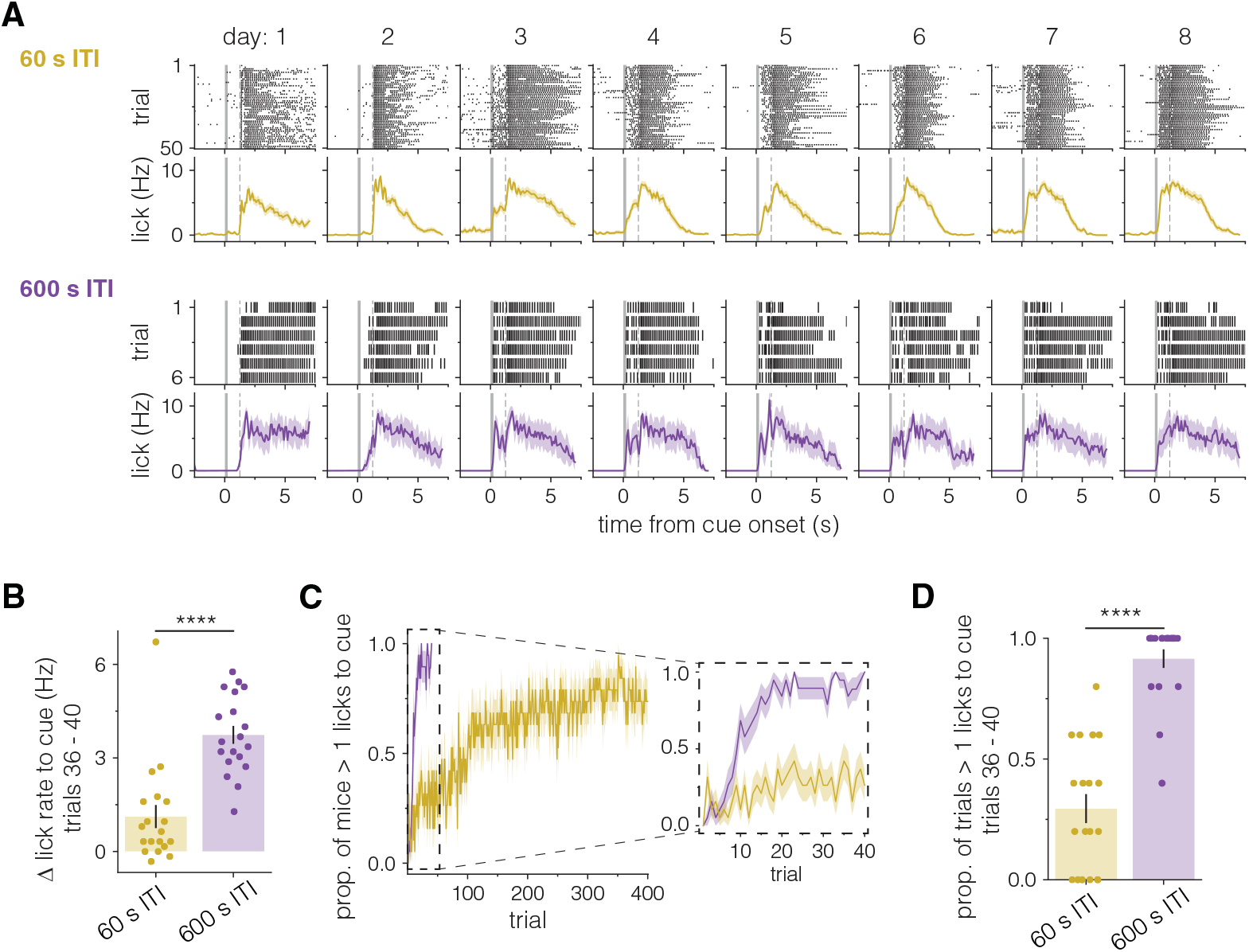
600 s ITI mice show evidence of learning in significantly fewer trials than 60 s ITI mice. **A**, Lick raster and PSTH plots from more individual 60 s ITI and 600 s ITI example mice, presented as in **Fig. 1C. B**, Average change in cue-evoked lick rate for trials 36 - 40 (timecourse in **Fig. 1D**). 600 s ITI mice show significantly more licking to cue in this period than 60 s ITI mice ****p< 0.0001, Welch’s t-test. **C-D**, 600 s ITI mice show asymptotic responding to the cue in fewer trials than 60 s ITI mice. **C**. *Left*, Timecourse showing the proportion of mice with more than one cue-evoked lick on each trial over 40 (600 s ITI, purple, n = 19) or 400 (60 s ITI, gold, n = 19) trials. *Inset, right*, Zoom in of first 40 trials for both groups. Lines represent mean across all animals and shaded area represents the SEM. **D**. Bar height represents proportion of trials in which animals responded to cue with more than one lick between trials 36 and 40. Error bars represent SEMs, and circles represent individual animals ****p< 0.0001, Welch’s t-test.

**Fig. S2.**
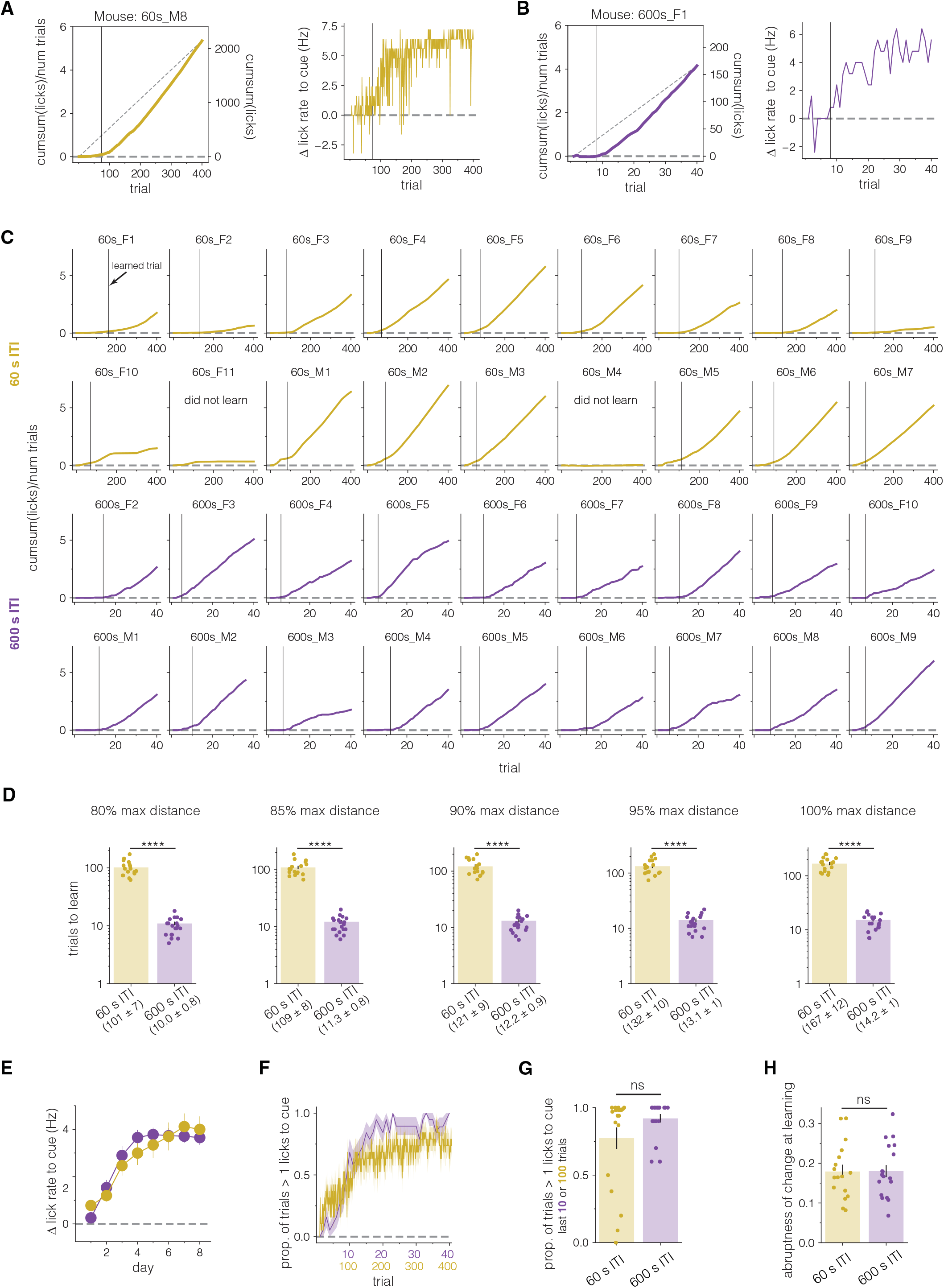
The cumulative sum of cue-evoked lick behavior allows for determination of learned trial in individual animals (A-D). Learning scales with total conditioning time or the ratio between inter-trial intervals (E-H). **A-B**, Cumsum and trial-by-trial plots of cue-evoked licking behavior of the same example mice as in **Fig. 1, C and E**. *Left*, Cumsum plot showing diagonal line used to calculate learned trials. Right y-axis shows the total cumsum of all cue-evoked licks as in **Fig. 1E**. Left y-axis shows cumsum of cue-evoked licks divided by total trial number to allow comparisons across groups that experienced a different number of cue-reward pairings. Trial normalized cumsum values represent the mean number of cue-evoked licks over all previous trials. Solid vertical line represents the calculated learned trial, the trial after which animals show evidence of learning. *Right*, The cue-evoked change in lick rate plotted for the same individual example mouse. Note how the vertical line representing the learned trial, which corresponds to the point on cumsum plots where the cumsum curve takes off from the x-axis, captures the trial at which cue-evoked change in lick rate becomes consistently positive. **C**, Cumsum plots for all remaining mice included in behavior analysis plotted with trial normalized units (see **A**-**B**). Vertical line represents calculated learned trial. Animals which did not meet learning criteria (see *Methods*) are noted, and no vertical line is drawn. These animals were excluded from comparison of learned trials between groups. **D**, For analysis, learned trial was calculated as the first trial that fell within 75% of the maximum distance from a diagonal drawn from the point on the cumsum curve at trial 1 through trial 40 or 400. 75% of maximum distance was chosen rather than the overall maximum distance (which would be the largest inflection point in the curve) to account for variability in post-learning behavior that occasionally caused the maximum distance from the diagonal to be at a point after a mouse has consistently licked to the cue for many trials. This choice did not affect our main conclusions as using 80%, 85%, 90%, 95%, or the maximum distance from the diagonal in our algorithm yielded a roughly similar result of ten times more trials needed to learn in 60 s ITI mice than 600 s ITI mice. Bar heights represent mean number of trials to learn, error bars represent SEMs, and circles represent individual animals plotted on a log scale. Values under labels represent mean ± SEM. ****p<0.0001, Welch’s t-test. **E**, Learning is similar between groups when trials are averaged across days. Timecourse of average cue-evoked lick rate as a function of days of conditioning. Circles represent mean change in lick rate per day, and error bars represent SEMs. **F - G**, Responding to cue scales with total conditioning time between groups and is not different at the end of conditioning. **F**. Timecourse showing the proportion of mice with more than one cue-evoked lick on each trial (same data as **fig. S1C**) plotted on scaled x-axis units. Lines represent means per group and shading represents SEM. **G**. Bar height represents proportion of trials in which animals responded to cue with more than one lick between trials 301 – 400 (60 s ITI) or 31 – 40 (600 s ITI). Error bars represent SEMs, and circles represent individual animals. ns: not significant, Welch’s t-test. **H**, The abruptness of change, a measure of how quickly an animal’s behavior changes at learning, determined by the steepness of the lick behavior cumsum curve (see *Methods*), is not different between groups. Bar height represents mean abruptness of change parameter for each group. Error bars represent SEMs, and circles represent individual animals. ns: not significant, Welch’s t-test.

**Fig. S3.**
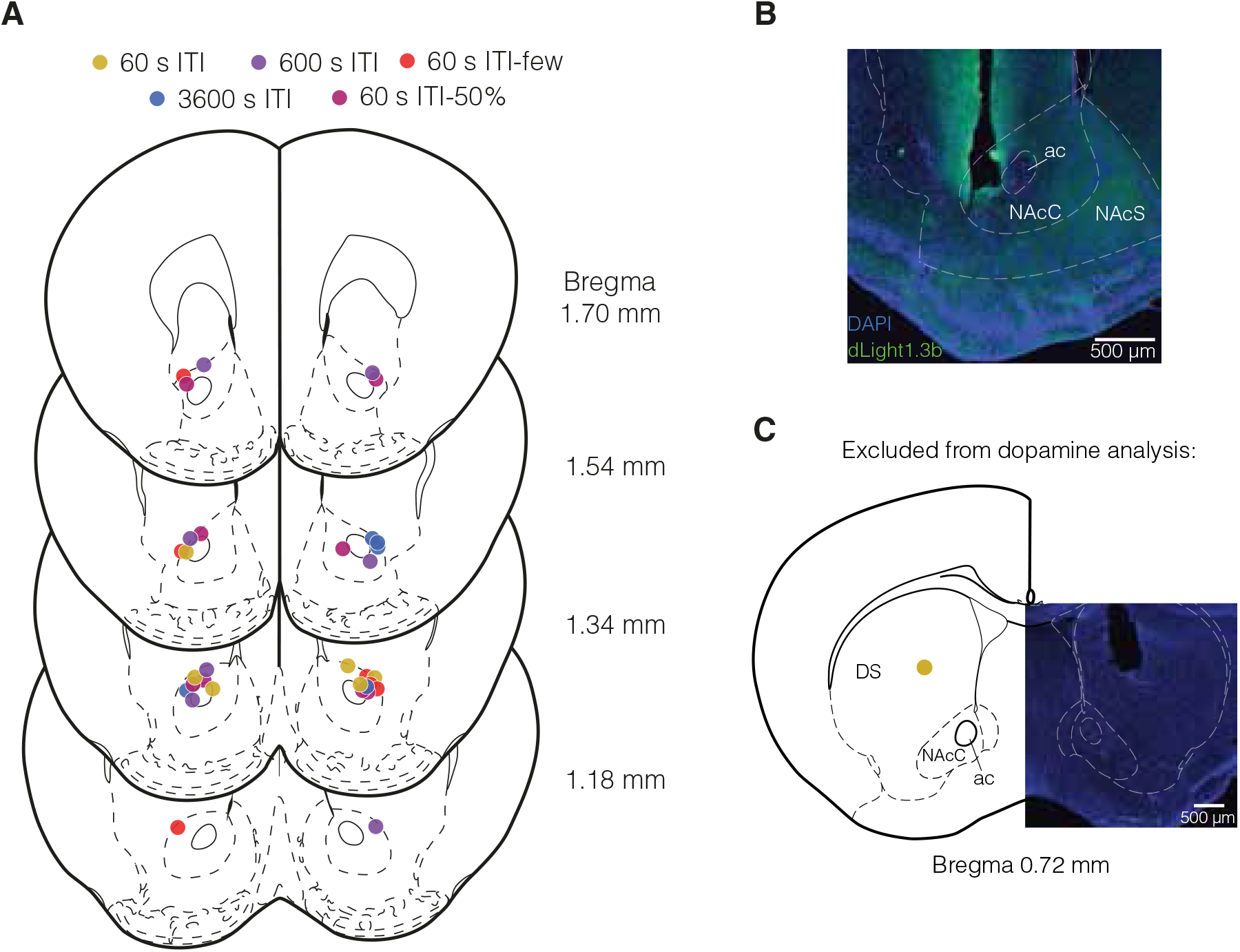
Fiber placements for dopamine measurement mice. **A**, Locations of center of optical fiber tip for subset of mice with fiber photometry dLight recordings from 60 s ITI (gold), 600 s ITI (purple), 3600 s ITI (blue), 60 s ITI-few (red), and 60 s ITI-50% (magenta) groups. **B**, Example histology from a single mouse. Blue is DAPI staining and green is dLight1.3b. NAcC, nucleus accumbens core; NAcS, nucleus accumbens shell; ac, anterior commissure. **C**, Fiber location from one 60 s ITI mouse excluded from dopamine analysis.

**Fig. S4.**
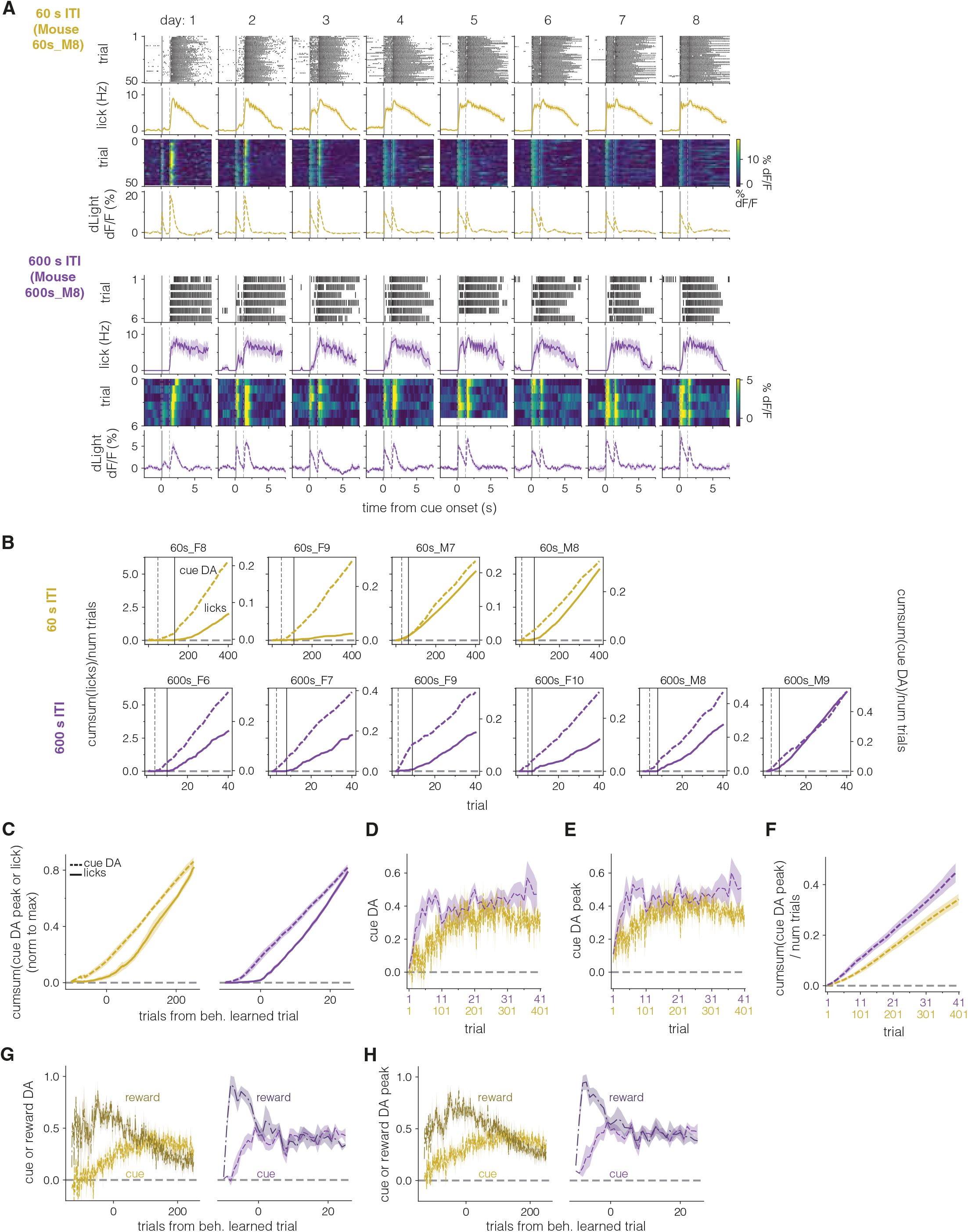
Taking the cumsum of cue-evoked dopamine and licking allows for determination of trials at which dopaminergic learning and behavioral learning occur in individual mice (A-B). Reward- and cue-evoked dopamine increase prior to behavioral learning (C-H). **A**, More lick raster, lick PSTH, dopamine response by trial, and average session dopamine PSTH plots from individual 60 s ITI and 600 s ITI example mice presented as in **Fig. 2C. B**, Cumsum of cue-evoked licks (solid, left axis) or of normalized cue-evoked dopamine (dashed, right axis) for all 60 s and 600 s ITI dopamine recording mice not shown in **Fig. 2D**. Both lick and cue-evoked dopamine values were divided by total trial number to display average responses across conditioning. Solid vertical lines represent learned behavior trial and dashed vertical lines represent dopamine learned trial. **C**, Average cumsum of cue-evoked licks (solid) and dopamine response (dashed) for 60 s ITI (*left*, gold) and 600 s ITI (*right*, purple) mice. Data were normalized by each animal’s final trial of conditioning and aligned to their learned trial before averaging, as in **Fig. 2F**, but using measurements of the peak of the dopamine response rather than the AUC. Lines represent means, and shading represents SEM. **D-E**, Average normalized AUC (**D**) or peak (**E**) cue-evoked dopamine across scaled trial units. Lines represent means, and shading represents SEM. **F**, Average cumsum of normalized cue-evoked dopamine responses in 60 s ITI (gold) and 600 s ITI (purple) mice as in **Fig. 2G** but using measurements of the peak of the dopamine response instead of the AUC. Cumsum curves divided by number of trials to account for different trial numbers between groups. Lines represent means, and shading represents SEM. **G-H** Average normalized AUC (**G**) or peak (**H**) cue (lighter, dashed) and reward (darker, dashdot) evoked dopamine responses aligned to each animal’s behavior learned trial. Note the increase in reward response prior to behavioral learning. Lines represent means, and shading represents SEM.

**Fig. S5.**
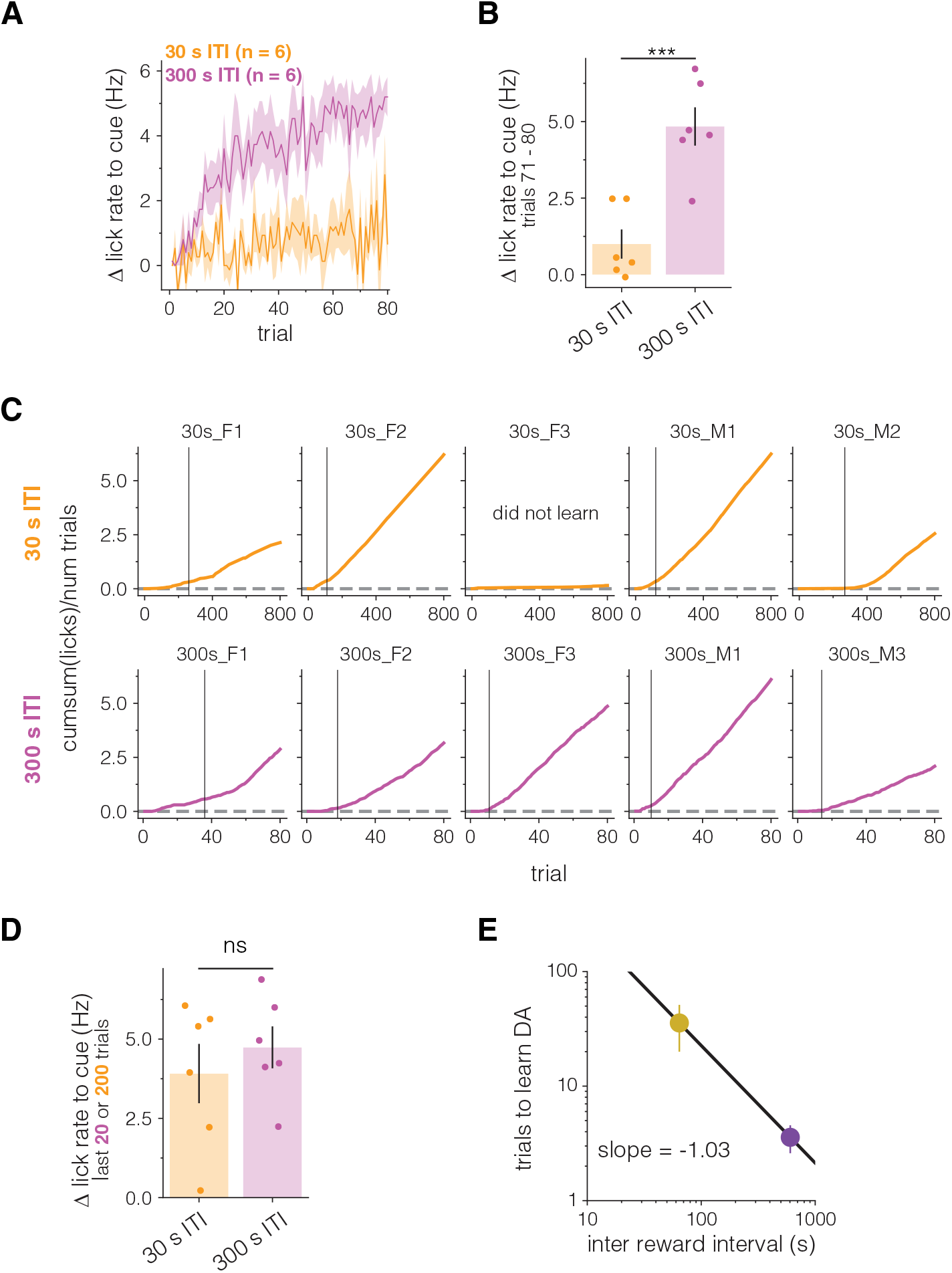
300 s ITI mice show evidence of learning in significantly fewer trials than 30 s ITI mice but do not differ in asymptotic behavior. A-B. 300 s ITI mice show significantly more cue-evoked licking than 30 s ITI mice over the first 80 trials of conditioning. **A**. Timecourse of average cue-evoked lick rate across 80 trials. **B** Mean change in cue-evoked lick rate for trials 71 – 80. ***p< 0.001, Welch’s t-test. **C**, Cumsum of cue-evoked licks for all 30 s and 300 s ITI mice not shown in **Fig. 3C**. Learned trial denoted by solid vertical line. **D**, Asymptotic cue-evoked lick rates are not significantly different between 30 s and 300 s ITI mice. Bars represent mean cue-evoked lick rates during trials 601–800 (30 s ITI) or trials 61-80 (300 s ITI). Error bars represent SEMs and circles represent individual mice. ns: not significant, Welch’s t-test. **E**, Mean trials to learn DA as a function of IRI for 60 s and 600 s ITI mice plotted on a log-log axis. Slope of line between points is -1.03, suggesting that dopaminergic learning scales proportionally with IRI, similar to behavioral learning. Error bars represent the standard deviation across mice.

**Fig. S6.**
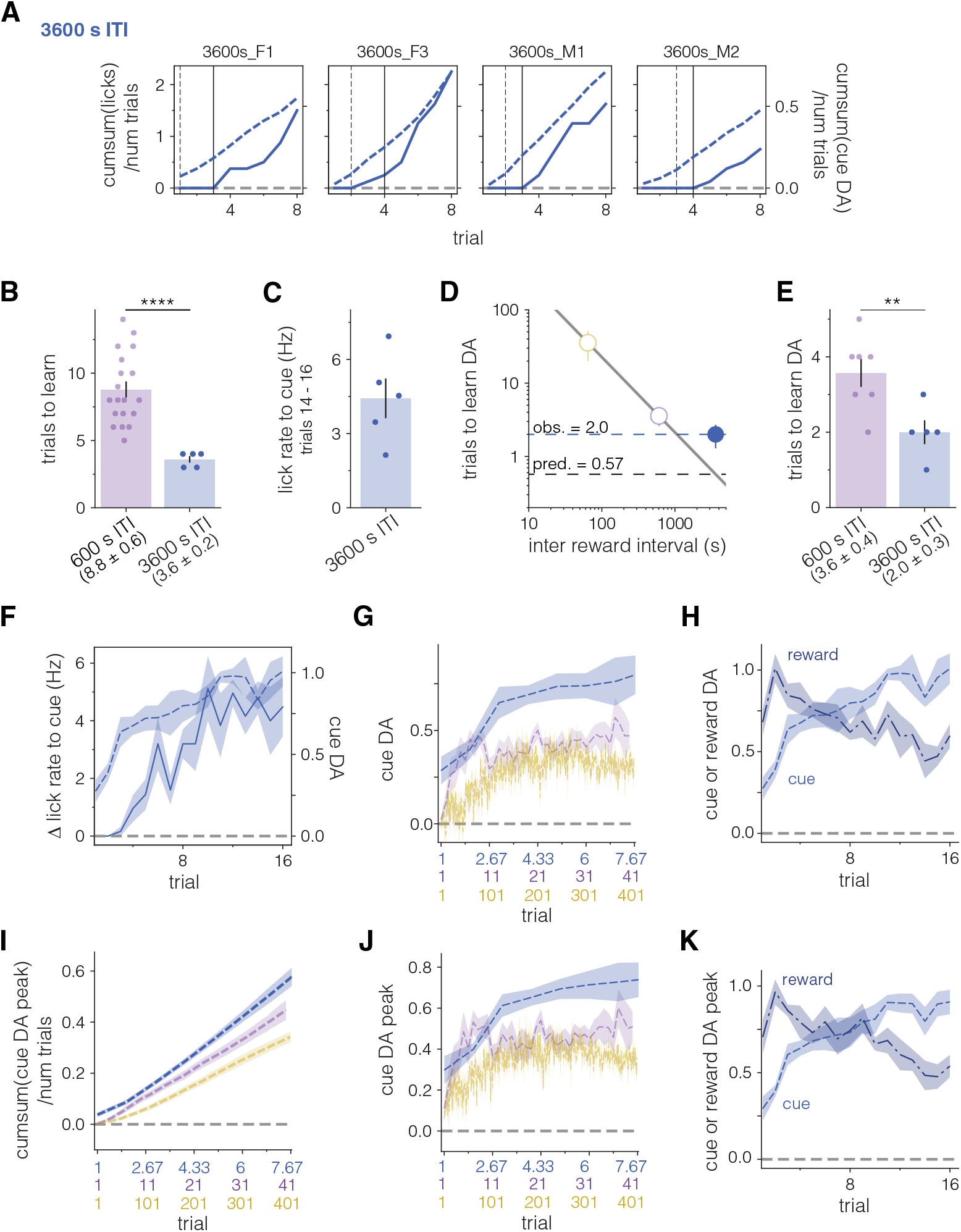
Cue-evoked dopamine in 3600 s ITI mice appears in fewer trials and reaches a higher asymptote than in 600 s ITI mice. **A**, Cumsum of cue-evoked licks (solid, left axis) or of normalized cue-evoked dopamine (dashed, right axis) for all 3600 s ITI mice not shown in **Fig. 3K**. Both lick and cue-evoked dopamine values were divided by total trial number to display average responses across conditioning. Solid vertical lines represent learned behavior trial and dashed vertical lines represent dopamine learned trial. **B**, Mean trials to learn for 3600 s and 600 s (same data as **Fig. 1F**) ITI mice. 3600 s ITI mice learn in significantly fewer trials. ****p<0.0001, Welch’s t-test. **C**, Asymptotic (trials 14-16) cue-evoked lick rate in 3600 s ITI mice. **D**, 3600 s ITI mice (blue circle, filled) take more trials to learn than predicted from the scaling relationship in **fig. S5E**. White filled circles and line are same data as **fig. S5E**. Dashed lines represent predicted and observed mean trials to learn for 3600 s ITI. **E**, Mean trials to learn cue-evoked dopamine for 3600 s and 600 s (same data as **Fig. 2E**) ITI mice. 3600 s ITI mice learn in significantly fewer trials. **p<0.01, Welch’s t-test. **F**, Average cue-evoked lick rate and normalized cue-evoked dopamine across all eight days of conditioning in 3600 s ITI mice (n = 5). **G**, Average normalized cue-evoked dopamine in 3600 s (blue, n = 5), 600 s (purple, same data as **fig. S4D**), and 60 s (gold, same data as **fig. S4D**) mice across scaled trials. **H**, Average normalized cue (dashed, lighter) or reward (dashdot, darker) evoked dopamine on each trial across all eight days of conditioning in 3600 s ITI mice (n = 5). **I**, Average cumsum of normalized cue-dopamine as in **Fig. 3Q**, analyzing peak of dopamine response rather than AUC. **J**, Average normalized cue-evoked dopamine as in **G**, analyzing peak of dopamine response rather than AUC. **K**, Average normalized cue- or reward-evoked dopamine as in **H**, analyzing peak of dopamine response rather than AUC.

**Fig. S7.**
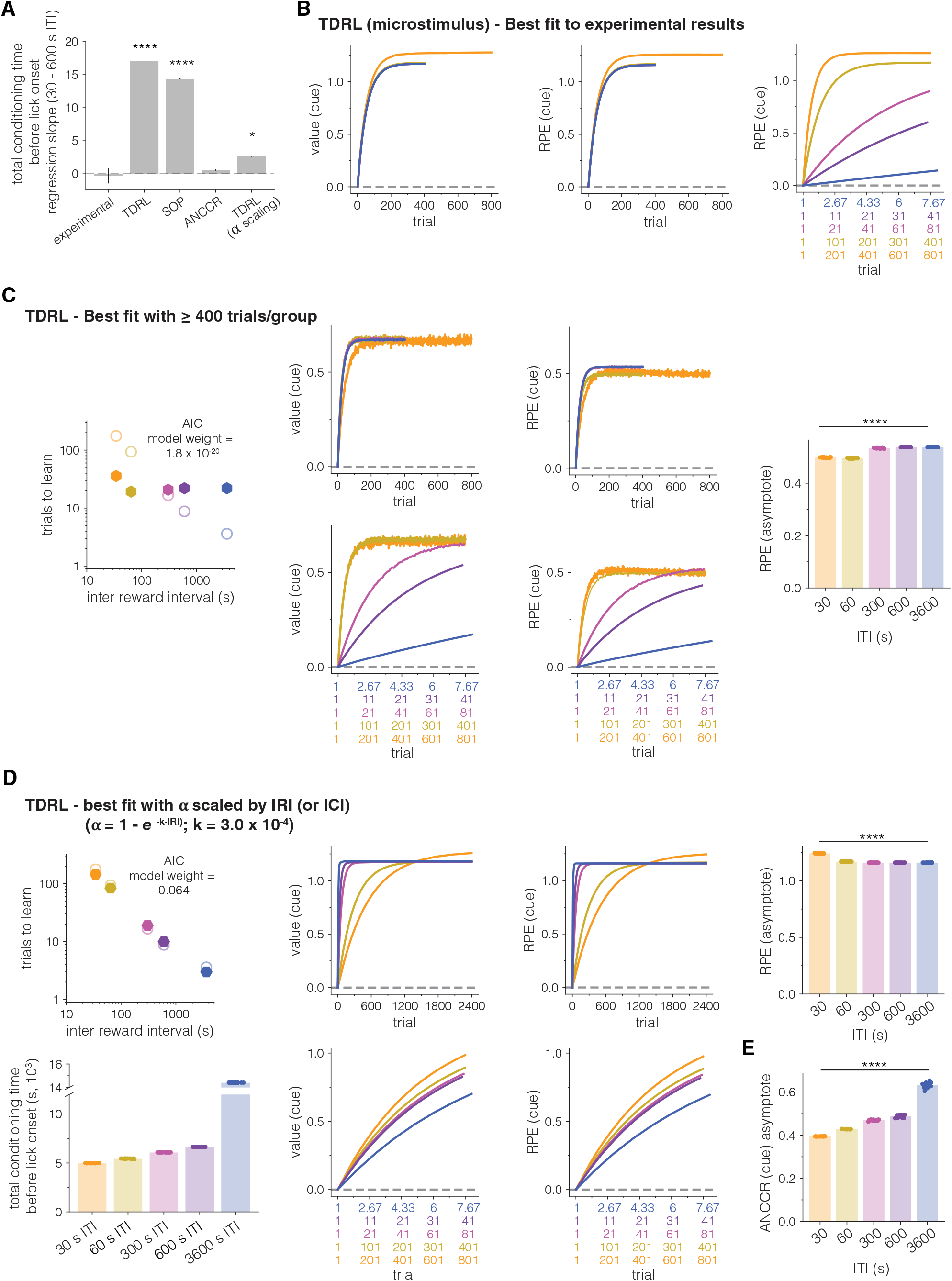
Total conditioning time analyses and additional TDRL simulation results. **A**, Slope of regressions for total time before lick onset as a function of ITI for 30 – 600 s ITI conditions from experimental results (**Fig. 4B**), TDRL (**Fig. 4F**), SOP (**Fig. 4J**), ANCCR (**Fig. 4N**), and TDRL with alpha scaling (**fig. S7D**). Error bars represent standard error. Asterisks represent significant difference from experimental results. **** p<0.0001, * p<0.05. t-test. **B**, Average value (maximum between cue and reward) (*left)* along with RPE at time of cue across unscaled (*middle)* and scaled (*right*) trials for best fit TDRL model (**Fig. 4, C to F)**. **C**, For comparisons between models (**Fig. 4**), simulations were run for the same number of trials as experimental groups. If each condition is instead run for at least 400 trials, qualitative scaling is observed between 30 and 60 s ITI conditions in the best fit model. This scaling, however, does not hold across all ITI groups. *Left*, Trials to learn (when value > threshold) as a function of IRI for the best-fit TDRL simulation with 400 trials for each ITI (see *Methods*). Hexagons represent mean trials to learn across iterations (n = 20 each). Open circles represent experimental data (same data as **Fig. 3, H and N**). Akaike information criterion (AIC) model weight comes from comparison with best fit SOP (**Fig. 4H**), and ANCCR (**Fig. 4L)** models. Please note that this AIC model does not penalize free parameters, i.e., is a conservative estimate biased against ANCCR (see *Methods* for rationale and **table S1** for AIC model weight when penalizing for parameters). *Middle*, Timecourse of value on each trial (maximum between cue and reward) or RPE at time of cue across iterations (n = 20 each) and plotted across scaled (*bottom*) or unscaled (*top*) trials. SEM occluded by thickness of mean lines. *Right*, Average RPE at time of cue during final conditioning trials. Bar height represents mean across iterations. Error bars represent SEMs, and circles represent individual iteration. **** p<0.0001, Kruskal-Wallis. Post-hoc Mann–Whitney U planned comparisons: 60 s vs. 600 s, p<0.0001; 60 s vs 3600 s, p<0.0001; 600 s vs. 3600 s, p = 0.38. **D**. In principle, a similar rule derived in ANCCR could be applied ad hoc to any model of associative learning. Here, we demonstrate that applying such a rule to TDRL improves fit to experimental results. Using the best fit model parameters determined during initial TDRL parameter sweep (**Fig. 4, C to F**), we replaced the learning rate, α, based on the equation α = 1 – *e* (−k· IRI) and performed another parameter sweep to determine the best fit *k*.*Left, top*, Trials to learn (when value > threshold) as a function of IRI for the best-fit TDRL simulation with scaled learning rate (see *Methods*). Hexagons represent mean trials to learn across iterations (n = 20 each). Open circles represent experimental data (same data as **Fig. 3, H and N**). Akaike information criterion (AIC) model weight comes from comparison with best fit SOP (**Fig. 4H**), and ANCCR (**Fig. 4L)** models. Please note that this AIC model does not penalize free parameters, i.e., is a conservative estimate biased against ANCCR (see *Methods* for rationale and **table S1** for AIC model weight when penalizing for parameters). *Left, bottom*, Total conditioning time prior to the emergence of behavior (threshold crossing). Symbols represent time for a single iteration, and bar height represents mean (n = 20 iterations per ITI condition). *Middle*, Timecourse of value on each trial (maximum between cue and reward) or RPE at time of cue for best fit *k* averaged across iterations (n = 20 each) and plotted across scaled (*bottom*) or unscaled (*top*) trials. SEM occluded by thickness of mean lines. *Right, top*, Mean RPE at time of cue during final conditioning trials. Bar height represents mean across iterations. Error bars represent SEMs, and circles represent individual iteration. **** p<0.0001, Kruskal-Wallis. Post-hoc Mann–Whitney U planned comparisons: 60 s vs. 600 s, 60 s vs 3600 s, 600 s vs. 3600 s, all p<0.0001. **E**. Mean ANCCR (the variable postulated to be encoded by dopamine in the ANCCR framework) during final conditioning trials from simulations in **Fig. 4, K to N**. Bar height represents mean across iterations. Error bars represent SEMs, and circles represent individual iteration. **** p<0.0001, Kruskal-Wallis. Post-hoc Mann–Whitney U planned comparisons: 60 s vs. 600 s, 60 s vs 3600 s, 600 s vs. 3600 s, all p<0.0001.

**Fig. S8.**
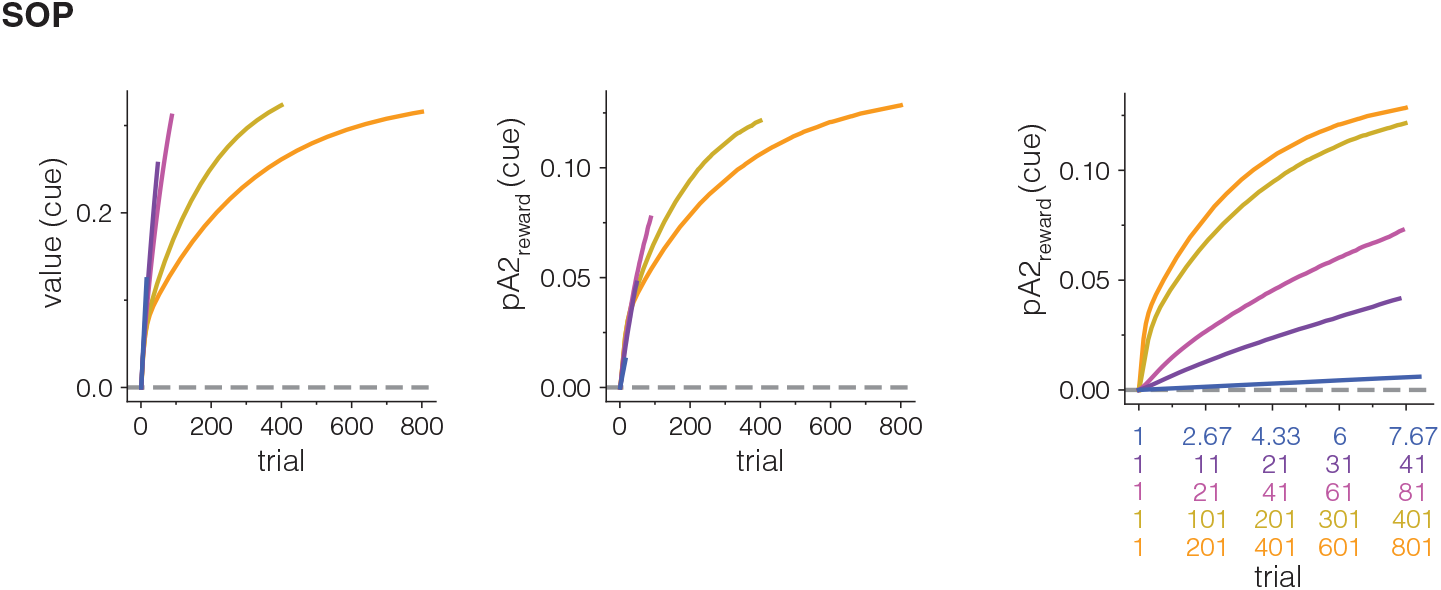
Additional SOP simulation results. Average value (maximum between cue and reward) (*left)* and *pA*2_*reward*_ (*middle)* across unscaled trials and *pA*2_*reward*_ across scaled trials (*right*) for best fit SOP model (**Fig. 4, G to J**). In SOP, when a cue-reward association is learned, the cue attains the ability to activate the secondary, decayed representation of the reward (A2reward), which is thought to underlie conditioned responding to the cue after learning. *pA*2_*reward*_ at time of cue refers to the proportion of reward representation elements driven into the secondary activation state (A2) by cue presentation.

**Fig. S9.**
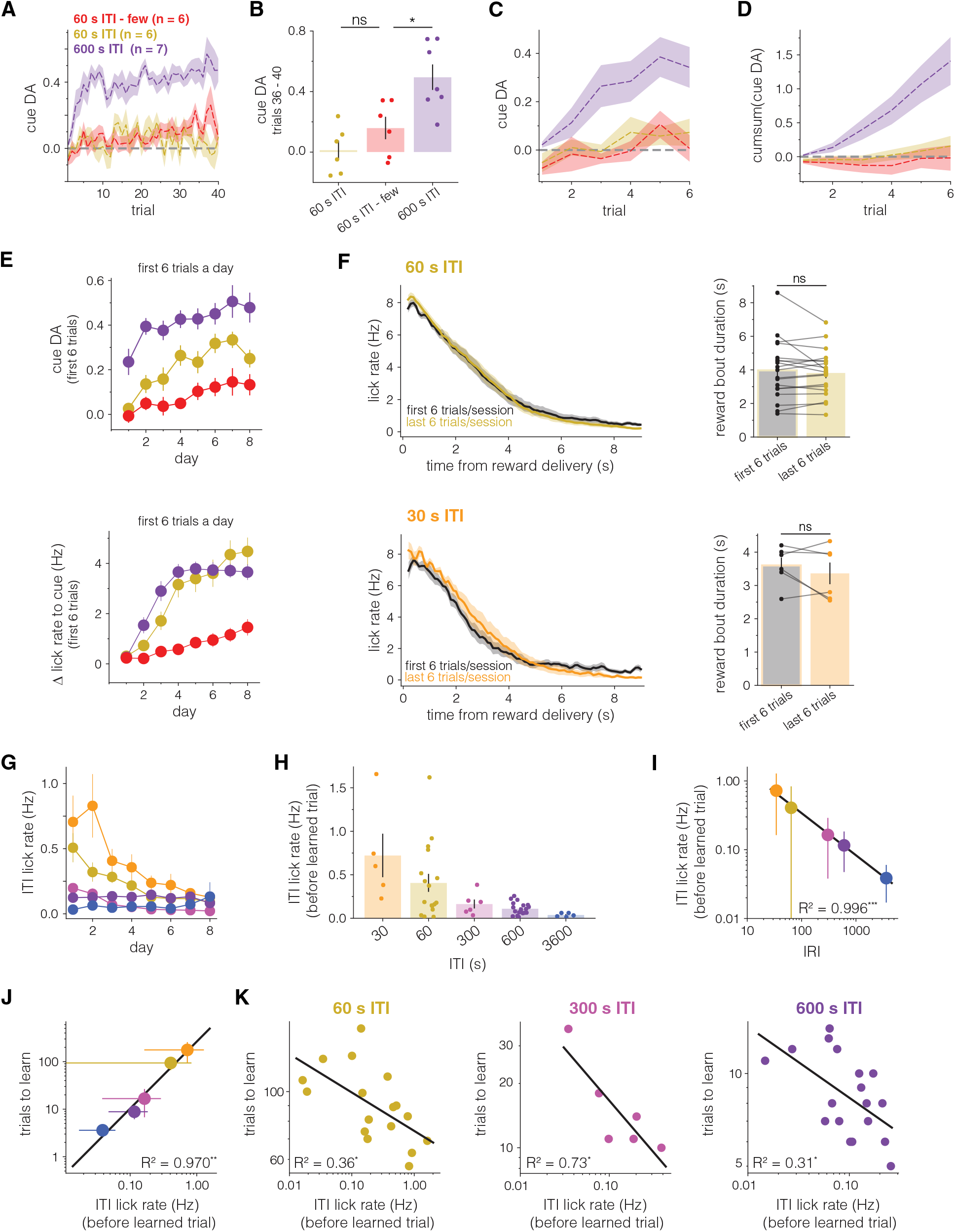
Satiety does not account for differences in learning between groups (A-F). Lick rate during ITI scales with reward rate, and therefore, trials to learn (G-K) A-B, Cue-evoked dopamine in 60 s ITI-few mice (red, n = 6) evolves similarly to that of 60 s ITI mice and is significantly lower than that of 600 s ITI dopamine at the end of conditioning. **A**. Timecourse of normalized cue-evoked dopamine across first 40 trials of conditioning. Lines represent animal means and shaded region represents SEM. **B**. Mean cue-evoked dopamine from trials 36-40. *p<0.05, ns: not significant. **C-D**, Average (**C)** or cumsum (**D)** of normalized cue-evoked dopamine responses in 60 s ITI-few (red, n = 6), 60 s ITI (gold, n = 5), and 600 s ITI (purple, n = 7) over the first 6 trials on the first day of conditioning, which are the first exposures to both cues and rewards and during which satiety is not expected to differ between groups. Lines represent means across animals and shaded region represents SEM. Note the similarities between 60 s ITI-few and 60 s ITI groups, and the greater dopamine responses in 600 s ITI despite equivalent cumulative number of rewards experienced between groups. **E**, Average cue-evoked dopamine (*top*) and cue-evoked lick rates (*bottom*) for the first six trials on each day in 60 s ITI-few (lick: n = 18, cue DA: n = 6), 60 s ITI (lick: n = 19, cue DA: n = 6), and 600 s ITI (lick: n = 19, cue DA: n = 7) mice. Dopamine responses between 60 s ITI and 60 s ITI-few mice diverge across conditioning, suggesting that satiety following the first 6 trials throughout a session does not drive the differences in dopamine signaling or lick behavior seen between groups. **F**, Average reward bout duration for first 6 trials and last 6 trials a session is not different in 60 s ITI (n = 19, *top*) or 30 s ITI (n = 6, *bottom*) mice, suggesting a lack of within session satiety and decreased motivation. *Left*, Lick PSTHs aligned to reward delivery. *Right*, Average duration of reward bout for first 6 trials and last 6 trials per session averaged across all conditioning days. Error bar represents SEM. Circles represent individual mice, and lines connect data from a single mouse. ns: not significant, paired t-test. **G**, Average lick rate during the ITI (defined as licks following cessation of consummatory bout until next cue onset) over days. Note the clear separation between groups in early days that greatly reduces over conditioning, consistent with greater context-reward associations in shorter ITI groups that extinguish over time. **H**, Average ITI lick rates preceding each individual animal’s learned trial. **I**, Average ITI lick rates preceding learning, as a function of inter-reward interval (IRI) plotted on log-log axis. Circles represent mean trials to learn per group and error bars represent standard deviation. Solid black line is best fit regression line (R^2^ = 0.996, *** p<0.001). This suggests a strong positive relationship between reward rate and ITI licking, consistent with differential context-reward associations between groups due to an effect of context extinction (i.e. lower rate of reward in the context during long ITI). **J**, Average learned trial as a function of ITI lick rates preceding learning plotted on log-log axis. Circles represent mean trials to learn per group and error bars represent standard deviation. Solid black line is best fit regression line (R^2^ = 0.970, ** p<0.001), suggesting an inverse relationship between ITI lick rate and learning rate. **K**, Contrary to what may be predicted from the between-group relationship, within each ITI group there is a positive relationship between ITI lick rate and learning rate, suggesting reduced context-reward associations are not the primary driver of cue-reward learning. *Left*, 60 s ITI (R^2^ = 0.36, ** p<0.05), *middle*, 300 s ITI (R^2^ = 0.73, ** p<0.05), *right*, 600 s ITI (R^2^ = 0.31, ** p<0.05).

**Fig. S10.**
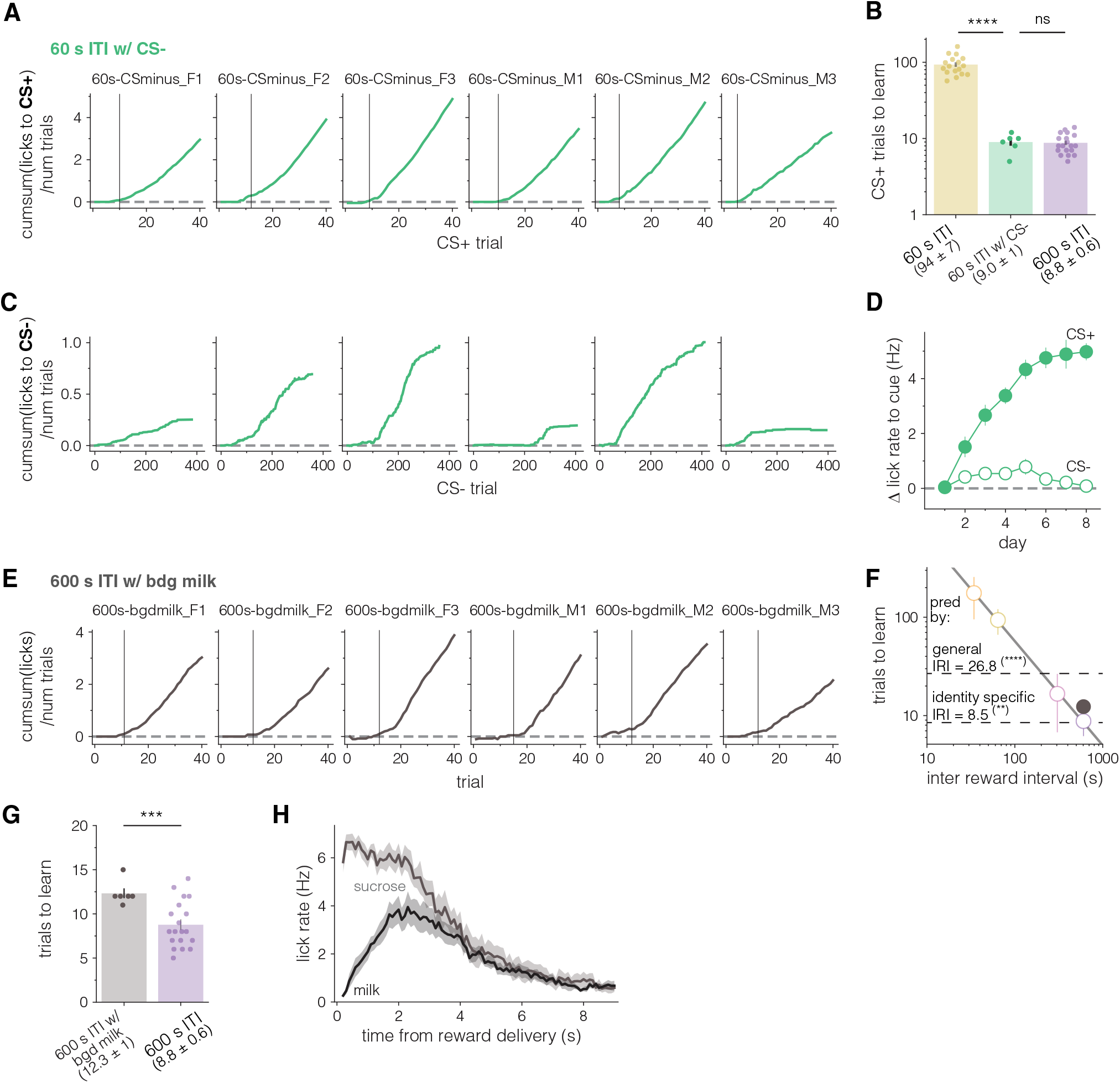
Trials to learn in “60 s ITI with CS-” and “600 s ITI with background milk” mice. **A**, Cumsum of CS+ evoked licks for all “60 s ITI with CS-mice” across 40 trials. **B**, 60 s ITI with CS-mice learn in significantly fewer CS+ trials than 60 s ITI, similar to 600 s ITI mice (60 s and 600 s ITI: same data as **Fig. 1F)**. **** p < 0.0001 not significant, Welch’s t-test. **C**, Cumsum of CS-evoked licks for all 60 s ITI with CS-mice across all CS-presentations before the 40th CS+ trial. Note that despite the lower average licks to CS-compared to CS+ licking (**A**), every mouse showed at least some CS-evoked licking, evidence that all mice could hear and respond to the tone. **D**, Timecourse of average CS+ (filled) or CS-(open) evoked lick rate as a function of days of conditioning. Circles represent mean change in lick rate per day, and error bars represent SEMs. **E**, Cumsum of cue-evoked licks for all 600 s ITI with background milk mice across 40 trials. **F**, Predicted and actual trials to learn from experiment shown in **Fig. 5J**. Dashed lines represent predicted trials to learn based the relationship determined in **Fig. 3H** using either the general IRI including both chocolate milk and sucrose (upper line) or using the interval between sucrose deliveries alone (lower line). The actual observation (brown filled circle) is much closer to a scaling by an identity-specific IRI of the sucrose than a general IRI of either sucrose or chocolate milk. The fact that the actual trials to learn do not exactly match the identity-specific IRI suggests slight generalization between sucrose and chocolate milk rewards due to shared sensory features (e.g., both are sweet, liquid rewards). Open circles and gray line same data as **Fig. 3H**. ****p<0.0001, **p<0.01. One-sample t-test. **G**, Mean trials to learn in 600 s ITI with background milk vs. 600 s ITI (same data as **Fig. 1F)** mice. **H**, Average lick PSTH aligned to reward delivery for sucrose (gray) or chocolate milk (black) for all 600 s ITI with background milk over eight days of conditioning. Shaded error represents SEM across mice. *** p < 0.001, Welch’s t-test.

**Fig. S11.**
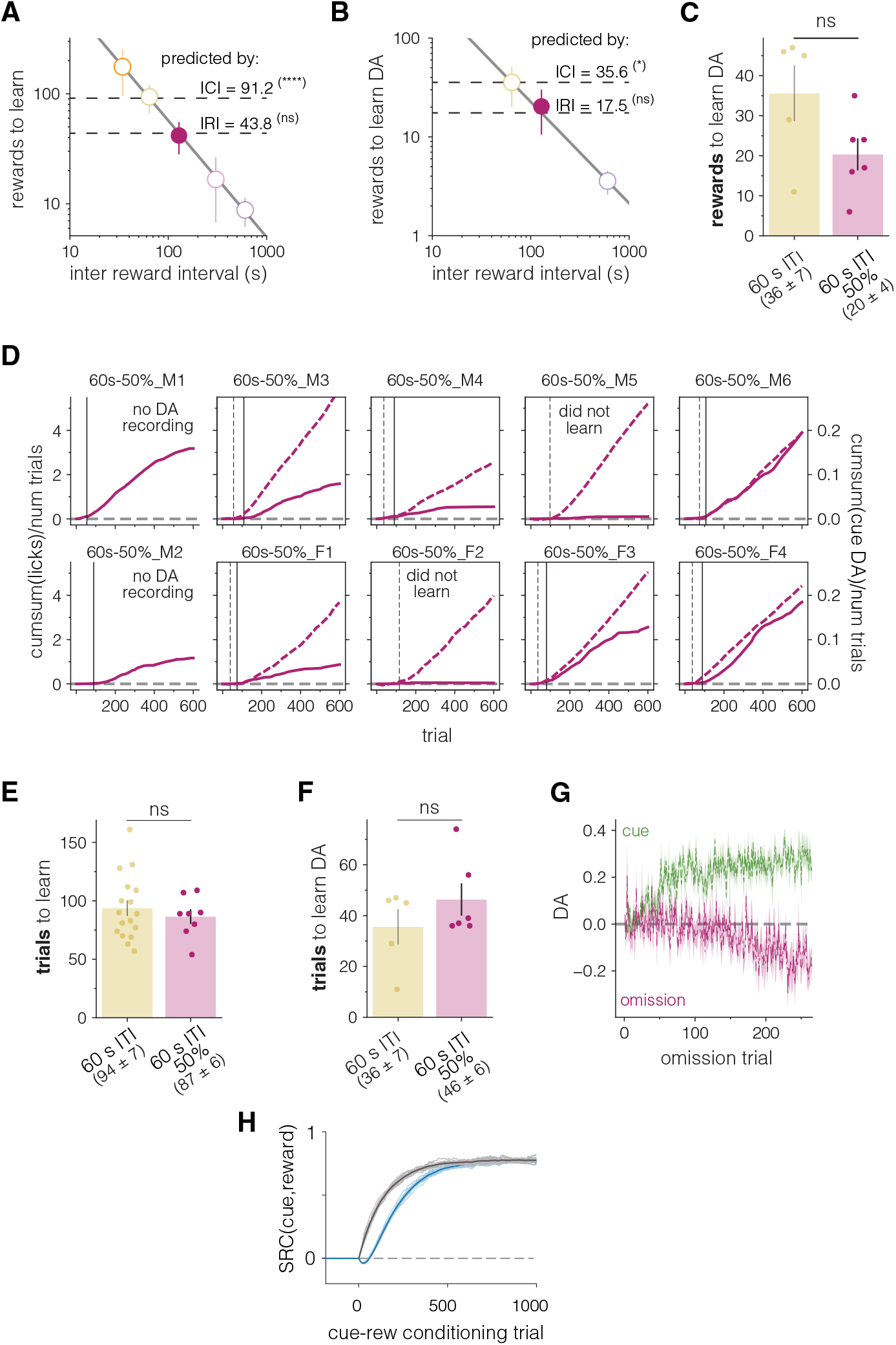
60 s ITI-50% mice learn in the same number of trials as 60 s ITI mice despite receiving half the number of rewards. **A**, Predicted and actual rewards to learn in 60 s ITI-50% mice. The previous experiments did not disambiguate the inter-cue interval (ICI) from the IRI because they are equivalent when rewards follow cues with 100% probability. If the learning rate is scaled by the CS+ ICI rather than the IRI, 60 s ITI-50% mice would be expected to learn after 91.2 rewards based on the relationship determined in **Fig. 3H** (upper dashed line), similar to 60 s ITI mice, which have an identical inter-cue interval. However, if IRI is the driver of learning rate scaling 60 s ITI-50% would be expected to learn in 43.8 rewards (lower dashed line). The observed number of rewards to learn (magenta filed circle), 42, is consistent with learning rate scaling by IRI. Open circles and gray line same data as **Fig. 3H**. ****p<0.0001, ns: not significant. One-sample t-test. **B**, Predicted and actual rewards until the emergence of cue-evoked dopamine in 60 s ITI-50% mice. As in **A**, ICI scaling of learning rate predicts dopaminergic learning in 35.6 rewards, while IRI scaling of learning rate predicts 17.5 rewards. The observed (magenta filed circle) dopaminergic learning in 20 rewards is consistent with learning rate scaling by IRI. Open circles and gray line are the same data as **fig. S5E**. *p<0.05, ns: not significant. One-sample t-test **C**, Number of rewards after which cue-evoked dopamine emerges in 60 s ITI-50% and 60 s ITI (same data as **Fig. 2E**) mice. Mice that did not show evidence of learning or did not have dopamine recordings (**Fig. 6B**) were excluded from analysis. ns: not significant. Welch’s t-test. **D**, Cumsum of cue-evoked licks (solid, left axis) or of normalized cue-evoked dopamine (dashed, right axis) as a function of all trials for all 60 s ITI-50% mice. Both lick and cue-evoked dopamine values were divided by total trial number to display average responses across conditioning. Solid vertical lines represent learned behavior trial and dashed vertical lines represent dopamine (cue) learned trial. Dopamine was not recorded from two initial mice and two other mice did not show evidence of behavioral learning (see *Methods*). *(Fig. S11. Continued)* **E**, 60 s ITI-50% mice learn the cue-reward association in about the same number of cue presentations/trials as 60 s ITI mice (same data as **Fig. 1F**). Mice that did not show evidence of learning (**D**) were excluded from analysis. Note that for 60 s ITI mice, cues/trials to learn and rewards to learn represent the same values (**Fig. 6C**). Ns: not significant. Welch’s t-test. **F**, Number of trials after which cue-evoked dopamine emerges in 60 s ITI-50% and 60 s ITI (same data as **Fig. 2E)** mice. Note that for 60 s ITI mice, cues/trials to learn and rewards to learn dopamine represent the same values (**fig. S11C**). ns: not significant. Welch’s t-test. **G**, Average normalized dopamine responses to cue presentation (green, dashed) or reward omission (magenta, dashdot) across reward omission trials in 60 s ITI-50% mice (n = 6). **H**, Growth curves for the prospective association in simulations of the best-fit ANCCR model from **Fig. 4** as a function of trials with (blue line) and without (grey line) prior exposure to cues, indicating the presence of latent inhibition when “omission” trials are presented together prior to conditioning. For the latent inhibition condition, 200 cue-only presentations were delivered prior to cue-reward conditioning. ITI in all cases was set to 30 s.

## Supplementary Note 1

### Formal Derivation of Proportional Scaling of Retrospective Learning Rate by Inter-Reward Interval

In the intuitive derivation above, we made the critical assumption that the time constant of decay of the eligibility trace *T* is very short. Here we will derive the expression for the timescale of history for *M*_←*cr*_ and *M*_←*r*−_ or *M*_←*c*−_ for a general *T* and show that they are 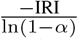 and 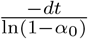 respectively provided 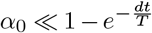 (the formal condition of “very short” T, which as it turns out, need not be very short if *α*_0_ is low).

We will first calculate the timescale of history for *M*_←*r*−_ or *M*_←*c*−_, both of which are updated using successive delta-rule like updates every *dt* with learning rate *α*_0_. To this end, we will consider an event train of event type *x* (either cue or reward) and denote the estimate of *M* ←_*x*−_ at the time of the *n*th timepoint/sample (i.e., at time *ndt*) as *M* (*n*) and the corresponding instantaneous eligibility trace as *E*(*n*). The delta-rule for update of *M* (*n*) can then be written as:

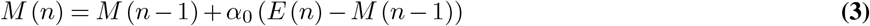

This update occurs every *dt*. Simplifying, we get

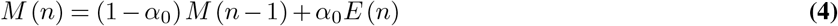

Similarly, expanding *M* (*n* − 1) in terms of *M* (*n* − 2) we get

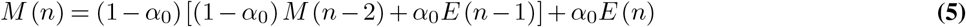

Expanding this recursive expression till *M* (0) i.e., the initialized value of *M*_←*x*−_ prior to the first sample/timepoint, we get

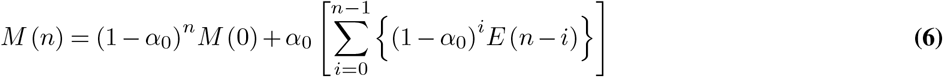

We will assume that *M* (0) = 0 and replace *i* with *n* − *i* to count from the first sample to get

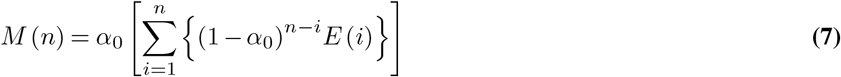

To calculate the timescale of history of *M*_←*x*−_, we now just need to substitute an expression of *E*(*i*) for a given event train of event type *x* and then identify the exponential time constant of growth.

We will next show that the eligibility trace for an event train occurring at a constant rate with period *t*_*x*_ rides on a trendline that is a saturating exponential growth curve with a time constant equal to the eligibility trace decay time constant *T*. In other words, the growth curve of the eligibility trace is 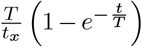.

To this end, we will assume a periodic event train for *x* from here on for analytical simplicity. Despite this simplistic assumption, our simulations show that the key results from this derivation hold for more realistic event trains.

Assume that the period of this periodic event train is *t*_*x*_ and that just before the *j*th occurrence of event *x* the eligibility trace is *E*_*j*_. If so, by definition, the eligibility trace immediately after the occurrence of the *<u>j</u>*<u>t</u>h occurrence is *E*_*j*+1_. Therefore, the mean eligibility trace in the time between the *j*th and (*j* + 1)th occurrences, denoted as 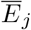 is by definition:

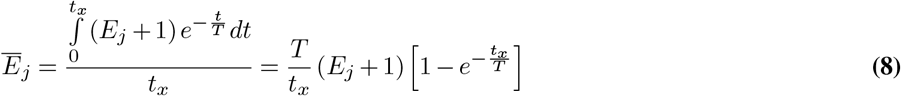

For a periodic event train, *E*_*j*_ can be calculated as:

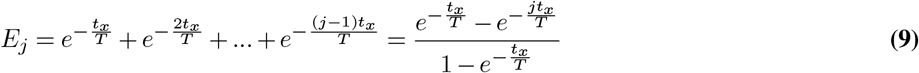

Substituting this into the previous equation, we see that

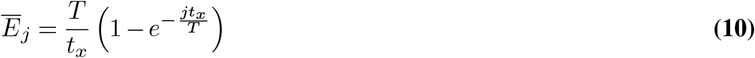

This means that the average trendline of the eligibility trace grows as a saturating exponential curve with time constant *T* and asymptote *T/t*_*x*_. An interpolated continuous time estimate of the above trendline is simply 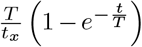. The exact eligibility trace curve will see-saw around this trendline as it grows by 1 on every occurrence of the event and then decreases exponentially until the next occurrence. However, since our goal here is to just find the overall timescale of growth for the mean eligibility trace *M*_←*x*−_, it is sufficient to substitute the above interpolated trendline for the eligibility trace into Equation (7).

Thus, at the *i*th sample in Equation (7), *E*(*i*) can be written as 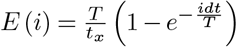. Please note that the *i* here refers to the *i*th sample and not the complex square root of -1. Therefore, *M* (*n*) from from Equation (7) can be rewritten as:

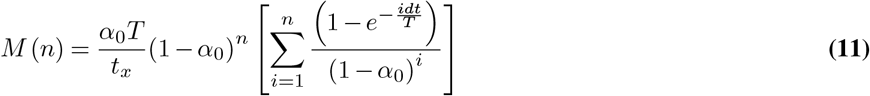

It can be shown that this simplifies to:

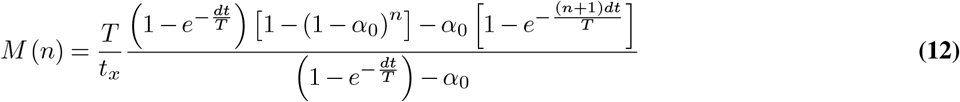

This equation shows the growth curve for the average estimate of the trendline of the eligibility trace of event *x*. It can be verified that when *α*_0_ is set to 1 (i.e., there is no averaging), the above curve reduces to the trendline of the eligibility trace that we assumed earlier i.e., 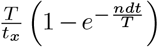. When 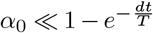 (i.e., eligibility trace decays quickly and there is slow averaging of eligibility trace), the first term in the numerator and denominator dominate and therefore *M*_←*x*−_ grows as a saturating exponential curve of the form [1− (1− *α*_0_)^*n*^]. Equating [1− (1− *α*_0_)^*n*^] with a saturating exponential growth curve of the form 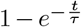, we can see that the time constant equals 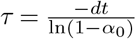. Notice that this is also the same time constant that measures the exponentially decaying influence with which an event that occurred in the past influences the current estimate of *M*_←*x*−_. This is thus the timescale of history for *M*_←*x*−_.

We can now similarly estimate the time constant of growth for the predecessor representation *M*_←*xy*_ (typically *x* is a cue and *y* is a reward). Intuitively, one can infer this from the above time constant for *M*_←*x*−_ of 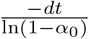 by noting that the numerator measures the time interval between successive delta-rule updates and the alpha in the denominator measures the corresponding learning rate for the delta-rule. By similarity, the interval between successive delta-rule updates for *M*_←*xy*_ is the inter-*y*-interval (denoted IYI) and the corresponding learning rate is *α*. Therefore, we can intuitively infer that the corresponding time constant should be 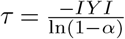. To see this more formally, let us assume a periodic event train for *y* with *x*’s occurring just prior to each occurrence of *y*. In this case, if one sets the continual sampling to occur with *dt* = IYI, the baseline updates for *x* will be identical in timing to the predecessor representation updates calculated when *y* occurs. The only difference will be that the delta-rule updates for predecessor representation will have a learning rate *α* instead of *α*_0_. Thus the same results as in Equation (12) will hold for the predecessor representation but with *dt* set to IYI and *α*_0_ set to *α*. This means that the corresponding time constant of growth for the predecessor representation is 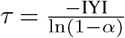.

Collectively, we can now see that to obtain the same time constant of growth for the baseline and the predecessor representation one needs to satisfy the following relationship:

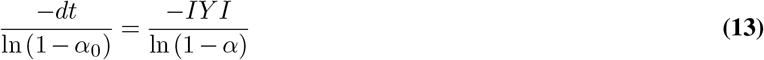

This implies that *α* should be set to

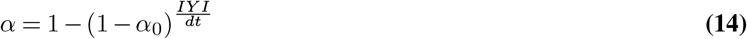

For calculating the predecessor representation between a cue and reward, the retrospective updates occur at reward and hence the corresponding learning rate should be:

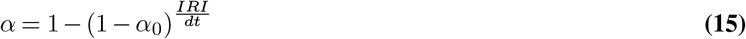

Notice that the IRI here refers to the inter-reward interval of the specific type of reward and is thus reward identity-specific, a prediction that is tested in **Fig 5, J to L**.

### Setting of Eligibility Trace Time Constant

It is intuitively clear that the eligibility trace time constant *T* needs to be set to match the timescales operating in the environment. This is because if the eligibility trace decays too quickly, there will be no memory of past events, and if it decays too slowly, it will take a long time to correctly learn event rates in the environment.

Further, as we showed in the previous section, the asymptotic value of *M*_←*x*−_ for an event train at a constant rate *λ*_*x*_ with average period *t*_*x*_ is *T/t*_*x*_ = *Tλ*_*x*_. This means that the neural representation of *M*_←*x*−_ will need to be very high if *T* is very high and very low if *T* is very low. Since every known neural encoding scheme is non-linear at its limits with a floor and ceiling effect (e.g., firing rates can’t be below zero or be infinitely high), the limited neural resource in the linear regime should be used appropriately for efficient coding. A linear regime of operation for *M*_←*x*−_ is especially important in ANCCR since the estimation of the successor representation by Bayes’ rule depends on the ratio of *M*_←*x*−_ for different event types. Such a ratio will be highly biased if the neural representation of *M*_←*x*−_ is in its non-linear range. Assuming without loss of generality that the optimal value of *M*_←*x*−_ is *M*_opt_ for efficient linear coding, we can define a simple optimality criterion for the eligibility trace time constant *T*. Specifically, we postulate that the net sum of squared deviations of *M*_←*x*−_ from *M*_opt_ for all event types should be minimized at the optimal *T*. The net sum of squared deviations denoted by *SS* can be written as:

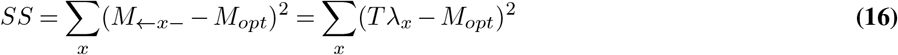

Where the second equality assumes asymptotic values of *M* _←*x*−_. The minimum of *SS* with respect to *T* will occur when 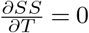. It is easy to show that this means that the optimal *T* is:

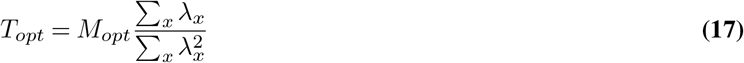

For typical cue-reward experiments with each cue predicting reward at 100% probability,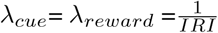. Substituting into the above equation, we get:

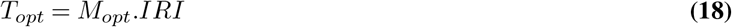

Thus, in typical experiments with 100% reward probability, the eligibility trace time constant should be proportional to the IRI (or equivalently the inter-cue interval). In the simulations of ANCCR in **Fig. 4**, we used Equations (15) and (18) to set the learning rate and eligibility trace time constant respectively. These relationships explain why the learning rate scaling for a one-hour IRI deviates from perfect linearity in **Fig. 4**. This is because the optimal *T* for one-hour IRI equaled 1800 s (since best-fit *M*_opt_ was 0.5). At this value, the best-fit *α*_0_ of 4 × 10^−5^ was not much smaller than 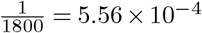 and thus the overall learning of the baseline *M*_←*c*−_ and the predecessor representation *M*_←*cr*_ was slower than the perfectly linear rate for the lower IRIs (see Equation (12)).

Both Equations (15) and (17) require the real-time estimation of inter-event intervals which should be learned separately by an animal. Since the goal of the current study was not to understand how the real-time learning of inter-event intervals is achieved by an animal, we simply assumed the correct rates of the events in the simulations of ANCCR. Additional theoretical and experimental work will be needed to better understand how these quantities are set in real-time.

**Supplementary Table S1.**
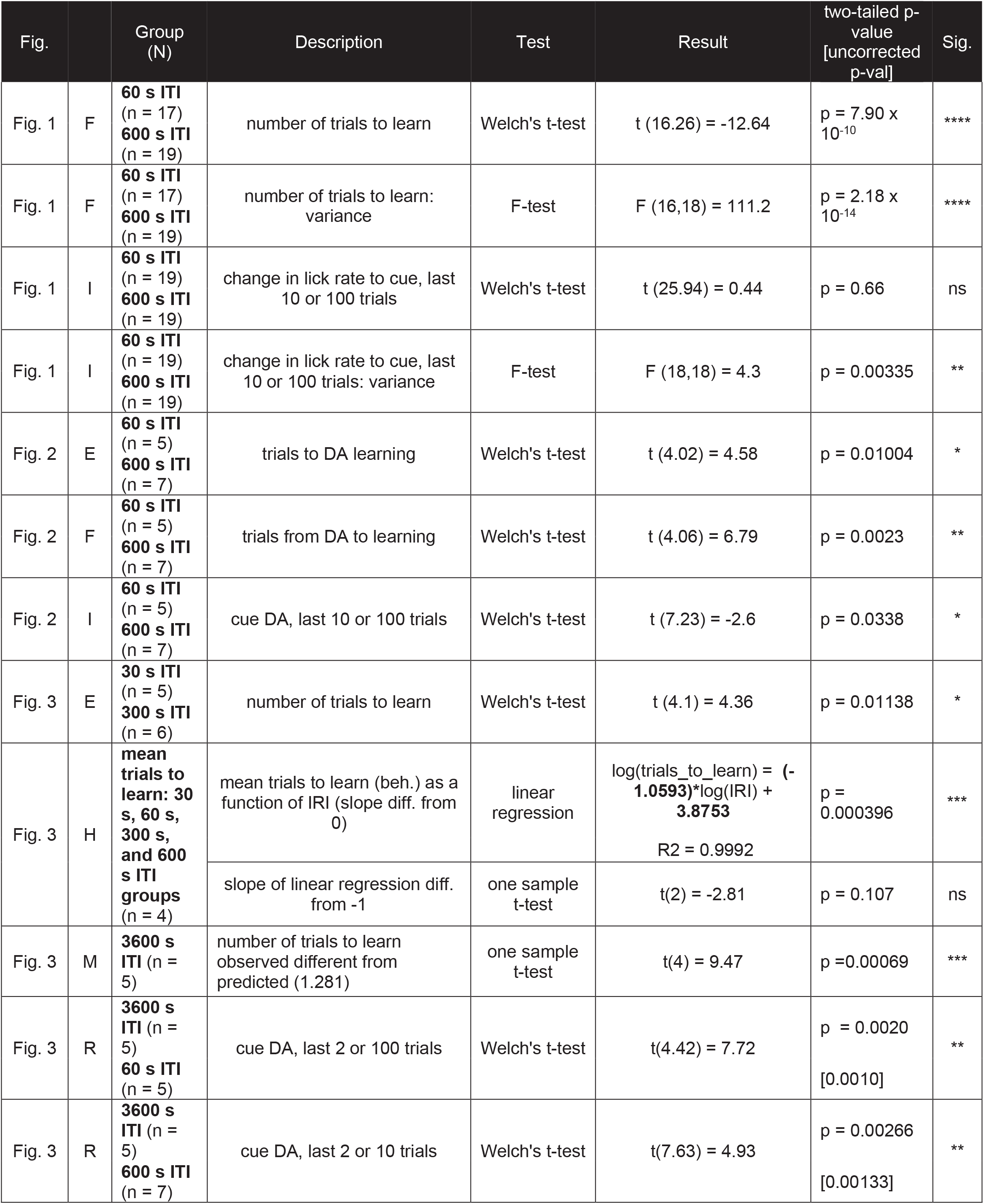
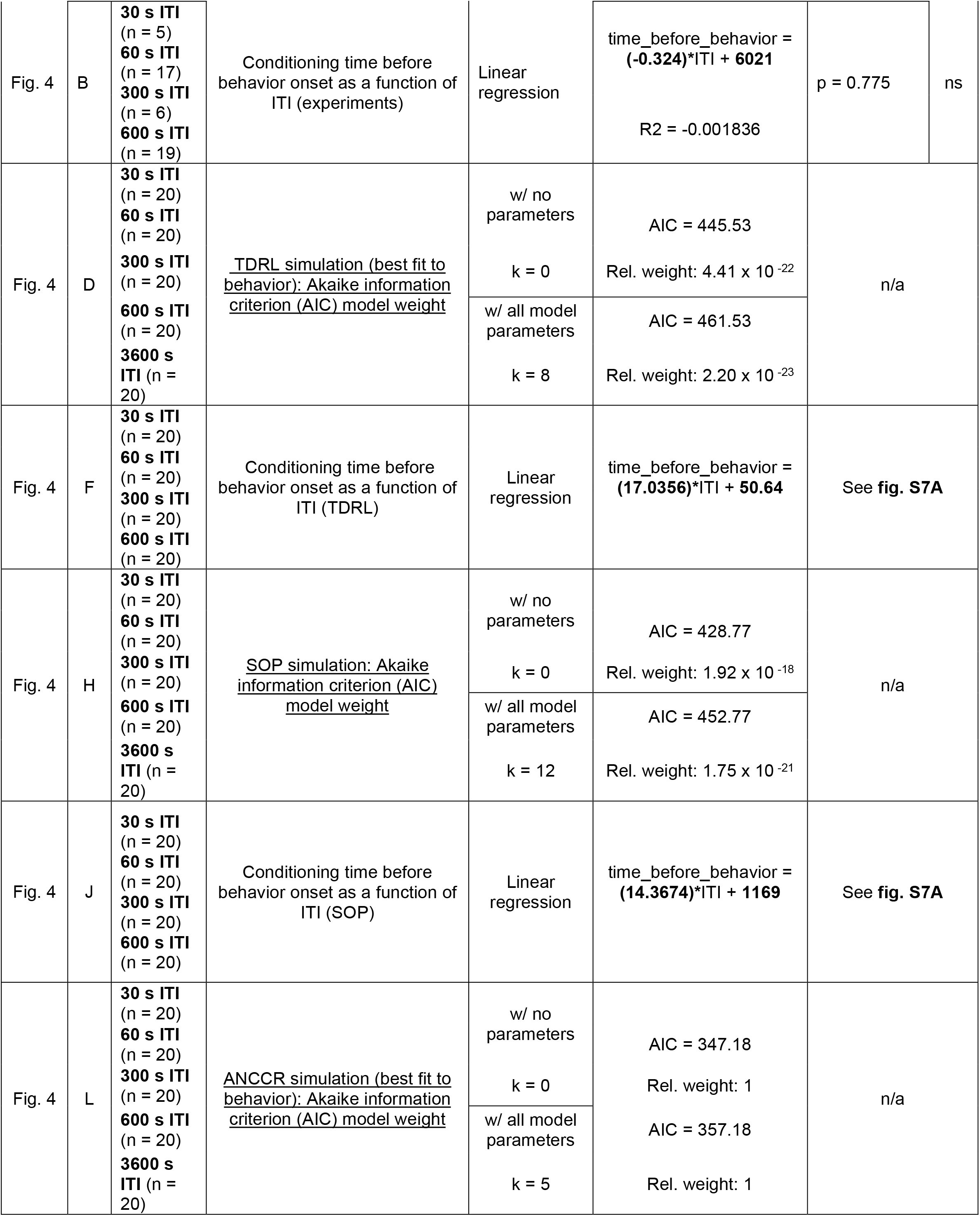
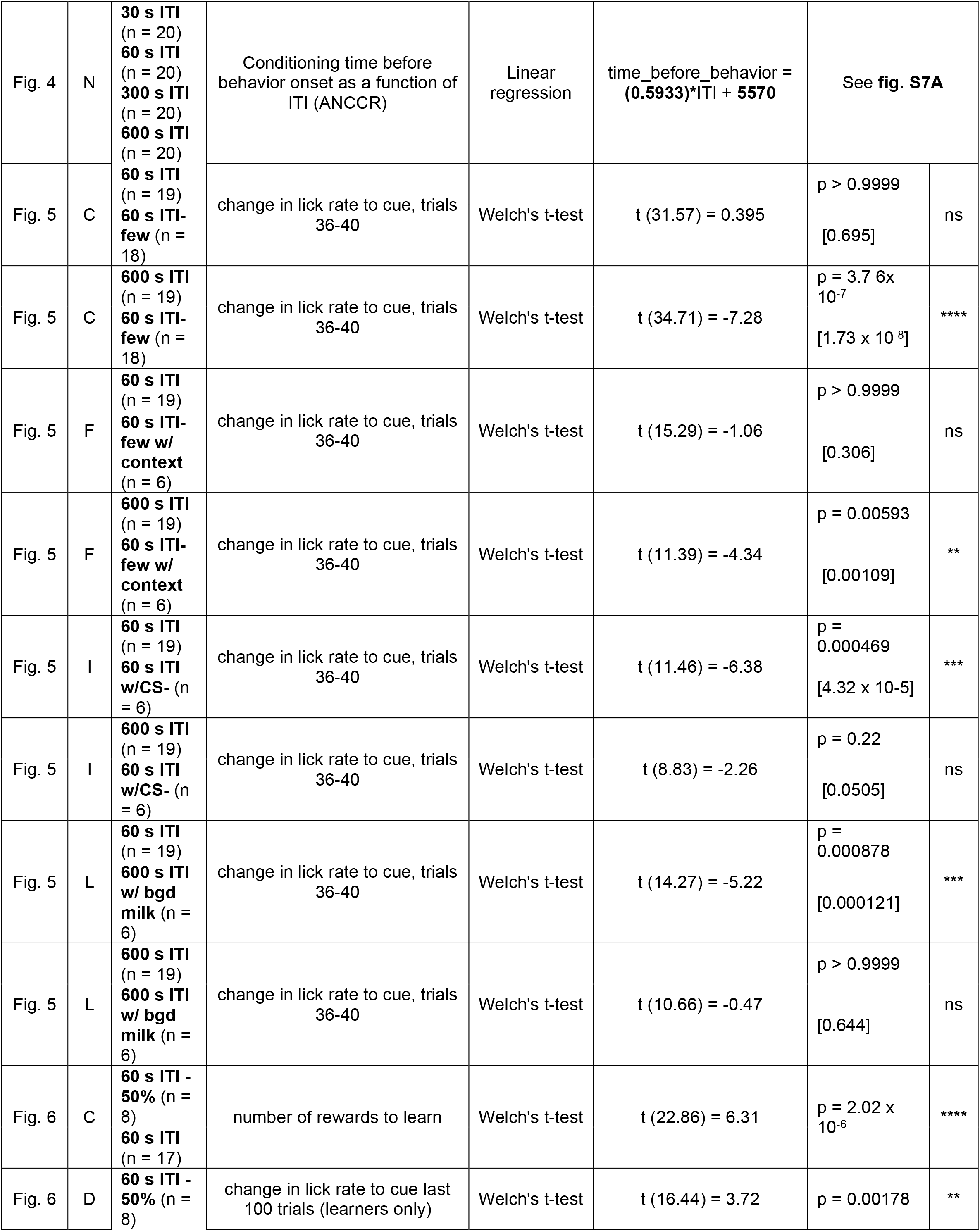
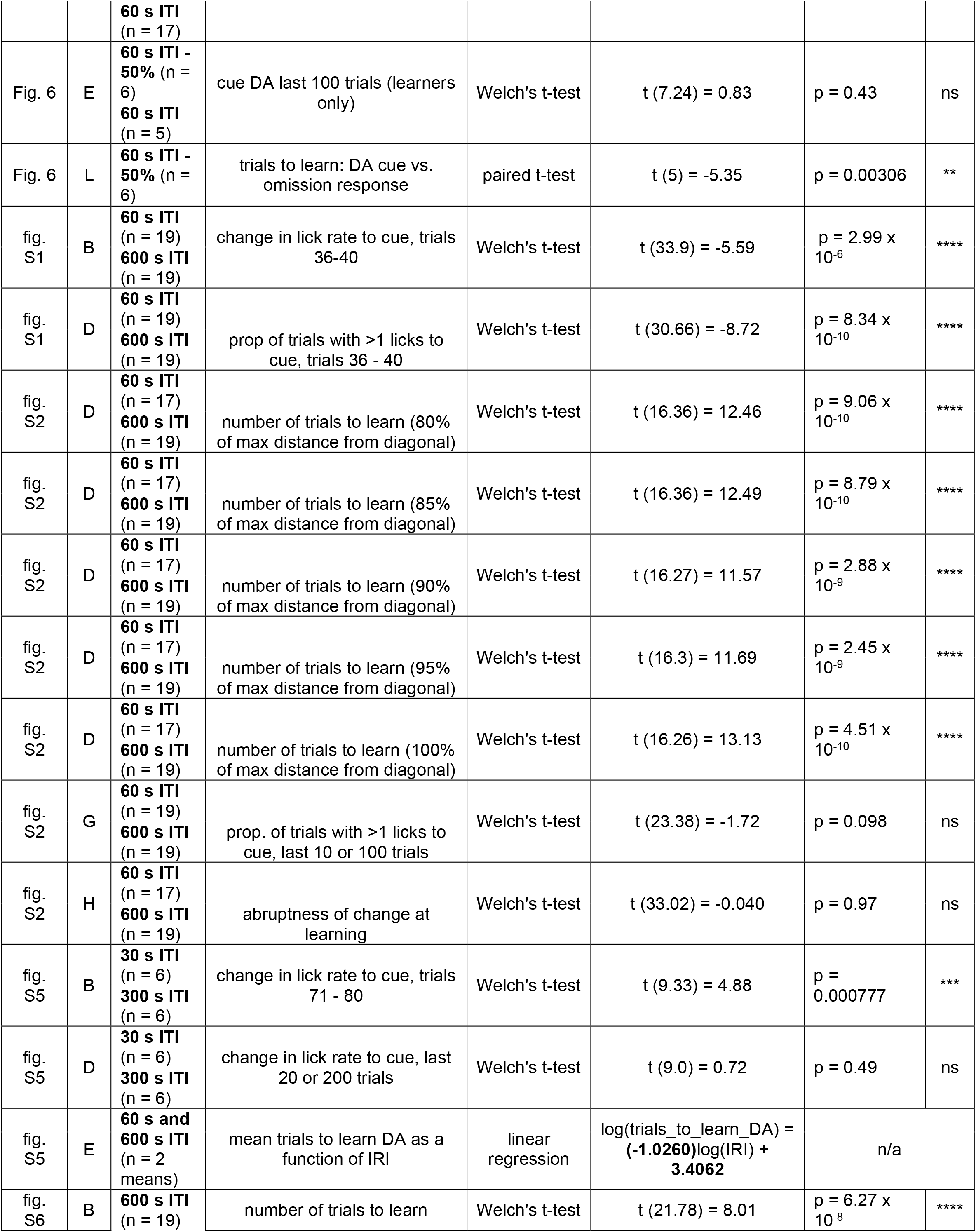
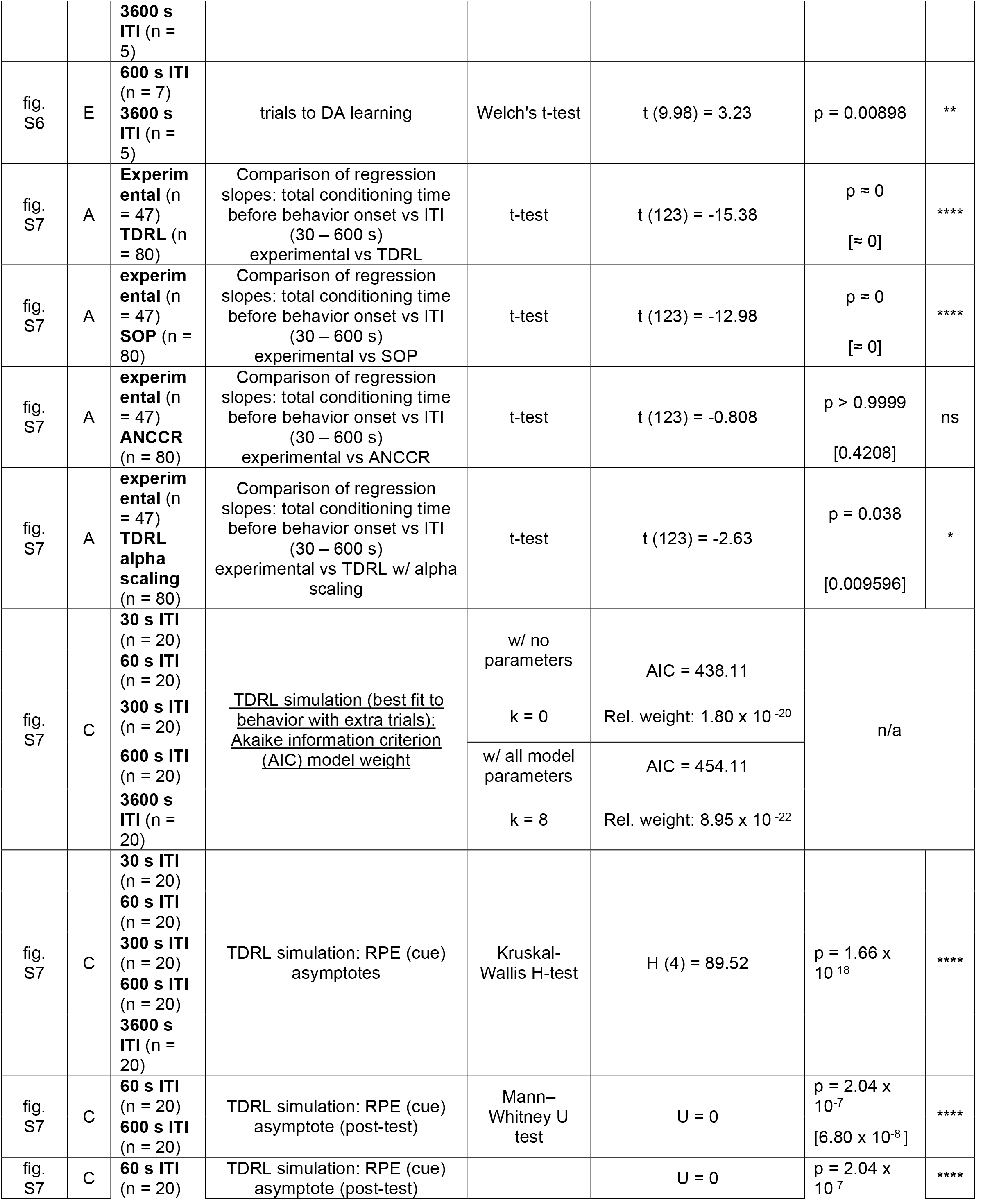
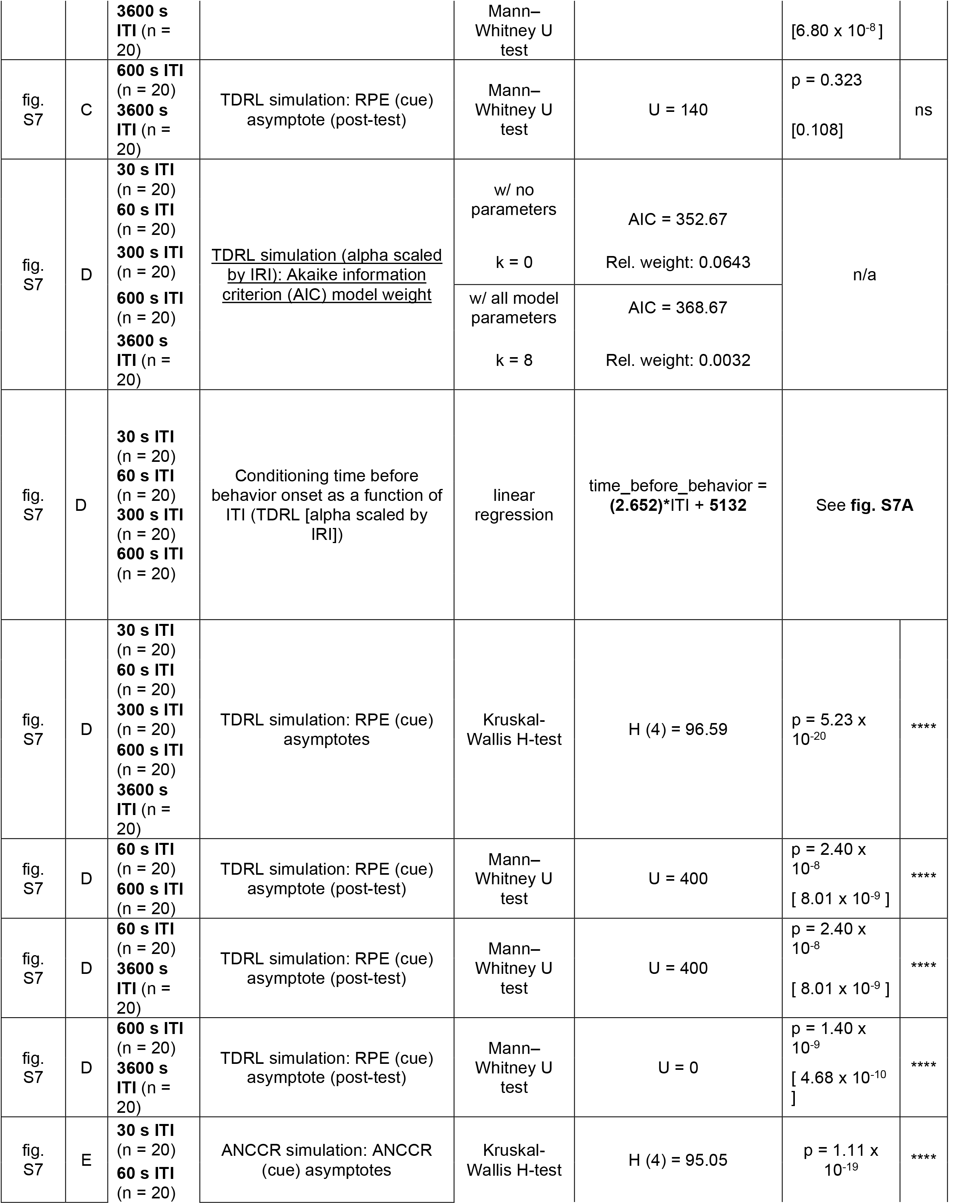
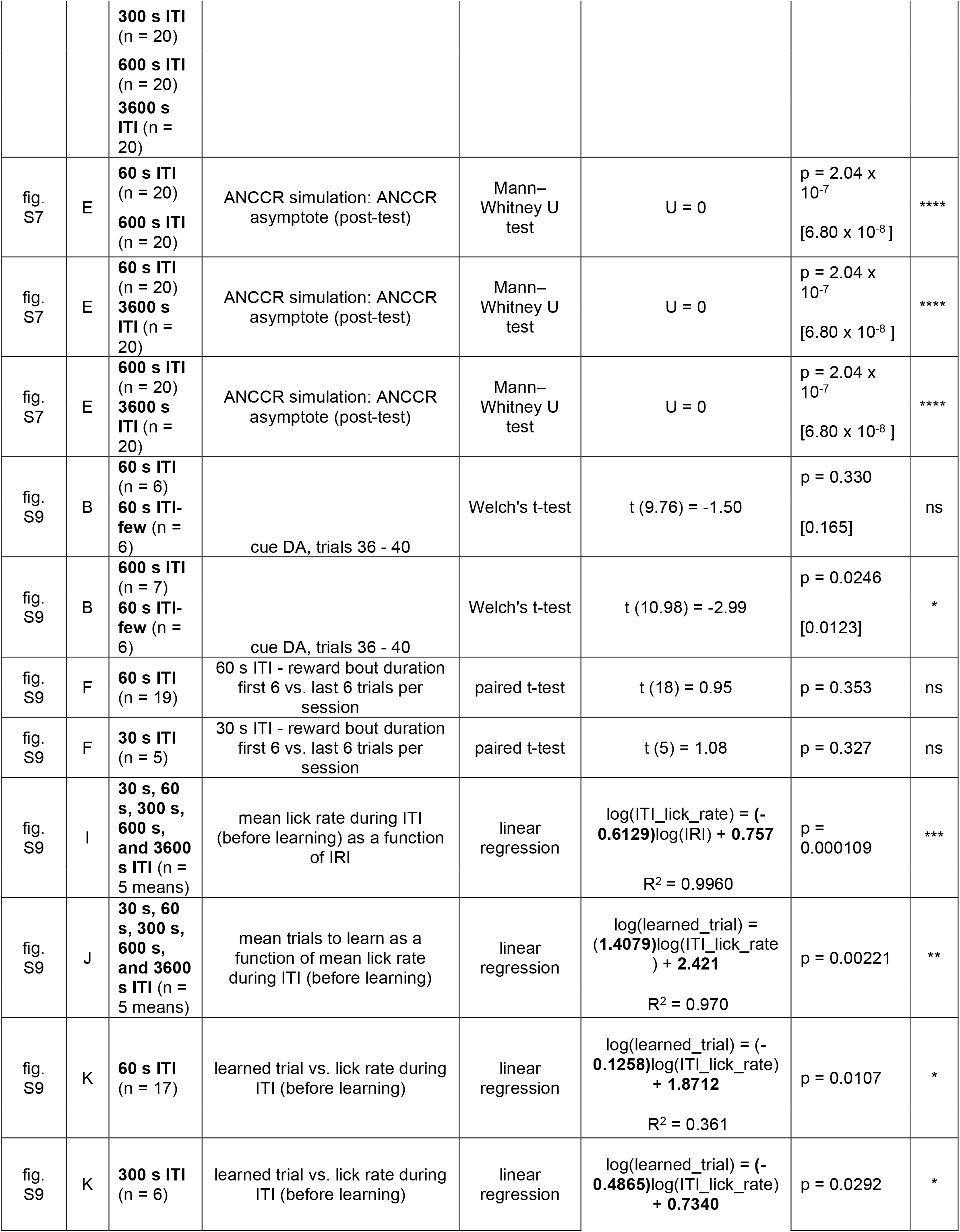
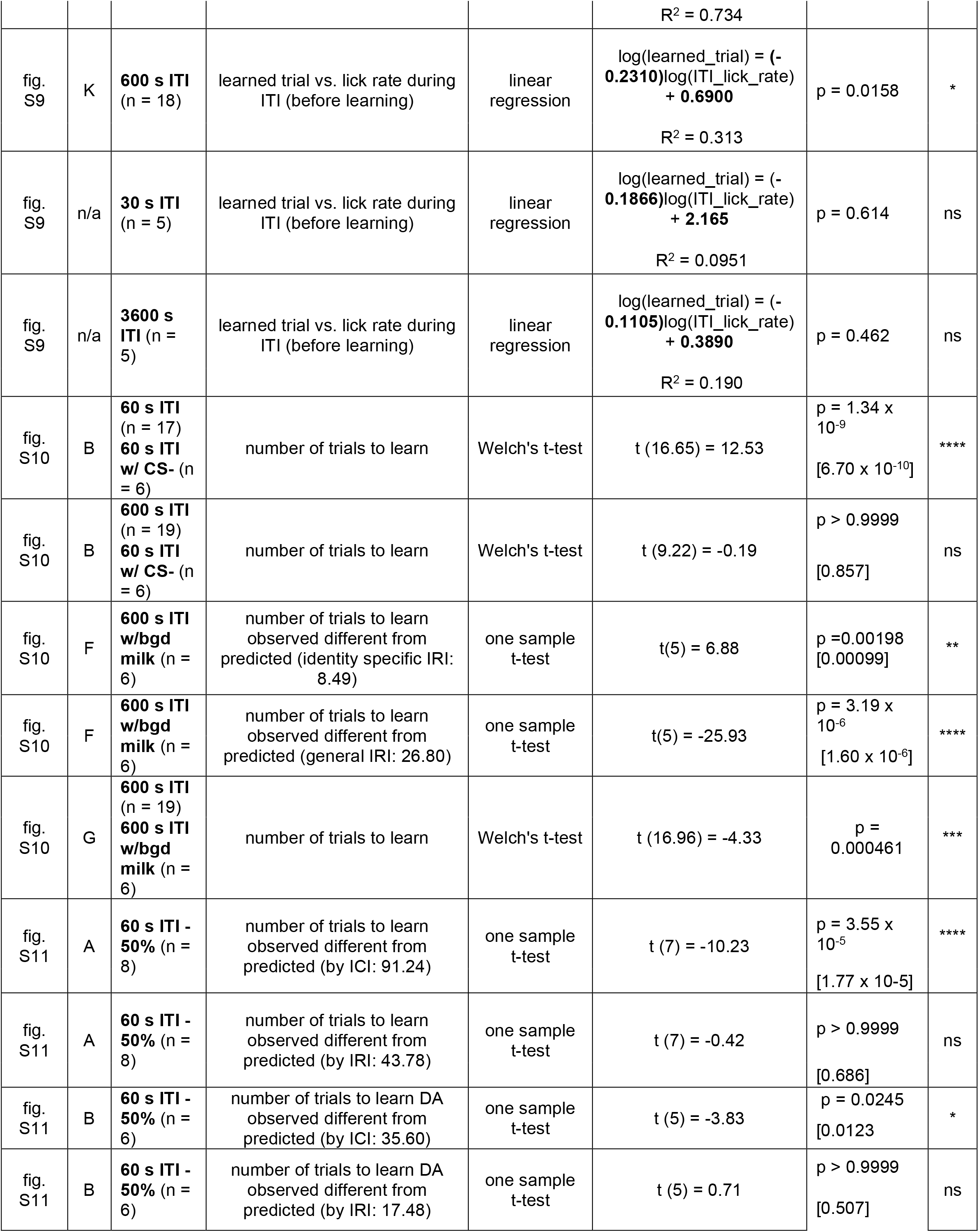
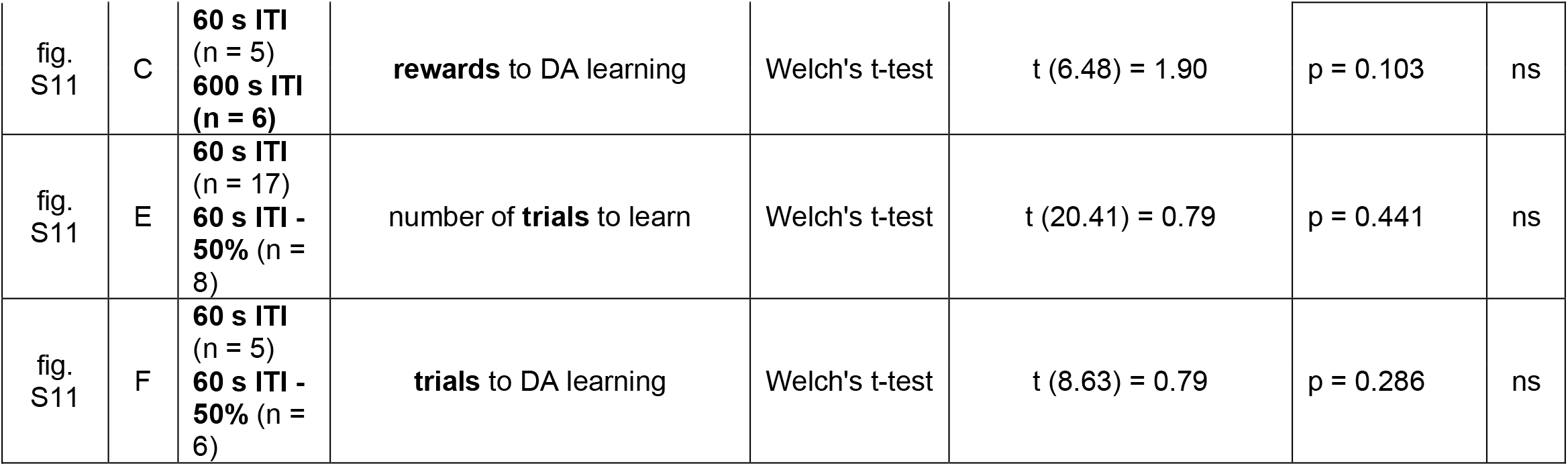

